# Does Unfairness Evoke Anger or Disgust? A Quantitative Neurofunctional Dissection Based on 25 Years of Neuroimaging

**DOI:** 10.1101/2024.10.17.618853

**Authors:** Xianyang Gan, Ran Zhang, Zihao Zheng, Lan Wang, Xi Yang, Benjamin Klugah-Brown, Ting Xu, Nan Qiu, Keith M Kendrick, Klaus Mathiak, Justin Tiwald, Dezhong Yao, Benjamin Becker

## Abstract

Over the last decades, the traditional ‘Homo economicus’ model has been increasingly challenged by convergent evidence underscoring the impact of emotions on decision-making. A classic example is the perception of unfairness operationalized in the Ultimatum Game where humans readily sacrifice personal gains to punish those who violate fairness norms. While the emotional mechanism underlying costly punishments has been widely acknowledged, the distinct contributions of moral emotions (anger or disgust) remain debated, partly due to methodological limitations of the conventional experiments. Here, we capitalize on a quantitative neurofunctional dissection approach by combining recent developments in neuroimaging meta-analyses, behavioral-level, network-level, and neurochemical-level decoding and data from 3,266 participants from functional neuroimaging studies to determine the common and distinct neural representations between unfairness and the two moral emotions. Experience of unfairness engaged a widespread bilateral network encompassing insular, cingulate, and frontal regions, with dorsal striatal regions mediating the decision to reject unfair offers. Disgust engaged a defensive-avoidance circuit encompassing amygdalar, occipital, and frontal regions, while anger engaged non-overlapping systems including mid-cingulate, thalamic, and frontal regions. Unfairness and anger or disgust respectively commonly engaged the anterior and mid-insula, while the latter additionally showed common recruitment of ventrolateral prefrontal and orbitofrontal cortices. Multimodal network, behavioral, and serotonergic decoding provided a more granular and convincing dissection of these results. Findings indicate a shared neuroaffective basis underlying the impact of emotions on unfairness-induced punishment behavior and suggest a common brain circuit has been evolutionarily shaped to protect individuals from personal harm and enforce societal norms.

## 1. Introduction

The term ‘Homo economicus’ conceptualizes humans as ‘economic agents’ who make rational choices serving their own self-interest to maximize outcome during economic decisions (Persky, 1995; A. Smith, 1776). This rationality assumption in modern economic theory has been increasingly challenged since the seminal work of Kahneman, Tversky, and colleagues stressing the impact of emotions and heuristics on decision-making (Kahneman, 2003; Kahneman, Ritov, & Schkade, 1999; Tversky & Kahneman, 1971, 1974) (see also Shafir, 2024 for an overview on their contributions in this field). These findings have spurred the development of more formalized strategies in resource allocation based paradigms from behavioral economics, with convergent evidence underscoring that emotions strongly influence decisions, such that for instance the experience of negative emotions in response to unfair distributions leads to costly punishment behavior that overrides self-interest outcome maximization (Feng, Luo, & Krueger, 2015; McAuliffe, Blake, Steinbeis, & Warneken, 2017). Such seemingly ‘irrational’ behavior has been experimentally well-documented in the Ultimatum Game (UG) paradigm. In this game theory paradigm, two players split an endowment of a limited resource between them. One player acts in the role of the ‘proposer’ and gives an offer on how to split the endowment. The other acting in the role of the ‘responder’ can next either accept the offer, in which case the endowment is split accordingly, or reject the offer, in which case neither side receives anything. Intriguingly, the responder often focuses not on maximizing self-interest, but rather punishes the proposer by ‘irrationally’ rejecting an offer that he/she considers unfair, even though such a decision is costly (i.e., at the expense of receiving nothing) (Chapman, Kim, Susskind, & Anderson, 2009; Yamagishi et al., 2009). One potential explanation is that the satisfaction of reducing the strong negative emotional experience evoked by unfair offers through punishment outweighs the actual monetary reward, implying that punishment (i.e., rejection in this case) might be more rewarding than the actual monetary outcome (Crockett et al., 2013; Crockett, Clark, Lieberman, Tabibnia, & Robbins, 2010; de Quervain et al., 2004; Gabay, Radua, Kempton, & Mehta, 2014; Hallsson, Siebner, & Hulme, 2018), thus contributing to the maintenance of fairness norms (Fehr & Fischbacher, 2004; Fehr & Gächter, 2002; Fehr & Rockenbach, 2004).

Non-human primates show a basal sensitivity to unfairness and react negatively when they receive less than a partner (Brosnan & de Waal, 2003, 2014; McAuliffe et al., 2017), suggesting an evolutionary root for strong emotional reactions to unfairness, although evidence is still lacking that these animals enforce fairness through punishment (McAuliffe et al., 2017). Animal models of the UG showed that some of our closest living relatives (i.e., chimpanzees) are rational maximizers and behave according to traditional economic models of self-interest (Jensen, Call, & Tomasello, 2007). Nevertheless, ontogenetic research on human fairness revealed that children exhibit a costly response to unfair treatment towards themselves as early as mid-childhood (Blake & McAuliffe, 2011; McAuliffe et al., 2017). Punishing those who are unfair at a personal cost may thus be considered a characteristic feature of humans (McAuliffe et al., 2017; Sheskin & Santos, 2012; Talbot, Price, & Brosnan, 2016). Indeed, fairness as well as aligning with and enforcing fairness norms represent a central part of human morality and sociality (Chapman et al., 2009; Henrich et al., 2004; Huebner, Dwyer, & Hauser, 2009; Sheskin, 2017; Tomasello, 2016). Within this context punishment of norm violations is considered to play an important role in maintaining pro-sociality (Boyd, Gintis, Bowles, & Richerson, 2003; Boyd & Richerson, 1992; Fehr & Fischbacher, 2004; Fehr & Gächter, 2002; Fehr & Rockenbach, 2004). Similar to the traditional economic conceptualization, moral psychology has also long emphasized the rationalist approach (Hallsson et al., 2018; Huebner et al., 2009; Kohlberg, 1981; Nucci, 1981; Royzman, Leeman, & Baron, 2009), however, accumulating evidence suggests that it is the negative emotional reaction (especially disgust and anger) elicited by unfair treatment in the UG that plays a key role in the costly rejection decision-making process (Figure 1a, left panel; Bargh, Schwader, Hailey, Dyer, & Boothby, 2012; Chapman & Anderson, 2013; Gan et al., 2024; Haidt, 2001; Hastie, 2001; McAuliffe et al., 2017; Oatley & Johnson-Laird, 2014; Rozin, Haidt, & Fincher, 2009; Russell & Giner-Sorolla, 2013; Vavra, Baar, & Sanfey, 2017; Xu et al., 2024; Yamagishi et al., 2009; Y. Zheng, Yang, Jin, Qi, & Liu, 2017).

**Figure 1.**
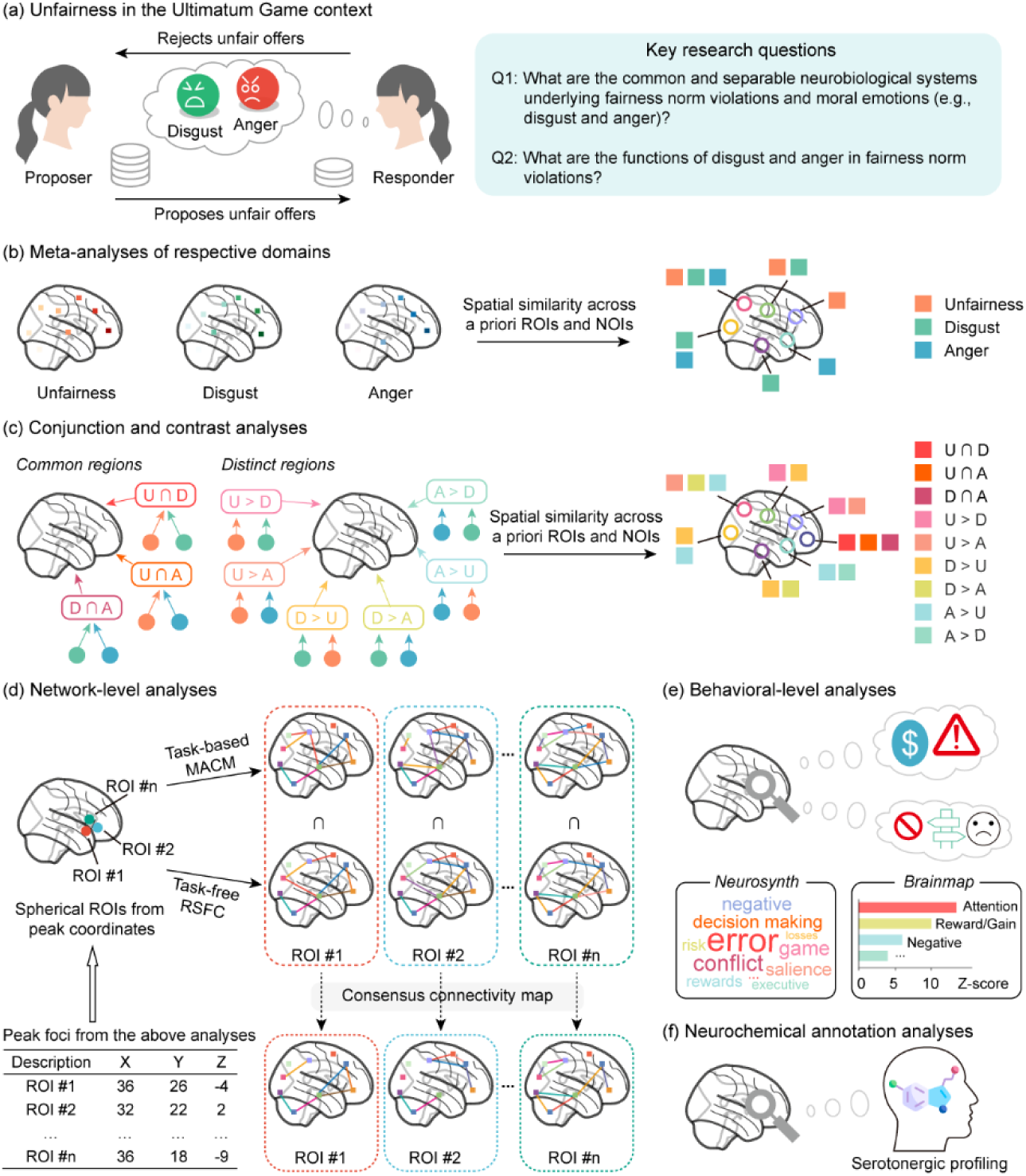
Unfairness in the UG context, key research questions, and multimodal data analytic workflow. (a) Diagram of moral emotions (e.g., disgust and anger) evoked by witnessing fairness norm violations in the UG context (left panel) and key research questions of this study (right panel). (b) Respective single-dataset meta-analysis for unfairness, disgust, and anger (left panel) and spatial similarity analyses (right panel). (c) Conjunction and contrast analyses between different domains (left panel) and spatial similarity analyses (right panel). (d) Determining the consensus connectivity profiles of each of the target ROIs based on task-based MACM and task-free RSFC. (e) Behavioral characterization analyses (i.e., functional decoding) of the identified brain regions in our meta-analyses based on the Neurosynth and Brainmap databases. (f) Hypothesis-driven neurochemical annotation analyses to examine the serotonergic profilings of the identified brain regions in our meta-analyses. Abbreviations: A = anger provocation, D = core disgust, MACM = meta-analytic connectivity modeling; NOI = network of interest, ROI = region of interest, RSFC = resting-state functional connectivity, U = unfairness.

Two emotions that involve condemning the actions of others (Haidt, 2003; Rozin, Lowery, Imada, & Haidt, 1999; Russell & Giner-Sorolla, 2013) are disgust and anger and these ‘moral emotions’ are experienced in response to a variety of moral violations (Chapman & Anderson, 2013; Giner-Sorolla, 2019; Jones, 2007; Molho, Tybur, Güler, Balliet, & Hofmann, 2017; Molho, Tybur, Van Lange, & Balliet, 2020; Olatunji & Puncochar, 2014; Prinz, 2006; Rozin & Haidt, 2013; Rozin et al., 2009; Rozin et al., 1999; Rozin, Markwith, & Stoess, 1997; Russell & Giner-Sorolla, 2013; Tybur, Lieberman, & Griskevicius, 2009; Tybur, Lieberman, Kurzban, & DeScioli, 2013). Notably, although disgust and anger may differ in several important features (e.g., physiological responses, facial expressions, action tendencies, etc.) (Chapman & Anderson, 2013; Russell & Giner-Sorolla, 2013), the fact that the two emotions are frequently triggered and experienced together in moral contexts (Chapman & Anderson, 2013; Chapman et al., 2009; Giner-Sorolla & Chapman, 2017; Giner-Sorolla, Kupfer, & Sabo, 2018; Gutierrez & Giner-Sorolla, 2007; Gutierrez, Giner-Sorolla, & Vasiljevic, 2012; Heerdink, Koning, van Doorn, & van Kleef, 2019; Russell & Giner-Sorolla, 2013; Salerno & Peter-Hagene, 2013; Simpson, Carter, Anthony, & Overton, 2006; Tybur et al., 2013; van der Eijk & Columbus, 2023) may indicate that disgust and anger share a common socioemotional core that is vital for shaping our social lives (Hutcherson & Gross, 2011).

For example, both anger and disgust are characterized by “certainty appraisals”, that is, the subjective feeling of being certain about the interpretation of the situation and what is going to happen next (C. A. Smith & Ellsworth, 1985). Tiedens & Linton found that certainty appraisals were associated with heuristic processing in decision-making, which enables people to reach decisions more quickly without systematically considering all involved factors (Tiedens & Linton, 2001). Given this, each emotion may prompt greater dependence on the other in making judgments (Olatunji & Puncochar, 2014; Salerno & Peter-Hagene, 2013), and the co-occurrence of the two emotions provides the motivational force for people to live up to and enforce punishment actions against moral violators (Molho et al., 2017; Molho et al., 2020; Olatunji & Puncochar, 2014; Skitka, Bauman, & Mullen, 2008). In the present study, we capitalize on the vast data provided by a large body of original brain studies that examined the neural basis of unfairness, disgust, and anger processing, to determine the underlying neural mechanism of the often intertwined disgust and anger responses to moral norm violations (as operationalized in the UG) given that fairness norm transgressions provide a prototypical instance to explore such questions (McCall & Singer, 2015; Rozin et al., 2009; Russell & Giner-Sorolla, 2013).

Numerous empirical studies from behavioral, electrophysiological, and neuroimaging perspectives highlight the important role of disgust and anger in fairness-related moral judgment and decision-making. On the behavioral level, for example, several studies reported that small offers that are perceived as unfair in the UG elicit anger and disgust which in turn impacts the decision to reject an offer (Chapman et al., 2009; Pillutla & Murnighan, 1996; Srivastava, Espinoza, & Fedorikhin, 2009). In addition to evidence on the level of subjective experiences, assessment of the physiological and expressive correlates of the emotional reaction (i.e., electromyography, EMG) indicate that facial disgust expressions increased with increasing inequality of offers, which moreover predicted increasing rejection probabilities of unfair offers (Chapman et al., 2009) (for the generalization of facial disgust expressions to fairness violations beyond the UG context, see also Cannon, Schnall, & White, 2011; and Danovitch & Bloom, 2009). The results from both, subjective and objective measurements thus indicate that negative emotions induced in response to unfairness – in particular disgust and anger - strongly motivate subsequent punishment behavior (i.e., irrationally rejecting an unfair offer) (see also Heffner, Son, & FeldmanHall, 2021; Xu et al., 2024). Further support for the importance of anger and disgust in explaining behavior in the UG comes from emotion induction studies showing that both priming disgust and priming anger lead to perceiving unequal offers as more unfair and further increased rejection of unfair offers (Andrade & Ariely, 2009; Harlé & Sanfey, 2010; Moretti & di Pellegrino, 2010). While other researchers manipulated the time between the offer and subsequent acceptance decisions in the UG found that a time lag could effectively reduce (or “cool off”) the negative emotions (e.g., anger) resulting from receiving an unfair offer, which in turn led to decreased costly rejection rates (Grimm & Mengel, 2011; Oechssler, Roider, & Schmitz, 2015; Sutter, Kocher, & Strauß, 2003; Wang et al., 2011). Notably, when the responders were instructed to engage in emotional rumination rather than a distractor task during the delay the rejections increased, which was driven by the level of anger experience (Wang et al., 2011). Relatedly, acceptance rates of unfair offers significantly increased when the responders were given the opportunity to express their negative feelings (e.g., anger) by sending a written message to the proposer, suggesting that the rejection of unfair treatments might be used to express negative emotions triggered by unfair offers (Xiao & Houser, 2005). In line with this interpretation, Yamagishi et al. compared rejection rates of three UG variants and argued that the rejection of unfair offers was an “emotional commitment device” based on emotions like disgust or anger (Yamagishi et al., 2009). While behavioral evidence from studies using both forward and reverse strategies (e.g., examining whether moral transgressions elicit disgust experience, or whether disgust induction influences moral cognition, respectively) converges on supporting the notion that negative emotions like disgust and anger play a role in unfairness-related moral transgressions, the specific contributions remain controversial given the common and often inherent co-occurrence of the emotions. Based on the neurobiological evidence outlined below, we therefore propose a meta-analytic neurobiological approach that may allow to contribute to disentangling the contributions of the emotional domains.

Electrophysiological studies showed that unfair offers evoked negative emotions as well as larger feedback-related negativity (FRN) than fair offers in the UG, and the resulting FRN could further predict the rejection of unfair offers (Hewig et al., 2011) (for similar findings see Van der Veen & Sahibdin, 2011). Moreover, the amplitudes of the FRN increased as a function of experienced anger (Riepl, Mussel, Osinsky, & Hewig, 2016). Another electrophysiological study employed a dipole localization technique and found that unfair offers induced activation of the insula reflecting the induction of negative emotions (e.g., anger) evoked by unfair treatment with higher insula activity predicting a subsequent rejection (Güçlü, Ertaç, Hortaçsu, & List, 2012). Evidence from neuroimaging research echoes the findings above. For example, Sanfey et al. observed increased activation of the bilateral anterior insula (AI) elicited by unfair offers in comparison to fair offers and interpreted these results as emotional responses (i.e., disgust, anger) to the proposer’s unfair behavior. Importantly, the level of AI activation was positively correlated with rejection rates of unfair offers (Sanfey, Rilling, Aronson, Nystrom, & Cohen, 2003). Although the insula is a functionally heterogeneous region (Ferraro et al., 2022; Gan et al., 2022), an abundance of evidence suggests that its engagement in response to moral violations may reflect a negative emotional state (especially disgust and anger) in response to norm violations (e.g., unfair treatment) (Bellucci, Feng, Camilleri, Eickhoff, & Krueger, 2018; Feng et al., 2015; Gabay et al., 2014; Li et al., 2024; McAuliffe et al., 2017; Vicario, Rafal, Martino, & Avenanti, 2017; Yang, Zheng, Yang, Li, & Liu, 2019; Zinchenko & Arsalidou, 2018). In addition to evidence from the conventional functional magnetic resonance imaging (fMRI) approach, more fine-grained analyses using multivariate pattern decoding (Poldrack, 2006) on both regional (e.g., insula) and whole-brain levels confirmed shared neural representations of unfairness and disgust (as well as potentially anger) (Corradi-Dell’Acqua, Tusche, Vuilleumier, & Singer, 2016; Gan et al., 2024).

Additional behavioral and neurobiological support for the hypothesis that disgust and anger play a key role in fairness-related norm transgressions comes from research on psychiatric populations as well as research on the underlying neurotransmitters using the UG paradigm. Individuals with depression report greater levels of disgust and anger than controls in response to unfair offers (Harlé, Allen, & Sanfey, 2010). Patients with anxiety respond to the proposer’s unfair offers with less anger, rate these less unequal, and accept more unfair offers compared with controls (Grecucci et al., 2013). Furthermore, patients with bipolar disorder react with higher levels of anger to unfair offers and reject a higher rate of moderately unfair offers than controls (Duek, Osher, Belmaker, Bersudsky, & Kofman, 2014; similar findings see also Lois, Schneider, Kaurin, & Wessa, 2020). Evidence from studies focusing on the neurotransmitter basis of behavior in the UG additionally supports an association with emotional experiences. For example, serotonin (5-HT) has been found to influence decision-making in the UG. Transiently decreasing serotonin levels through dietary acute tryptophan depletion increased rejection rates of unfair offers (Crockett et al., 2013; Crockett, Clark, Lieberman, et al., 2010; Crockett, Clark, Tabibnia, Lieberman, & Robbins, 2008). The effect seems to be mediated by the striatal response to punishment, such that depletion increased dorsal striatal responses to punishing an unfair proposer (Crockett et al., 2013). However, increasing serotonin levels by selective serotonin reuptake inhibitors led to a less unlikely rejection of unfair offers (Crockett, Clark, Hauser, & Robbins, 2010). These results indicated that emotional reactions indiced by variations in serotonergic signaling may indicate variations in the perceived unfairness and may play a key role in punishment behavior (Crockett, Clark, Lieberman, et al., 2010). Along similar lines, a shared serotonergic system for core disgust and negative evaluations of moral violations (also term ‘moral disgust’; Chapman & Anderson, 2013; Chapman et al., 2009; Rozin & Haidt, 2013; Tybur et al., 2013) was verified based on evidence from abnormal disgust processing in clinical disorders characterized by dysregulated serotonergic transmission (Vicario et al., 2017).

Summarizing, multiple lines of evidence reviewed above suggest that fairness-related moral violations can reliably elicit negative emotions such as disgust and anger. Evaluation (i.e., appraisal) theories of emotion have been adopted to formalize the relationship between negative emotions (e.g., disgust, anger) and fairness norm violations (Chapman & Anderson, 2013; Gan et al., 2024; Mullen, 2007; Rozin et al., 2009). According to this perspective, emotions are not elicited directly by the stimulus event itself, but by the way an individual evaluates or appraises the significance of a stimulus or event (i.e., whether it is relevant to his or her current goals, concerns, and resources) (Chapman & Anderson, 2013; Delplanque & Sander, 2021; Oatley & Johnson-Laird, 2014; Pool & Sander, 2021; Scherer, 2004, 2005). Such events may include immoral acts committed by other individuals, such as unequal monetary offers, as well as provocative anger-inducing behavior (e.g., loud noise inflicted by others) or disgust-inducing stimuli that may pose harm. Within this context, the appraisal of an unfair event will elicit negative emotions such as anger and disgust, which should mirror the neural response directly induced by anger provocation and core disgust (for initial neurobiological evidence see also Gan et al., 2024). As such, capitalizing on this framework and the underlying neural processes allows to test shared appraisal-related brain processes and in turn to determine the extent to which unfairness-related moral appraisals are separable from the appraisals for core disgust and anger provocation (Chapman & Anderson, 2013).

Although behavioral research provides us with important evidence for the co-occurrence of disgust and anger in appraising unfair treatment, the majority of these studies rely on subjective measures (i.e., verbal self-reports), which makes it impossible to separate disgust from anger (Giner-Sorolla, 2019; Olatunji & Puncochar, 2014). Previous behavioral studies employed objective EMG recording in this field, however, the specific facial measurements for disgust and anger were often confounded with each other (Chapman & Anderson, 2013; Chapman et al., 2009; Rozin et al., 2009; Russell & Giner-Sorolla, 2013), making it impossible to disentangle the specific contributions of both emotions. Despite the high temporal resolution of electrophysiological measurements (Gan et al., 2020), MRI-based functional neuroimaging with a higher spatial resolution and in combination with advanced analytic approaches such as neural decoding or meta-analytic modeling may have the potential to determine brain activation patterns that are distinct and shared for emotional and affective processes (see e.g., Čeko, Kragel, Woo, López-Solà, & Wager, 2022; Costa et al., 2024; Feng, Eickhoff, et al., 2021; Gan et al., 2022; Liu et al., 2024; Putkinen et al., 2021; Woo, Chang, Lindquist, & Wager, 2017; Zhang et al., 2024; Zhou et al., 2023). However, the common and distinct neurobiological basis of unfairness, disgust, and anger has not been systematically determined. The neuroimaging-based meta-analytic approach could realize such aims as it allows to determine convergent and robust neurofunctional basis of emotional and cognitive processes as well as to determine common and distinct neurofunctional representations of different mental processes (Chen et al., 2018; Costa et al., 2024; Feng et al., 2018; Feng, Eickhoff, et al., 2021; Feng, Gu, et al., 2021; Ferraro et al., 2022; Gan et al., 2022).

By capitalizing on pre-registered activation likelihood estimation (ALE) neuroimaging meta-analyses, we aim to determine brain systems that are robustly engaged in unfairness, disgust, and anger, respectively, in healthy individuals (for an outline of key research questions and approaches see also Figure 1a, right panel and Figure 1b). Specifically, we here aim for the first time to systematically determine common and separable brain systems involved in unfairness and disgust as well as in unfairness and anger (Figure 1c), which – in a conceptual context – may enable us to assess the neural plausibility of the appraisal models outlined above. To further improve the interpretation of the findings the core meta-analyses will be flanked by multimodal data analytic techniques (i.e., network-level, behavioral-level, and neurochemical level) to provide comprehensive insights and guide interpretation of the findings (Figure 1d-f). Finally, we will perform a series of exploratory meta-analyses aiming at further determining common and separable brain systems engaged in unfairness-related decision-making (rejection response) and disgust as well as anger.

## 2. Methods

### 2.1. Literature search

The current meta-analytic review was performed in line with PRISMA guidelines for systematic reviews (Page et al., 2021) and the meta-analytic protocols were preregistered on the Open Science Framework (https://osf.io/fpc8v). Unfairness related, core disgust related and anger provocation related original studies published prior to Feb 28, 2023, were identified using the following search terms on the PubMed and Web of Science databases. (1) For Unfairness related studies: (“Ultimatum Game” OR “unfairness” OR “unfair”) AND (“fMRI” OR “functional magnetic resonance imaging”). (2) For core disgust related studies: (“disgust” OR “disgust elicitors” OR “disgusting elicitors” OR “disgust stimuli” OR “disgusting stimuli”) AND (“fMRI” OR “functional magnetic resonance imaging”). Of note, different from a recent meta-analysis that focused only on the visual modality (Gan et al., 2022), we here included studies that employed core disgust stimuli not only from the visual modality but also from other modalities such as gustatory, olfactory, etc. (3) For anger related studies we specifically focused on anger provocation related studies (e.g., Taylor aggression paradigm, Interpersonal provocation procedure, Anger-induction film clips, etc.) given that anger provocation has been demonstrated to induce robust subjective anger experience (Carver & Harmon-Jones, 2009; Denson, Pedersen, Ronquillo, & Nandy, 2009; Kjærvik & Bushman, 2024; Klaus & Schutter, 2021; Sorella, Grecucci, Piretti, & Job, 2021). Noteworthy, we will not include studies related to Cyberball Paradigm as its role in inducing anger is debated and likely rather unspecific. Based on this operational definition, the search terms for anger provocation related studies are: (“anger” OR “angry” OR “aggression” OR “provocation”) AND (“fMRI” OR “functional magnetic resonance imaging”). Moreover, reference lists of included articles were also searched for relevant articles not identified during the initial database search.

### 2.2. Inclusion and exclusion criteria

To identify suitable publications according to our criteria, two independent reviewers (X. G. and R. Z.) initially examined the titles and abstracts of identified papers. Full texts of eligible papers were downloaded for detailed examination. Any disagreement between the two reviewers was resolved by discussion and a final decision by a third independent reviewer (Z. Z.). Papers that passed the screening process were thoroughly evaluated for eligibility (B. B. and X. G.), studies with ambiguous experimental contrasts were removed at this stage and disagreements were further resolved by another reviewer (X. Y.). A flowchart illustrating the detailed article screening, exclusion, and inclusion process is shown in Supplemental Materials Figure S1a,b. All included studies irrespective of domains should meet the following inclusion criteria: (1) original paper published in a peer-reviewed journal in English; (2) single group data from healthy subjects reported (i.e., healthy volunteers only); (3) task-related fMRI employed; (4) results reported in standard stereotactic coordinates (either Talairach or Montreal Neurological Institute [MNI] space); (5) coordinates relative to psychophysiological or psychopathological correlations were excluded; (6) univariate (i.e., general-linear-model-based) whole-brain results reported (we exclude studies reporting only ROI analyses); (7) studies recruiting drug-free subjects. While the tailored inclusion criteria for each domain were as follows. For UG related literature: (1) specific contrasts reporting unfair > fair or reject > accept conditions or relevant continuous parametric analyses; (2) reported data from participants acted as responders rather than proposers in the UG. Of note, in line with the main aim of the present study, the unfairness related meta-analysis focused on neural reactivity in response to Unfairness (unfair > fair) as captured in the standard version equivalent of the UG.

The additional meta-analysis on neural activity related to the behavioral Response (reject > accept) represents an exploratory further analysis in the current project. For core disgust related literature: specific contrasts reporting disgust > baseline condition (e.g., disgust > neutral) or relevant continuous parametric analyses. For anger provocation related literature: specific contrasts reporting anger > baseline, high provocation > low provocation conditions, or relevant continuous parametric analyses. Also, we contacted corresponding authors of 16 studies where Talairach or MNI coordinates for the basic contrasts were not reported to reduce the possibility of a biased sample set.

After the in-depth screening procedure, a total of 108 papers (Supplemental Materials Table S1) were included in the meta-analyses, and according to the aims of the analyses were organized as follows: (1) unfairness: 28 papers, 28 experiments, 280 foci, 1090 subjects; (2) core disgust: 43 papers, 45 experiments, 488 foci, 949 subjects; (3) anger provocation: 24 papers, 24 experiments, 276 foci, 785 subjects; (4) rejection: 13 papers, 14 experiments, 128 foci, 442 subjects. Based on recommendations for ALE meta-analyses suggesting that at least 20 (and 17 in the worst case) experiments are sufficient to determine moderate effects with sufficient statistical power (80%) (Eickhoff et al., 2016), the first three separate meta-analyses allowed us to draw reliable and stable inferences, while for the exploratory meta-analysis on rejection the comparably low number of experiments warrants a cautious evaluation of the robustness of the findings.

### 2.3. Activation likelihood estimation

We employed the current version of the ALE algorithm (GingerALE v.3.0.2, available via http://www.brainmap.org/ale/) to conduct a series of coordinate-based meta-analyses (Eickhoff, Bzdok, Laird, Kurth, & Fox, 2012; Eickhoff et al., 2011; Eickhoff et al., 2009). Coordinates and sample sizes were manually extracted from the original articles, and the Talairach coordinates were converted to MNI space using the Lancaster transform algorithm implemented in GingerALE (Laird et al., 2010; Lancaster et al., 2007). Briefly, the ALE method computes the probability of each voxel being activated under multiple independent experiments with consideration of participants in each experiment. In other words, ALE aims to determine whether convergent activation foci across a number of included experiments occur at a level greater than chance (Laird et al., 2005). Specifically, ALE treats the reported peak foci from single studies as three-dimensional Gaussian probability distributions and provides empirical estimates to address the spatial uncertainty which may be associated with the between-subject and between-template variability of the neuroimaging foci (Eickhoff et al., 2009). For each Gaussian probability distribution function, the full width at half maxima (FWHM) is weighted by the number of subjects included in the experiment, such that experiments with larger sample sizes will have smaller Gaussian distributions, thus yielding more reliable estimates of “true” activation for larger sample sizes (Eickhoff et al., 2009).

Modeled activation maps for each included study were created by combining the probabilities of all activation foci for each voxel (Eickhoff et al., 2009; Turkeltaub et al., 2012). The union of individual modeled activation maps across experiments was calculated to produce an ALE map that reflected areas of the brain (clusters) corresponding to the location of convergence (Eickhoff et al., 2012; Eickhoff et al., 2009). In line with recommendations, for the single dataset analysis all ALE maps were thresholded using a cluster-level familywise error (cluster-FWE) correction at p < 0.05, with a voxel-level threshold of p < 0.001 (uncorrected cluster forming) (Eickhoff, Laird, Fox, Lancaster, & Fox, 2017; Eickhoff et al., 2016; Ferraro et al., 2022; Gan et al., 2022; Klugah-Brown et al., 2020; Klugah-Brown et al., 2021; Müller et al., 2018). The p value accounts for the proportion of the null distribution of random spatial relation between the various experiments. In our analyses, p values were generated by 5,000 permutations (Gan et al., 2022; Laird et al., 2010; Tao, He, Lin, Liu, & Tao, 2021).

### 2.4. Contrast and conjunction analyses

Contrast analyses compare and contrast two ALE datasets that are generated from previous single dataset analyses (Eickhoff et al., 2012). When performing contrast analyses, GingerALE randomly divides the pooled foci datasets into two new datasets of the same size as the original individual computed datasets. Next, voxel-wise differences between the two new datasets are determined through a subtracting procedure: an ALE image is created for each new dataset, then subtracted from the other, and finally compared with the true data. After many permutations, this generates a voxel-wise p value image showing where the true data’s values locate on the distribution of values in that voxel. Different from the statistical procedure used for contrast analyses, conjunction is conducted using voxel-wise minimum value of the input ALE images to create the output image (Nichols, Brett, Andersson, Wager, & Poline, 2005), which is equivalent to determination of the overlap between two ALE maps. The resulting results represent shared neural correlates between two tasks/domains (Eickhoff et al., 2011).

In the current study, a series of contrast and conjunction analyses were performed to identify separable (e.g., unfairness > core disgust, unfairness > anger provocation) and shared (e.g., unfairness ∩ core disgust, unfairness ∩ anger provocation; where ∩ indicates conjunction) neural substrates between two domains. Finally, a conjunction across the networks of unfairness, core disgust and anger provocation was conducted to identify neural correlates commonly recruited by all three domains.

Consistent with previous studies, for the contrast analysis the ALE images were thresholded at p < 0.05 (Alain, Du, Bernstein, Barten, & Banai, 2018; Gan et al., 2022; Papitto, Friederici, & Zaccarella, 2020), with minimum cluster volume of 200 mm^3^ and 10,000 permutations (Gan et al., 2022; Qiu et al., 2022). Given that the conjunction analysis directly extracts ALE scores from the ALE images obtained from single dataset analysis, the threshold was identical to the one for single dataset analysis.

### 2.5. Spatial similarity between respective maps and selected brain systems

To further separate common and specific neural contributions on the individual level, we employed a sensitive approach that illustrates spatial similarity between respective maps derived from single dataset analyses as well as contrast and conjunction analyses using riverplots for a priori ROIs previously documented as regions associated with unfairness, core disgust, and anger provocation (Corradi-Dell’Acqua et al., 2016; Denson et al., 2009; Feng et al., 2015; Gabay et al., 2014; Gan et al., 2024; Gan et al., 2022; Sorella et al., 2021). We moreover calculated spatial similarity between respective maps and seven large-scale cerebral networks (Yeo et al., 2011). In line with recent studies (Čeko et al., 2022; Gan et al., 2024), spatial similarity was computed as cosine similarity between each ROI or network and corresponding map obtained from single dataset analyses as well as contrast and conjunction analyses.

### 2.6. Task-based connectivity: MACM analyses

To investigate the meta-analytic co-activation patterns of each insula region identified in contrast and conjunction analyses (details see the Results section), MACM analyses were employed using the peak coordinates of each insula region derived from contrast and conjunction analyses as ROIs (with sphere radius = 6 mm) based on the BrainMap database. Note that if one cluster encompassed two insula-related peak foci, we will focus on the peak with the higher ALE/Z value. Briefly, MACM examines whole-brain functional connectivity profiles by identifying the above-chance covariance between two or more brain regions (Laird et al., 2009; Robinson et al., 2012; Robinson, Laird, Glahn, Lovallo, & Fox, 2010). This approach is similar to the resting-state functional connectivity patterns (Kotkowski, Price, Mickle Fox, Vanasse, & Fox, 2018; Reid et al., 2017; S. M. Smith et al., 2009), but further complements the resting-state functional connectivity approach as it provides a measure of functional connectivity during a range of task-based mental states (Langner, Rottschy, Laird, Fox, & Eickhoff, 2014).

We conducted a series of MACM analyses using the BrainMap’s Sleuth v.3.0.4 software (https://brainmap.org/sleuth/). Consistent with the recommendations in the user manual for GingerALE (https://www.brainmap.org/ale/manual.pdf) and several prior studies (Gan et al., 2022; Meier, Ray, Mastan, Salvage, & Robin, 2021; Papitto et al., 2020), our search terms in the Sleuth were as follows: (1) “Locations: MNI Image” used to upload the corresponding spherical ROI in MNI space, (2) “Experiments: Activations, Activations only”, (3) “Experiments: Context, Normal Mapping”, (4) “Subjects: Diagnosis, Normals”. This allowed us to identify brain region covariance above-chance within a given seed ROI across a large and different pool of neuroimaging experiments from the functional database. After each respective ROI search, coordinates from experiments meeting the criteria were converted into MNI space and downloaded as a text file. The MACM analyses were based on the twelve datasets for each of the ROIs (see Table 1 for further information). To determine areas of convergence of co-activation with each ROI, these new datasets were analyzed separately using GingerALE v.3.0.2 with the same statistical criteria previously applied in the single dataset analyses (i.e., voxel-level of p < 0.001 (uncorrected), cluster-level of p < 0.05 corrected, 5,000 permutations).

**Table 1.** The composition of the twelve datasets used for MACM analyses.

| Condition | Insula | Number of experiments | Number of foci | Number of subjects |
| --- | --- | --- | --- | --- |
| Unfairness ∩ Core disgust | Right AI | 288 | 4858 | 4841 |
|  | Left AI | 358 | 6329 | 6227 |
| Unfairness ∩ Anger provocation | Right AI | 311 | 5405 | 6355 |
|  | Left AI | 317 | 5861 | 5163 |
| Core disgust ∩ Anger provocation | Left MI | 111 | 1929 | 1660 |
| Unfairness > Core disgust | Left AI | 97 | 1408 | 1790 |
|  | Right AI | 184 | 3194 | 2873 |
| Unfairness > Anger provocation | Right AI | 137 | 2229 | 2133 |
|  | Left AI | 159 | 2385 | 2451 |
| Anger provocation > Unfairness | Left AI | 136 | 1942 | 1979 |
| Anger provocation > Core disgust | Left AI | 89 | 1644 | 1313 |
|  | Right AI | 55 | 1146 | 693 |

Abbreviation: MI = middle insula

### 2.7. Task-free connectivity: RSFC analyses

To examine the functional connectivity profiles of each insula region identified from contrast and conjunction analyses (details see the Results section), RSFC analyses were employed on original data using the peak coordinates of each insula derived from contrast and conjunction analyses as seed ROIs (with sphere radius = 6 mm). Consistent with the MACM analyses, if one cluster reports two insula-related peak foci, the peak with the higher ALE/Z value will be selected. The data used for the RSFC analyses was from an original independent study. Specifically, resting-state fMRI data of 100 healthy subjects were collected from the Open Access Series of Imaging Studies (OASIS)-3 dataset (https://central.xnat.org). Images were acquired on a Siemens TimTrio 3T scanner. Resting-state functional images were collected using an echo-planner imaging sequence with the following parameters: repetition time = 2200 ms, echo time = 27 ms, flip angle = 90°, voxel size = 4.0 × 4.0 × 4.0 mm^3^, number of slices = 33 and slice thickness = 4 mm. Preprocessing was performed using Data Processing Assistant for Resting-State fMRI (DPARSF, v.5.0_201001) (http://rfmri.org/DPARSF). Briefly, functional imaging preprocessing procedures consisted of the following steps: (1) deletion of the initial 5 volumes to improve stability of the magnetic field, (2) slice-timing correction for differences in slice acquisition timing, (3) realignment of head motion, (4) normalization of functional images to the standard MNI space, (5) grey matter, white matter, cerebrospinal fluid, and head motion regression, (6) detrending, (7) filtering (0.01 - 0.1 Hz), (8) and spatial smoothing with a 6-mm FWHM Gaussian kernel.

Using the DPARSF toolbox, the processed time-course of each seed ROI was then correlated with the time-series of all other voxels in the brain. The voxel-level correlation coefficients were transformed into Fisher’s z-score and tested for consistency across participants (thresholded at p < 0.05 cluster-level FWE correction, with cluster-forming voxel-level threshold at p < 0.001; similar approach see Bellucci et al., 2018).

### 2.8. Consensus connectivity map

After obtaining brain regions showing task-based co-activation (i.e., MACM) and task-free functional connectivity (i.e., RSFC), conjunction analyses were conducted across the MACM and RSFC maps for each seed ROI using the minimum statistic approach (Nichols et al., 2005). The resultant consensus connectivity maps contained brain areas showing consistent interactions with each seed region across task-driven and task-independent functional networks (Bellucci, Camilleri, Iyengar, Eickhoff, & Krueger, 2020; Bore et al., 2024). Moreover, these connectivity profiles allowed us to capitalize on different connectivity profiles of insula subregions (Cauda et al., 2012; Klugah-Brown et al., 2023) and as such interpret the results in a more nuanced way although the majority of regions were identified corresponding to the anterior portion of the insula (AI) in the atlas. In accordance with the approach used in previous studies (Bellucci et al., 2020; R. Gu et al., 2019), if bilateral insula were both activated for one specific condition (e.g., Unfairness ∩ Core disgust), we would further examine the conjunction of the consensus connectivity maps of both seeds. Consistent with earlier studies (Bellucci et al., 2020; Camilleri et al., 2018), an additional extent-threshold of 10 continuous voxels was used in order to exclude smaller areas of putatively spurious overlap. Finally, the resulting regions of all consensus maps were identified using the SPM xjview toolbox (http://www.alivelearn.net/xjview/, v.9.7) and the automated anatomic labeling (AAL) atlas (Tzourio-Mazoyer et al., 2002).

### 2.9. Behavioral characterization based on the Neurosynth database

To infer mental processes most likely related to the identified brain regions in our meta-analyses, we performed a series of exploratory functional characterization analyses using data derived from Neurosynth and NiMARE (Salo et al., 2023; Yarkoni, Poldrack, Nichols, Van Essen, & Wager, 2011) which contain a large pool of automatically generated meta-analytic activation maps across a multitude of terms/topics. This approach enables us to discuss our results in relation to these terms/topics, without relying on acquiring data from a wide range of functional neuroimaging tasks in the same cohort. Consistent with the recommended input specifications as well as many recent publications (Gan et al., 2024; Maliske, Schurz, & Kanske, 2023; Pankey et al., 2022; Schurz et al., 2021), we subjected the unthresholded versions of ALE maps obtained from single dataset analyses and conjunction analyses for functional decoding. Here, we capitalized on the BrainStat toolbox (https://github.com/MICA-MNI/BrainStat, v.0.4.2) (Larivière et al., 2023) to run these analyses. We only displayed the top 100 most correlated terms/topics for clarity. Additionally, we excluded anatomic topics such as *dorsolateral prefrontal*, *anterior cingulate, etc*.

### 2.10. Behavioral characterization based on the BrainMap database

To complement the Neurosynth-based functional decoding, we conducted behavioral characterization analyses using the Behavioral Analysis plugin (Lancaster et al., 2012) implemented in Mango (http://ric.uthscsa.edu/mango/, v.4.1). This plugin allows us to access the meta-data of tens of thousands of neuroimaging experiments from the BrainMap database, where each experiment/contrast has been categorized into one of five behavioral domains (Action, Cognition, Emotion, Interoception, and Perception) and their related sub-domains. Briefly, the algorithm of behavioral analysis determines whether the number of foci related to a specific sub-domain is more/less likely to be located within the selected brain seeds than within the whole-brain space. Here, we conducted a series of behavioral analyses on the results from single dataset analyses and conjunction analyses at the whole-brain ALE map level (for a similar approach, see Liloia et al., 2021), only those sub-domains with z-score ≥ 3 (corresponding to p < 0.05) were reported (Lancaster et al., 2012).

### 2.11. Serotonergic decoding of the meta-analytic ALE maps

In a final step, based on the literature (Crockett et al., 2013; Crockett, Clark, Hauser, et al., 2010; Crockett, Clark, Lieberman, et al., 2010; Crockett et al., 2008; Vicario et al., 2017) we performed exploratory hypothesis-driven analyses to evaluate the relationship between the meta-analytic ALE maps and serotonin neurotransmitter systems (receptors 5HT1a, 5HT1b, 5HT2a, 5HT4, and 5HT6; and transporter 5HTT) which were acquired through positron emission tomography (PET) (recently compiled by Hansen et al., 2022; and available at https://github.com/netneurolab/hansen_receptors). Pearson’s correlations between the neurotransmitter receptor/transporter maps and unthresholded ALE maps obtained from single dataset analyses as well as conjunction analyses were calculated. Given that the PET distributions are different (Andrea et al., 2023), the significance of these correlations was examined using 10,000 spin-based permutation tests (p-spin) controlling for spatial autocorrelation (Alexander-Bloch et al., 2018; Shanmugan et al., 2022). Correlations were considered significant at p < 0.05.

## 3. Results

### 3.1. Single dataset analyses: determining robust brain systems underlying unfairness, core disgust, and anger provocation

#### 3.1.1. The unfairness network

The ALE meta-analysis for unfairness revealed five clusters of convergent brain activation that were robustly associated in response to unfairness in the UG context (Table 2 and Figure 2a). The clusters were located in bilateral anterior cingulate cortex (ACC), middle cingulate cortex (MCC), dorsolateral prefrontal cortex (dLPFC), AI and MI spreading into the left putamen, orbitofrontal cortex (OFC), ventrolateral prefrontal cortex (vLPFC), claustrum, dorsomedial prefrontal cortex (dMPFC), and left precentral gyrus.

**Figure 2.**
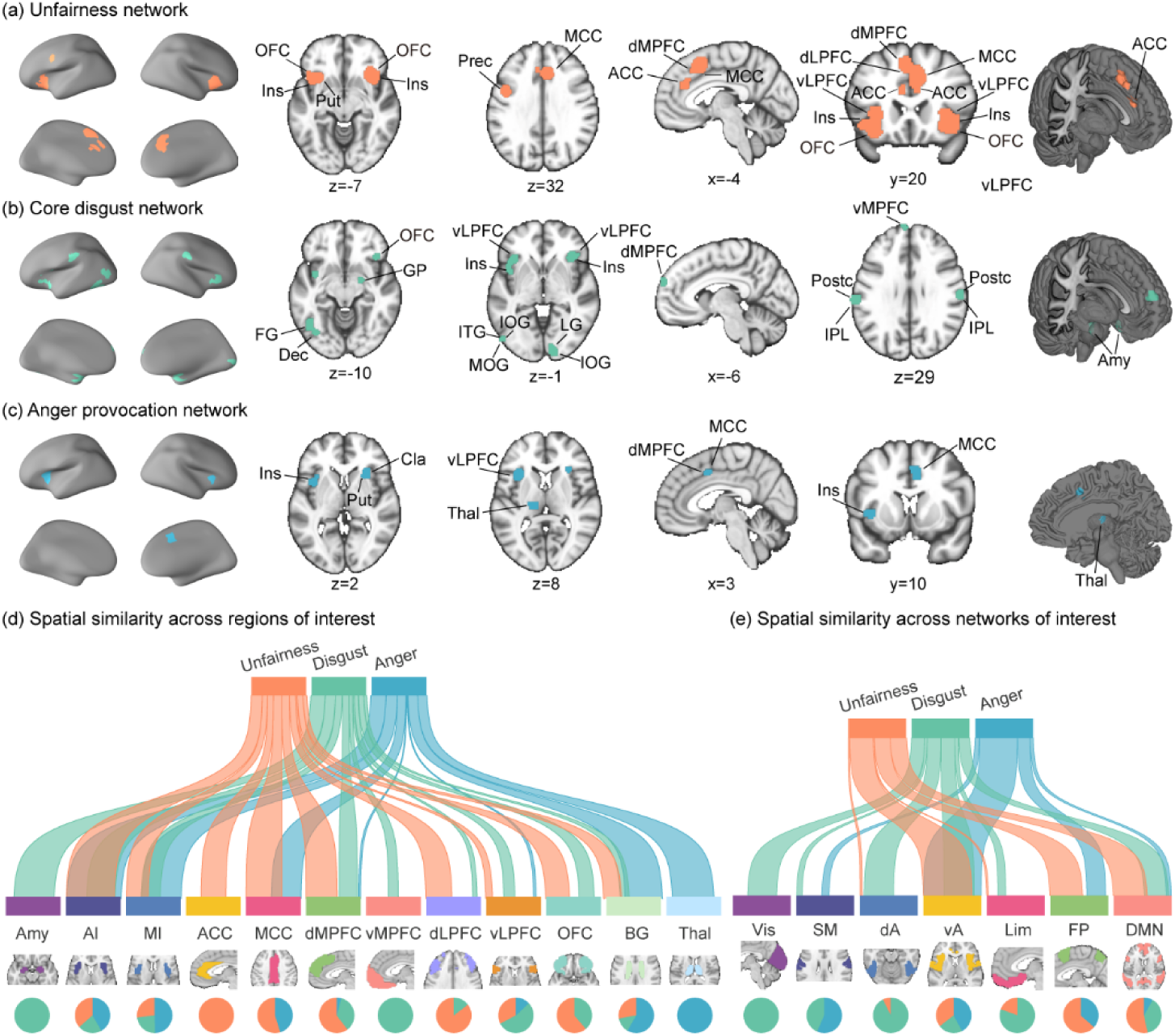
Overview of significant brain regions from single dataset analyses. (a) Unfairness network in healthy subjects. (b) Core disgust network in healthy subjects. (c) Anger provocation network in healthy subjects. (d-e) River plots showing spatial similarity (cosine similarity) between respective ALE-derived maps and selected brain systems. Ribbons are normalized by the max cosine similarity across all ROIs (d) and NOIs (e). Ribbon locations in relation to the boxes are arbitrary. Pie charts show relative contributions of each ALE-derived map to each ROI or network (i.e., percentage of voxels with the highest cosine similarity for each map). Coordinates are in the MNI space. Abbreviations: Amy = Amygdala, BG = basal ganglia, Cla = Claustrum, dA = dorsal attention network, Dec = Declive, FG = Fusiform Gyrus, FP = frontoparietal network, GP = Globus Pallidus, Ins = Insula, IOG = Inferior Occipital Gyrus, IPL = Inferior Parietal Lobule, ITG = Inferior Temporal Gyrus, LG = Lingual Gyrus, Limb = limbic network, MOG = Middle Occipital Gyrus, Postc = Postcentral Gyrus, Prec = Precentral Gyrus, Put = Putamen, SM = somatomotor network, Thal = Thalamus, vA = ventral attention network, Vis = visual network.

**Table 2.** Brain regions activated by unfairness in healthy subjects.

| Volume<br>(mm<sup>3</sup>) | Side | Brain region | BA | Peak voxel<br>location (x, y, z) |  |  | ALE<br>value<br>(10<sup>-2</sup>) | Z |
| --- | --- | --- | --- | --- | --- | --- | --- | --- |
| 6440 | L | dMPFC | 6/8 | -4 | 18 | 48 | 5.61 | 8.37 |
|  | R | MCC/ACC | 32/24 | 6 | 26 | 34 | 4.97 | 7.72 |
|  | R | MCC | 32 | 8 | 28 | 24 | 2.94 | 5.44 |
| 4056 | R | AI/MI/Clastrum/OFC/<br>dLPFC/vLPFC | 13/47/45 | 36 | 22 | -4 | 4.26 | 6.98 |
| 4016 | L | AI/MI/Clastrum/OFC/<br>vLPFC/Putamen | 13/47 | -32 | 18 | -10 | 4.56 | 7.30 |
|  | L | AI/MI/OFC/vLPFC | 13/47 | -30 | 22 | 4 | 2.79 | 5.26 |
| 912 | L | MCC/ACC | 32/24/9 | -6 | 36 | 24 | 2.09 | 4.33 |
|  | L | MCC/dMPFC/dLPFC | 32/9 | -8 | 24 | 24 | 1.62 | 3.62 |
| 832 | L | Precentral Gyrus | 6/9 | -44 | 4 | 32 | 2.97 | 5.47 |

Abbreviations: BA = Brodmann Area, L = left hemisphere, R = right hemisphere.
Voxel coordinates are in the MNI space.

#### 3.1.2. The core disgust network

In line with a recent meta-analysis focused on visually induced disgust (Gan et al., 2022), the multimodal core disgust network robustly engaged ten clusters of convergent activation (Table 3 and Figure 2b), including the bilateral amygdala (spreading into the right globus pallidus), parahippocampal gyrus, AI, MI, vLPFC, OFC, dMPFC, inferior parietal lobule, postcentral gyrus, left claustrum, left dLPFC, left ventromedial prefrontal cortex (vMPFC), left declive, left inferior temporal gyrus, left middle temporal gyrus, left fusiform gyrus, left middle occipital gyrus, and bilateral inferior occipital gyrus extending into the right lingual gyrus.

**Table 3.** Brain regions activated by core disgust in healthy subjects.

| Volume<br>(mm<sup>3</sup>) | Side | Brain region | BA | Peak voxel<br>location (x, y, z) |  |  | ALE<br>value<br>(10<sup>-2</sup>) | Z |
| --- | --- | --- | --- | --- | --- | --- | --- | --- |
| 2184 | L | Amygdala/Parahippocampal<br>Gyrus |  | -22 | -2 | -22 | 3.89 | 6.41 |
| 1760 | R | Amygdala/Parahippocampal<br>Gyrus |  | 22 | -4 | -16 | 3.47 | 5.91 |
|  | R | Amygdala/Globus Pallidus |  | 24 | -2 | -24 | 2.37 | 4.50 |
| 1712 | L | AI | 13 | -36 | 24 | -2 | 2.32 | 4.43 |
|  | L | Clastrum |  | -36 | 4 | -10 | 1.90 | 3.82 |
|  | L | MI/vLPFC/OFC/dLPFC | 13/45/47 | -40 | 8 | -4 | 1.88 | 3.80 |
| 1512 | L | Inferior Parietal Lobule/<br>Postcentral Gyrus | 40/2 | -62 | -24 | 34 | 3.99 | 6.53 |
| 1472 | L | Declive |  | -34 | -70 | -14 | 2.53 | 4.72 |
|  | L | Inferior Temporal Gyrus | 19 | -46 | -62 | -6 | 2.10 | 4.12 |
|  | L | Fusiform Gyrus | 37 | -42 | -54 | -18 | 1.69 | 3.51 |
| 1344 | L | Middle Occipital<br>Gyrus/Inferior Occipital Gyrus | 19/18 | -44 | -76 | 8 | 2.48 | 4.65 |
|  | L | Inferior Temporal<br>Gyrus/Middle Temporal Gyrus | 19/37 | -50 | -76 | 0 | 2.02 | 4.00 |
| 1280 | R | AI/MI/vLPFC/OFC | 13/47/45 | 38 | 26 | -4 | 2.77 | 5.04 |
| 1016 | L | dMPFC | 9 | -6 | 64 | 26 | 2.21 | 4.28 |
|  | L | dMPFC/vMPFC | 9/10 | 2 | 62 | 18 | 2.14 | 4.17 |
| 960 | R | Inferior Occipital Gyrus/<br>Lingual Gyrus | 17/18 | 14 | -92 | -2 | 2.64 | 4.87 |
| 776 | R | Postcentral Gyrus/Inferior<br>Parietal Lobule | 2/40 | 62 | -18 | 30 | 2.57 | 4.77 |

Voxel coordinates are in the MNI space.

#### 3.1.3. The anger provocation network

The ALE map for anger provocation revealed four significant clusters of convergent brain activation located in the bilateral AI and MI spreading into the right putamen, claustrum, left thalamus, left vLPFC, right MCC, and adjacent dMPFC (Table 4 and Figure 2c).

**Table 4.** Brain regions activated by anger provocation in healthy subjects.

| Volume<br>(mm<sup>3</sup>) | Side | Brain region | BA | Peak voxel<br>location (x, y, z) |  |  | ALE<br>value<br>(10<sup>-2</sup>) | Z |
| --- | --- | --- | --- | --- | --- | --- | --- | --- |
| 1672 | L | AI/vLPFC | 13/45 | -34 | 18 | 8 | 3.23 | 5.94 |
|  | L | MI/Clastrum | 13 | -38 | 10 | 2 | 1.89 | 4.14 |
| 928 | R | AI/MI/Clastrum/Putamen |  | 30 | 20 | 2 | 2.64 | 5.19 |
| 752 | L | Thalamus |  | -14 | -22 | 8 | 2.13 | 4.48 |
| 712 | R | MCC | 24 | 8 | 8 | 42 | 1.88 | 4.13 |
|  | R | MCC/dMPFC | 32/24 | 8 | 8 | 38 | 1.76 | 3.95 |

Voxel coordinates are in the MNI space.

#### 3.1.4. Spatial similarity between respective ALE-derived maps and literature-derived brain systems

The spatial similarity analyses across a set of ROIs as well as NOIs showed that the AI, MI, dMPFC, vLPFC, basal ganglia, ventral attention network (salience network), and default mode network (DMN) contained significant voxels across all three maps. The dLPFC, OFC, dorsal attention network, and limbic network encompassed significant voxels in both unfairness and core disgust maps, and the MCC and frontoparietal network included significant voxels in both unfairness and anger provocation maps, while the somatomotor network comprised significant voxels in both core disgust and anger provocation maps. Furthermore, the ACC was associated with unfairness, and the amygdala, vMPFC, as well as the visual network showed specificity to core disgust, whereas the thalamus uniquely contributed to anger provocation (Figure 2d-e).

### 3.2. Contrast and conjunction analyses

#### 3.2.1 Common and distinct brain regions for unfairness and core disgust

The conjunction analysis for unfairness and core disgust revealed that the bilateral AI and MI extending into vLPFC and OFC were robustly activated during both processes (Table 5 and Figure 3a).

**Figure 3.**
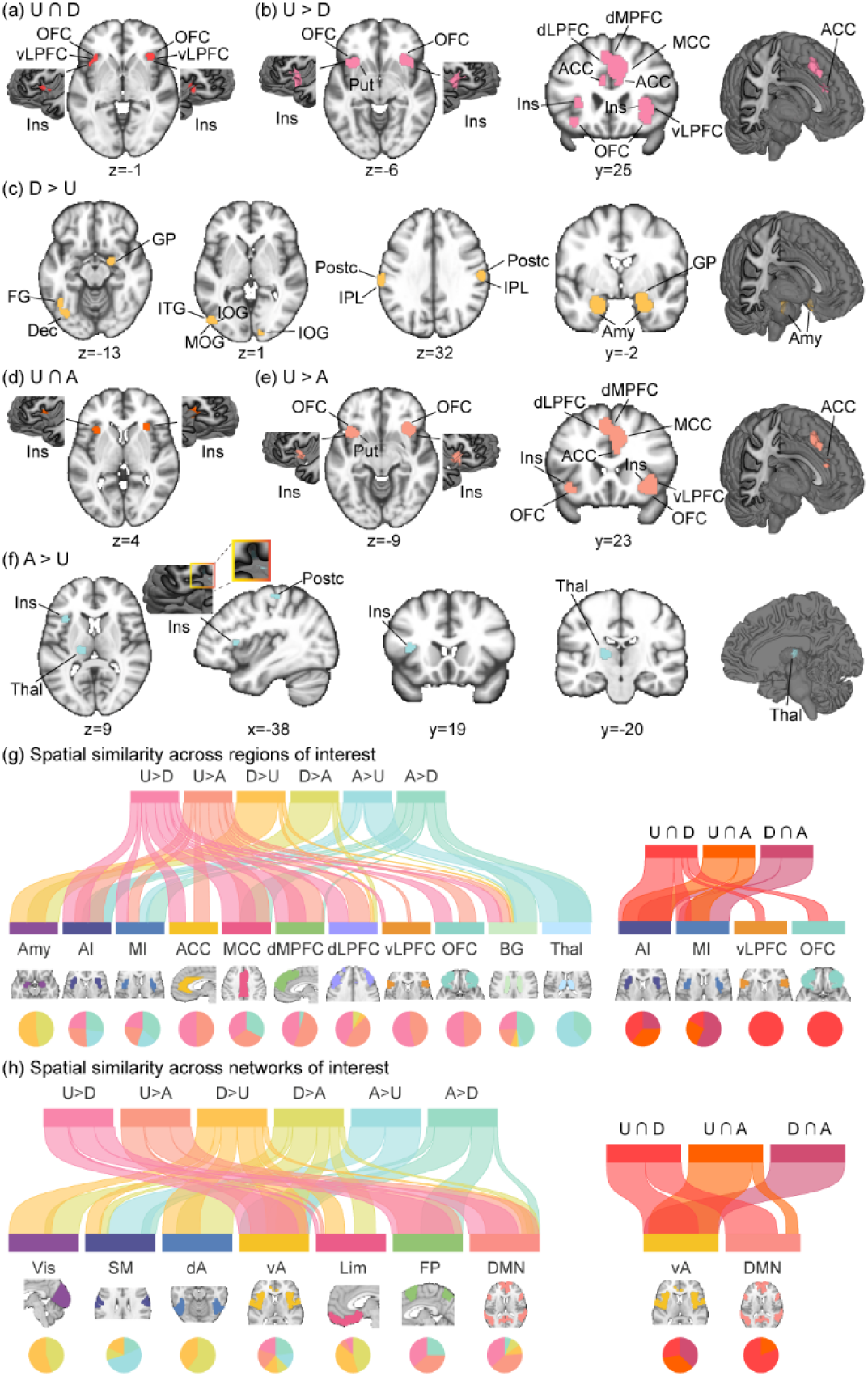
Overview of significant brain regions from conjunction and contrast analyses. (a) Common activations for unfairness and core disgust in healthy subjects. (b) Greater activations for unfairness than for core disgust (unfairness > core disgust) in healthy subjects. (c) Greater activations for core disgust than for unfairness (core disgust > unfairness) in healthy subjects. (d) Common activations for unfairness and anger provocation in healthy subjects. (e) Greater activations for unfairness than for anger provocation (unfairness > anger provocation) in healthy subjects. (f) Greater activations for anger provocation than for unfairness (anger provocation > unfairness) in healthy subjects. (g-h) River plots showing spatial similarity (cosine similarity) between respective contrast or conjunction maps and selected brain systems. Ribbons are normalized by the max cosine similarity across all ROIs (g) and NOIs (h). Ribbon locations in relation to the boxes are arbitrary. Pie charts show relative contributions of each map to each ROI or network (i.e., percentage of voxels with the highest cosine similarity for each map). Note that some brain systems presented in Figure 2d-e but not in Figure 3g-h, due to the fact that these systems have no spatial similarity to the corresponding maps. Coordinates are in the MNI space.

**Table 5.**
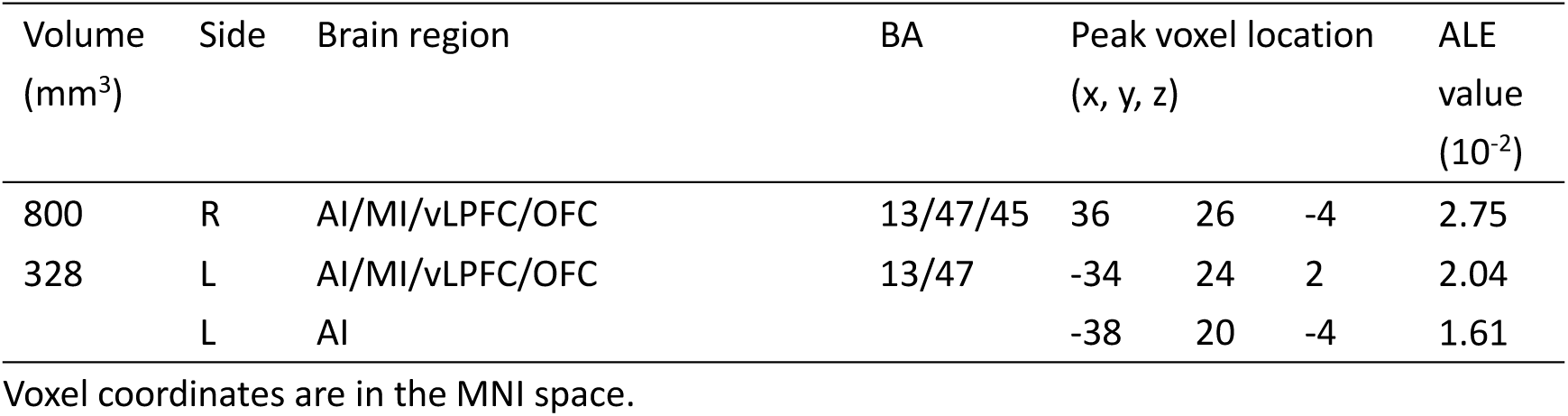
Brain regions activated by conjunction analysis between unfairness and core disgust in healthy subjects.

| Volume (mm<sup>3</sup>) | Side | Brain region | BA | Peak voxel location (x, y, z) |  |  | ALE value (10<sup>-2</sup>) |
| --- | --- | --- | --- | --- | --- | --- | --- |
| 800 | R | AI/MI/vLPFC/OFC | 13/47/45 | 36 | 26 | -4 | 2.75 |
| 328 | L | AI/MI/vLPFC/OFC | 13/47 | -34 | 24 | 2 | 2.04 |
|  | L | AI |  | -38 | 20 | -4 | 1.61 |

Voxel coordinates are in the MNI space.

The contrast analyses revealed stronger recruitment of the bilateral ACC, MCC, dMPFC, dLPFC, AI, and MI (spreading into adjacent claustrum), OFC, right vLPFC, and left putamen for unfairness compared with core disgust (Table 6 and Figure 3b). By contrast, core disgust processing was characterized by a stronger engagement of the bilateral limbic-striatal cluster (including the amygdala, parahippocampal gyrus, and right globus pallidus), inferior parietal lobule, postcentral gyrus, inferior occipital gyrus, left fusiform gyrus, left declive, left middle occipital gyrus, left inferior temporal gyrus, and left middle temporal gyrus as compared to unfairness (Table 6 and Figure 3c).

**Table 6.**
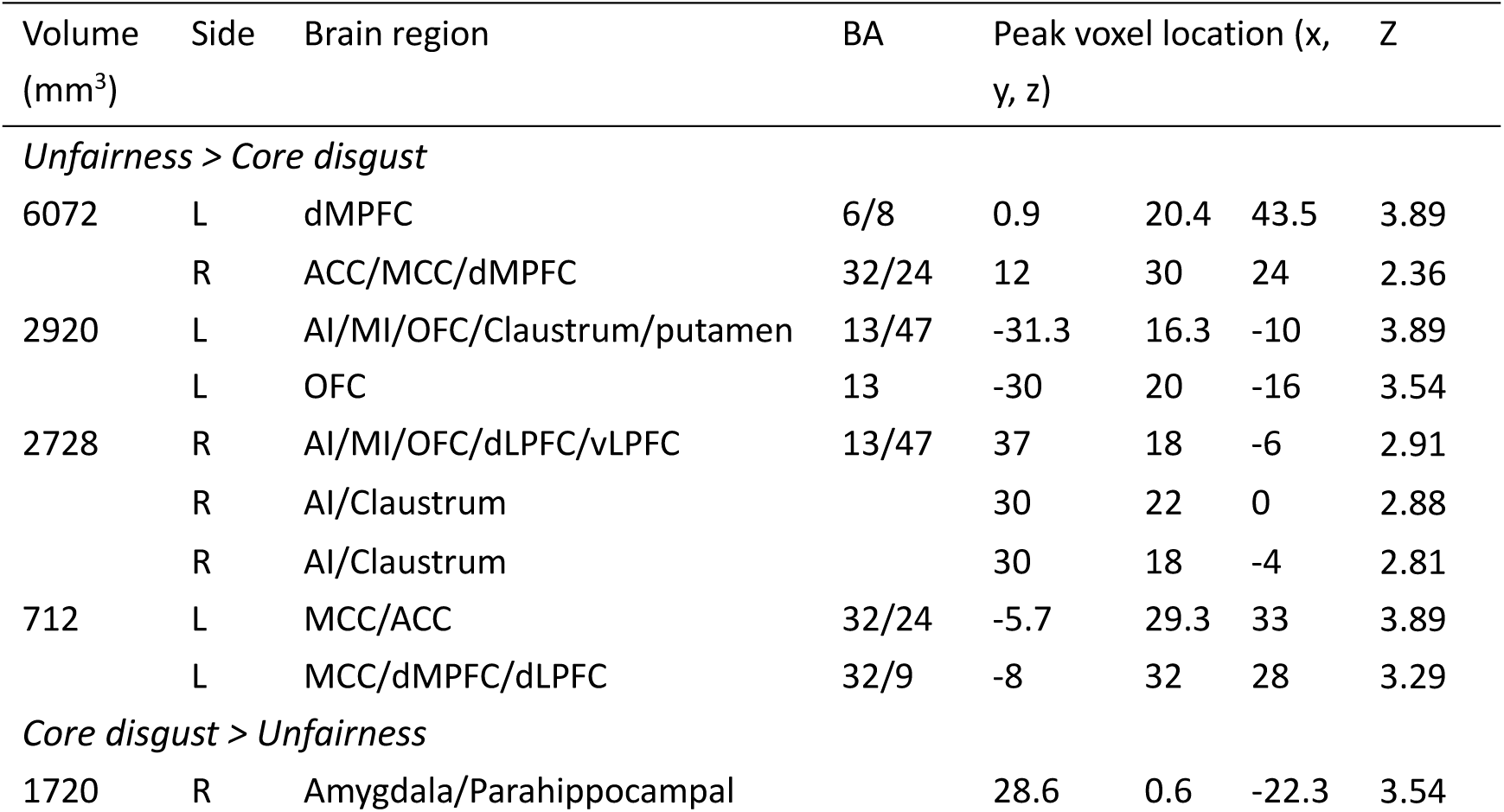
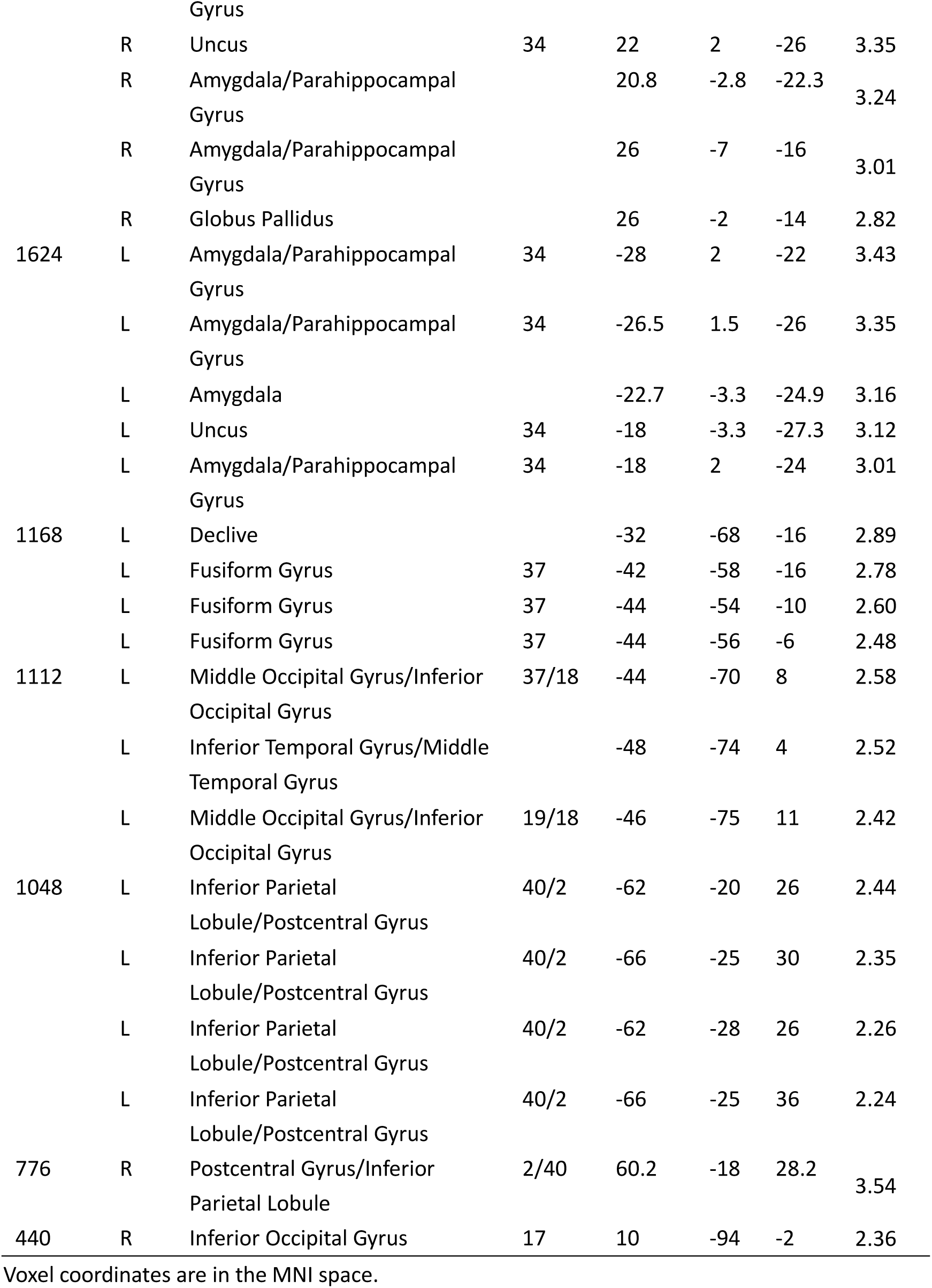
Brain regions activated by contrast analysis between unfairness and core disgust in healthy subjects.

| Volume<br>(mm<sup>3</sup>) | Side | Brain region | BA | Peak voxel location (x, y, z) |  |  |  | Z |
| --- | --- | --- | --- | --- | --- | --- | --- | --- |
| *Unfairness > Core disgust* |  |  |  |  |  |  |  |  |
| 6072 | L | dMPFC | 6/8 | 0.9 | 20.4 | 43.5 |  | 3.89 |
|  | R | ACC/MCC/dMPFC | 32/24 | 12 | 30 | 24 |  | 2.36 |
| 2920 | L | AI/MI/OFC/Claustrum/putamen | 13/47 | -31.3 | 16.3 | -10 |  | 3.89 |
|  | L | OFC | 13 | -30 | 20 | -16 |  | 3.54 |
| 2728 | R | AI/MI/OFC/dLPFC/vLPFC | 13/47 | 37 | 18 | -6 |  | 2.91 |
|  | R | AI/Claustrum |  | 30 | 22 | 0 |  | 2.88 |
|  | R | AI/Claustrum |  | 30 | 18 | -4 |  | 2.81 |
| 712 | L | MCC/ACC | 32/24 | -5.7 | 29.3 | 33 |  | 3.89 |
|  | L | MCC/dMPFC/dLPFC | 32/9 | -8 | 32 | 28 |  | 3.29 |
| *Core disgust > Unfairness* |  |  |  |  |  |  |  |  |
| 1720 | R | Amygdala/Parahippocampal |  | 28.6 | 0.6 | -22.3 |  | 3.54 |

|  |  |  |  |  |  |  |  |
| --- | --- | --- | --- | --- | --- | --- | --- |
|  |  | Gyrus |  |  |  |  |  |
|  | R | Uncus | 34 | 22 | 2 | -26 | 3.35 |
|  | R | Amygdala/Parahippocampal Gyrus |  | 20.8 | -2.8 | -22.3 | 3.24 |
|  | R | Amygdala/Parahippocampal Gyrus |  | 26 | -7 | -16 | 3.01 |
|  | R | Globus Pallidus |  | 26 | -2 | -14 | 2.82 |
| 1624 | L | Amygdala/Parahippocampal Gyrus | 34 | -28 | 2 | -22 | 3.43 |
|  | L | Amygdala/Parahippocampal Gyrus | 34 | -26.5 | 1.5 | -26 | 3.35 |
|  | L | Amygdala |  | -22.7 | -3.3 | -24.9 | 3.16 |
|  | L | Uncus | 34 | -18 | -3.3 | -27.3 | 3.12 |
|  | L | Amygdala/Parahippocampal Gyrus | 34 | -18 | 2 | -24 | 3.01 |
| 1168 | L | Declive |  | -32 | -68 | -16 | 2.89 |
|  | L | Fusiform Gyrus | 37 | -42 | -58 | -16 | 2.78 |
|  | L | Fusiform Gyrus | 37 | -44 | -54 | -10 | 2.60 |
|  | L | Fusiform Gyrus | 37 | -44 | -56 | -6 | 2.48 |
| 1112 | L | Middle Occipital Gyrus/Inferior Occipital Gyrus | 37/18 | -44 | -70 | 8 | 2.58 |
|  | L | Inferior Temporal Gyrus/Middle Temporal Gyrus |  | -48 | -74 | 4 | 2.52 |
|  | L | Middle Occipital Gyrus/Inferior Occipital Gyrus | 19/18 | -46 | -75 | 11 | 2.42 |
| 1048 | L | Inferior Parietal Lobule/Postcentral Gyrus | 40/2 | -62 | -20 | 26 | 2.44 |
|  | L | Inferior Parietal Lobule/Postcentral Gyrus | 40/2 | -66 | -25 | 30 | 2.35 |
|  | L | Inferior Parietal Lobule/Postcentral Gyrus | 40/2 | -62 | -28 | 26 | 2.26 |
|  | L | Inferior Parietal Lobule/Postcentral Gyrus | 40/2 | -66 | -25 | 36 | 2.24 |
| 776 | R | Postcentral Gyrus/Inferior Parietal Lobule | 2/40 | 60.2 | -18 | 28.2 | 3.54 |
| 440 | R | Inferior Occipital Gyrus | 17 | 10 | -94 | -2 | 2.36 |

Voxel coordinates are in the MNI space.

#### 3.2.2 Common and distinct brain regions for unfairness and anger provocation

The conjunction analysis for unfairness and anger provocation revealed that the bilateral AI and MI were robustly activated during both processes (Table 7 and Figure 3d).

**Table 7.**
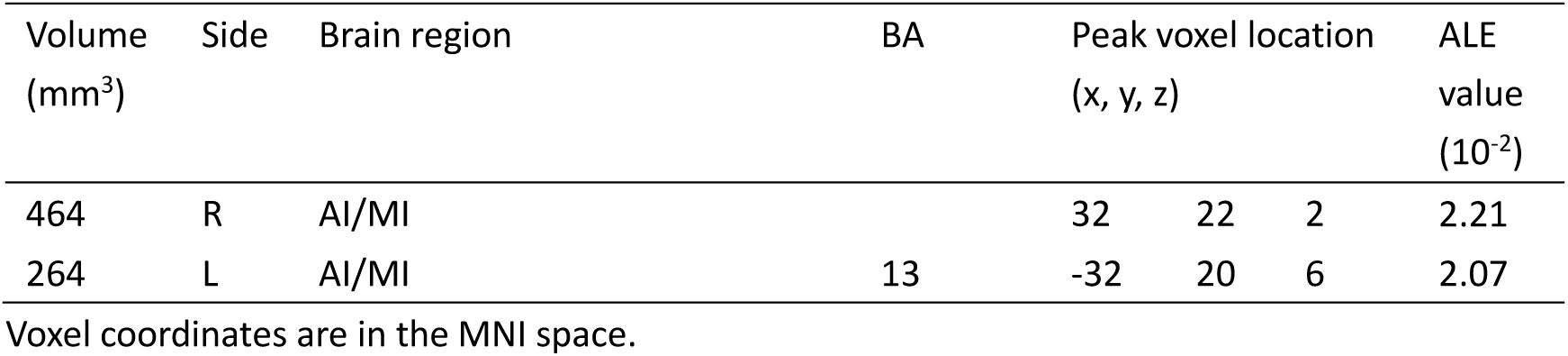
Brain regions activated by conjunction analysis between unfairness and anger provocation in healthy subjects.

| Volume (mm<sup>3</sup>) | Side | Brain region | BA | Peak voxel location (x, y, z) |  |  | ALE value (10<sup>-2</sup>) |
| --- | --- | --- | --- | --- | --- | --- | --- |
| 464 | R | AI/MI |  | 32 | 22 | 2 | 2.21 |
| 264 | L | AI/MI | 13 | -32 | 20 | 6 | 2.07 |

Voxel coordinates are in the MNI space.

The contrast analyses revealed a stronger engagement of cortical and insular regions, including the bilateral ACC, MCC, dMPFC, dLPFC, AI, and MI (spreading into adjacent OFC and claustrum), and right vLPFC for unfairness compared with anger provocation (Table 8 and Figure 3e). In contrast, anger provocation was characterized by stronger recruitment of the left thalamus, left AI and MI, and left postcentral gyrus as compared to unfairness (Table 8 and Figure 3f).

**Table 8.** Brain regions activated by contrast analysis between unfairness and anger provocation in healthy subjects.

| Volume<br>(mm<sup>3</sup>) | Side | Brain region | BA | Peak voxel location<br>(x, y, z) |  |  | Z |
| --- | --- | --- | --- | --- | --- | --- | --- |
| *Unfairness > Anger provocation* |  |  |  |  |  |  |  |
| 4920 | L | dMPFC | 6/8 | -3.2 | 22.2 | 47.8 | 3.89 |
|  | L | dMPFC | 6/8 | -8.7 | 15.1 | 52.7 | 3.72 |
|  | R | MCC/ACC | 32/24 | 10 | 22 | 40 | 3.35 |
|  | R | MCC/ACC | 32/24 | 8 | 22 | 30 | 2.99 |
| 2176 | R | AI/MI/Claustrum/OFC/<br>dLPFC/vLPFC | 13/45/47 | 36 | 18 | -9 | 3.35 |
| 1440 | L | OFC/AI | 47 | -40 | 17 | -12 | 3.35 |
|  | L | AI/MI/Claustrum/OFC/Putamen | 13 | -38 | 18 | -8 | 3.29 |
| 256 | L | ACC/MCC | 32/24 | -8 | 30 | 22 | 2.00 |
|  | L | dMPFC/dLPFC | 9 | -8 | 36 | 28 | 1.94 |
| *Anger provocation > Unfairness* |  |  |  |  |  |  |  |
| 608 | L | Thalamus |  | -16 | -18 | 11.2 | 2.75 |
|  | L | Thalamus |  | -18 | -20 | 8 | 2.74 |
| 280 | L | AI/MI | 13 | -38 | 18 | 10 | 1.91 |
| 256 | L | Postcentral Gyrus | 3/40 | -42 | -26 | 60 | 2.32 |

Voxel coordinates are in the MNI space.

#### 3.2.3 Common and distinct brain regions for core disgust and anger provocation

The conjunction analysis for core disgust and anger provocation revealed that the left MI and AI were robustly activated during both processes (Supplemental Materials Table S2 and Supplemental Materials Figure S2a).

The contrast analyses revealed stronger involvement of the bilateral AI and MI (extending into adjacent claustrum), left thalamus, right putamen, and right MCC (spreading into dMPFC) for anger provocation as compared to core disgust (Supplemental Materials Table S3 and Supplemental Materials Figure S2b). By contrast, core disgust processing was characterized by a stronger engagement of the bilateral amygdala, parahippocampal gyrus, inferior parietal lobule, postcentral gyrus, right globus pallidus, right inferior occipital gyrus and left fusiform gyrus compared with anger provocation (Supplemental Materials Table S3 and Supplemental Materials Figure S2c).

#### 3.2.4. Common brain regions activated across unfairness, core disgust, and anger provocation

A grand conjunction across unfairness, core disgust, and anger provocation revealed no meaningful overlap, in other words, we did not yield shared neural substrates across all the domains.

#### 3.2.5. Spatial similarity between respective maps and literature-derived brain systems

The spatial similarity analyses between the maps obtained from contrast analyses and a set of ROIs as well as NOIs revealed that the ACC, dMPFC, dLPFC, vLPLFC, and OFC were specific for unfairness (i.e., domain-specific for unfairness) because significant voxels from the five ROIs are almost only implicated in unfairness > disgust and unfairness > anger provocation; the amygdala, visual network, and dorsal attention network were specific for core disgust (domain-specific core disgust); while the thalamus was specific for anger provocation (domain-specific anger provocation). Moreover, the AI, MI, MCC, and frontoparietal network exhibited no contribution to core disgust compared with unfairness or anger provocation; the MCC and frontoparietal network contributed more to unfairness as compared to anger provocation; and the limbic network showed no contribution to anger provocation (yet the limbic network contributed stronger to core disgust than to unfairness). Furthermore, the somatomotor network failed to contribute to unfairness compared with disgust or anger provocation, and the DMN showed no contribution to anger provocation compared with unfairness. In addition, the basal ganglia and ventral attention network contained significant voxels in all six contrast maps, whereas the former was strongly involved in anger provocation and unfairness (Figure 3g-h).

The spatial similarity analyses between the conjunction maps and a set of ROIs as well as NOIs showed that the AI, MI, and ventral attention network included significant voxels in all three conjunction maps; and the DMN contributed to the conjunction of unfairness and core disgust as well as the conjunction of unfairness and anger provocation; while the vLPFC and OFC uniquely contributed to the conjunction of unfairness and disgust (Figure 3g-h). A further breaking down of the vLPFC revealed that its left side specifically contributed to the conjunction of unfairness and core disgust, whereas its right side contributed more to the conjunction than to the contrast of unfairness and core disgust (Supplemental Materials Figure S3a); the breaking down of the OFC showed that its left side contributed more to the contrast of unfairness and core disgust, while its right side contributed more to the conjunction of unfairness and core disgust (Supplemental Materials Figure S3b).

### 3.3. Consensus connectivity maps of MACM and RSFC profiles

The consensus connectivity map of the bilateral insula from the conjunction of unfairness and core disgust was determined based on the connectivity profiles of MACM and RSFC analyses. This analysis yielded a consensus connectivity network of the bilateral AI during both task and resting states, including clusters mainly converged at the salience network and DMN (Supplemental Materials Table S4 and Figure 4a). Likewise, the consensus connectivity map of the bilateral AI from the conjunction of unfairness and anger provocation also encompassed two main networks: the salience network and DMN (Supplemental Materials Table S5 and Figure 4b). While the consensus connectivity map of the left MI from the conjunction of core disgust and anger provocation comprised of the salience network and DMN, it also contained motor regions from the parietal lobe such as the postcentral gyrus which forms the somatomotor network, inferior parietal lobule, and supramarginal gyrus (Supplemental Materials Table S6 and Figure 4c). See also Supplemental Materials Figure S4 for the detailed methodological workflow of obtaining respective consensus connectivity maps and Supplemental Materials Figure S5 for the network-level assignment of corresponding consensus connectivity maps.

**Figure 4.**
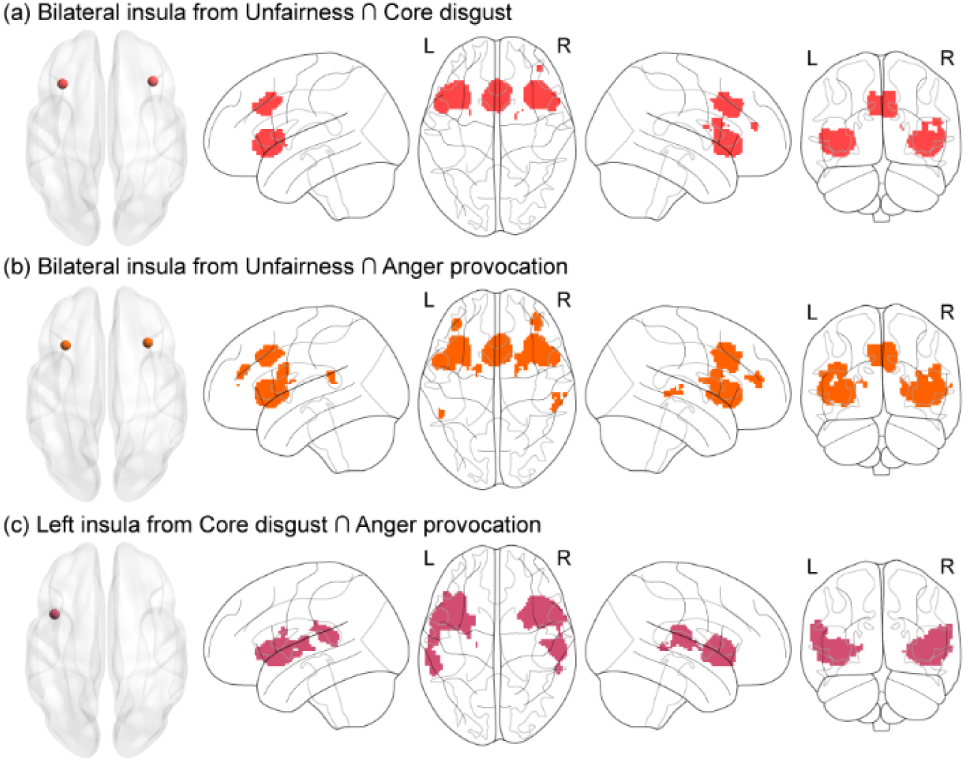
Consensus connectivity networks for the insula from conjunction analyses. (a) Results of consensus connectivity map for bilateral AI from the conjunction of unfairness and core disgust. (b) Results of consensus connectivity map for bilateral AI from the conjunction of unfairness and anger provocation. (c) Results of consensus connectivity map for left MI from the conjunction of core disgust and anger provocation. Abbreviations: L = left, R = right.

Identifying the consensus connectivity map of the bilateral AI from the contrast of unfairness > core disgust revealed convergence in three networks: the salience network, DMN, and frontoparietal network (Supplemental Materials Table S7 and Figure 5a). The same was true for the consensus connectivity map of the bilateral AI from the contrast of unfairness > anger provocation (Supplemental Materials Table S8 and Figure 5b). Furthermore, both consensus connectivity maps (i.e., Figure 5a-b) did not contain robust engagement of motor-related areas, such as somatomotor regions.

**Figure 5.**
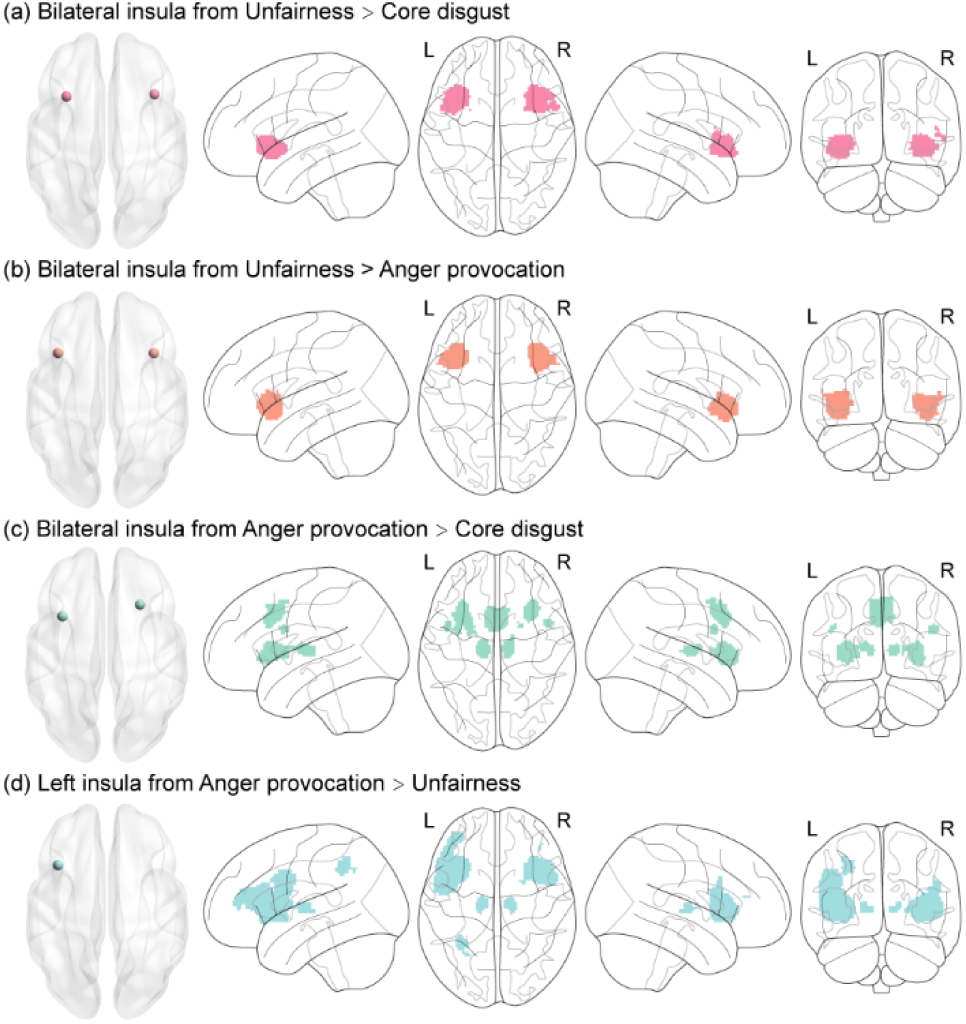
Consensus connectivity networks for the insula from contrast analyses. (a) Results of consensus connectivity map for bilateral AI from the contrast of unfairness > core disgust. (b) Results of consensus connectivity map for bilateral AI from the contrast of unfairness > anger provocation. (c) Results of consensus connectivity map for bilateral AI from the contrast of anger provocation > core disgust. (d) Results of consensus connectivity map for left AI from the contrast of anger provocation > unfairness.

Determining the consensus connectivity map of the bilateral AI from the contrast of anger provocation > core disgust showed convergence mainly at the salience network, with the frontoparietal network and DMN also contributing, albeit less than the salience network; moreover, the bilateral thalamus were robustly engaged (Supplemental Materials Table S9 and Figure 5c). The consensus connectivity map of the left AI from the contrast of anger provocation > unfairness exhibited the largest convergence at the salience network, with the motor-related regions such as the primary motor cortex (e.g., precentral gyrus) and inferior parietal lobule robustly involved. More importantly, consistent with the consensus connectivity map of the bilateral AI from anger provocation > core disgust (i.e., Figure 5c), the bilateral thalamus were significantly engaged (Supplemental Materials Table S10 and Figure 5d). See also Supplemental Materials Figure S6 for the detailed methodological workflow of obtaining respective consensus connectivity maps and Supplemental Materials Figure S7 for the network-level assignment of corresponding consensus connectivity maps.

### 3.4. Behavioral characterization results based on the Neurosynth database

Here, we capitalized on the Neurosynth database to functionally characterize the networks from single dataset analyses and conjunction analyses in terms of associations with other meta-analytic maps. The top 100 terms were displayed with a larger font size indicating a higher convergence (i.e., Pearson correlation coefficient). Briefly, decoding for the unfairness network from single dataset analysis showed terms primarily related to norm violation, decision-making, and salience (Figure 6a); for the core disgust regions, decoding revealed terms mainly associated with threat- (e.g., disgust), avoidance- and salience-related processes (Figure 6b); and for the anger provocation network, decoding showed terms mostly linked with negative and salient processes (Figure 6c).

**Figure 6.**
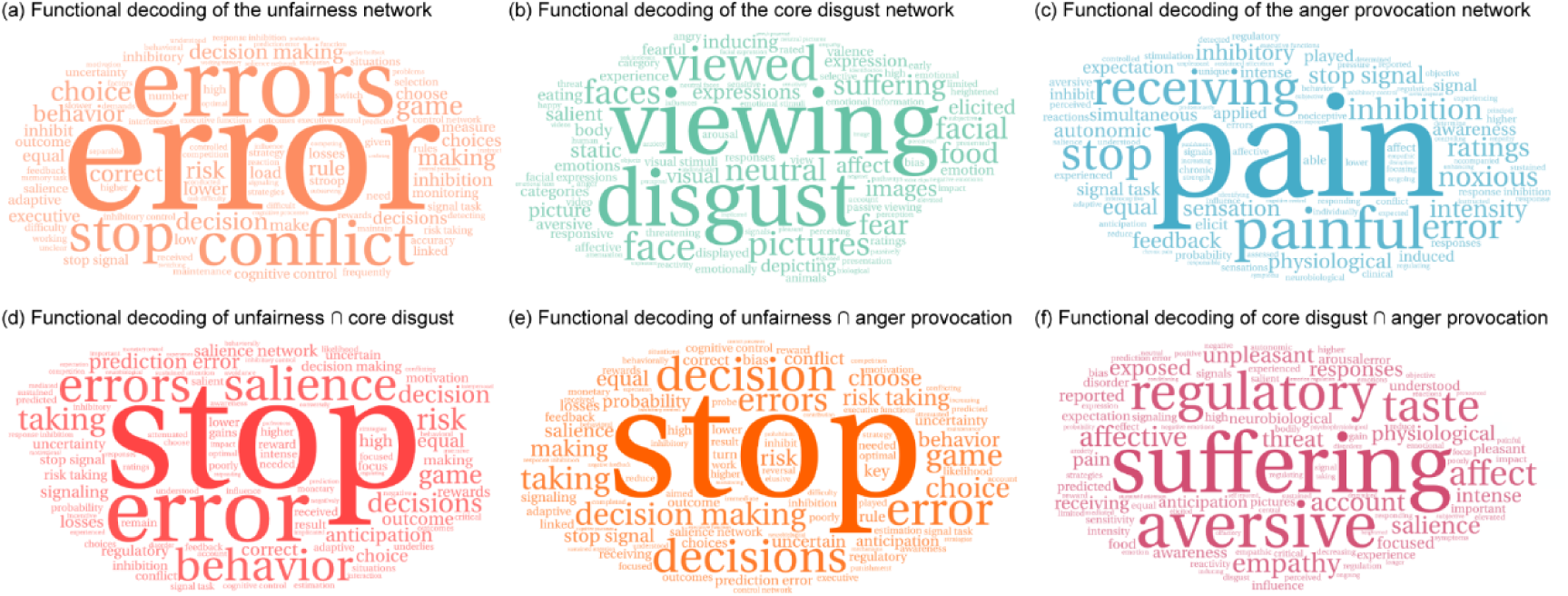
Neurosynth behavioral characterization of maps from single dataset analyses and conjunction analyses. (a) Functional decoding of the unfairness network from single dataset analysis. (b) Functional decoding of the core disgust network from single dataset analysis. (c) Functional decoding of the anger provocation network from single dataset analysis. (d) Functional decoding of unfairness ∩ core disgust from conjunction analysis. (e) Functional decoding of unfairness ∩ anger provocation from conjunction analysis. (f) Functional decoding of core disgust ∩ anger provocation from conjunction analysis. For each map, the 100 most strongly correlated terms were displayed, with a larger font size indicating a stronger correlation.

More interestingly, decoding for the conjunction of unfairness and core disgust (Figure 6d) as well as the conjunction of unfairness and anger provocation (Figure 6e) revealed mostly norm violation, decision-making, and salience-related terms, while the decoding for the conjunction of core disgust and anger provocation showed terms mainly related to negative affect and salience (Figure 6f).

See Supplemental Results and Supplemental Materials Figure S8 for the behavioral characterization results based on the BrainMap database.

### 3.5. Serotonergic profiling results

Finally, we explored the relationship between the meta-analytic ALE maps and serotonin receptors/transporters (Figure 7a and Supplemental Materials Table S11) across the neocortical cortex according to a recent work (Hansen et al., 2022). The 5HT1b significantly correlated with the unfairness network (Figure 7b), 5HT1b and 5HT2a significantly associated with the core disgust network (Figure 7c), and 5HTT significantly related to the anger provocation network (Figure 7d). Furthermore, 5HT1b and 5HTT were observed for the conjunction of unfairness and core disgust (Figure 7e), 5HTT was present in the conjunction of unfairness and anger provocation (Figure 7f), and 5HT6 and 5HTT were observed for the conjunction of core disgust and anger provocation (Figure 7g).

**Figure 7.**
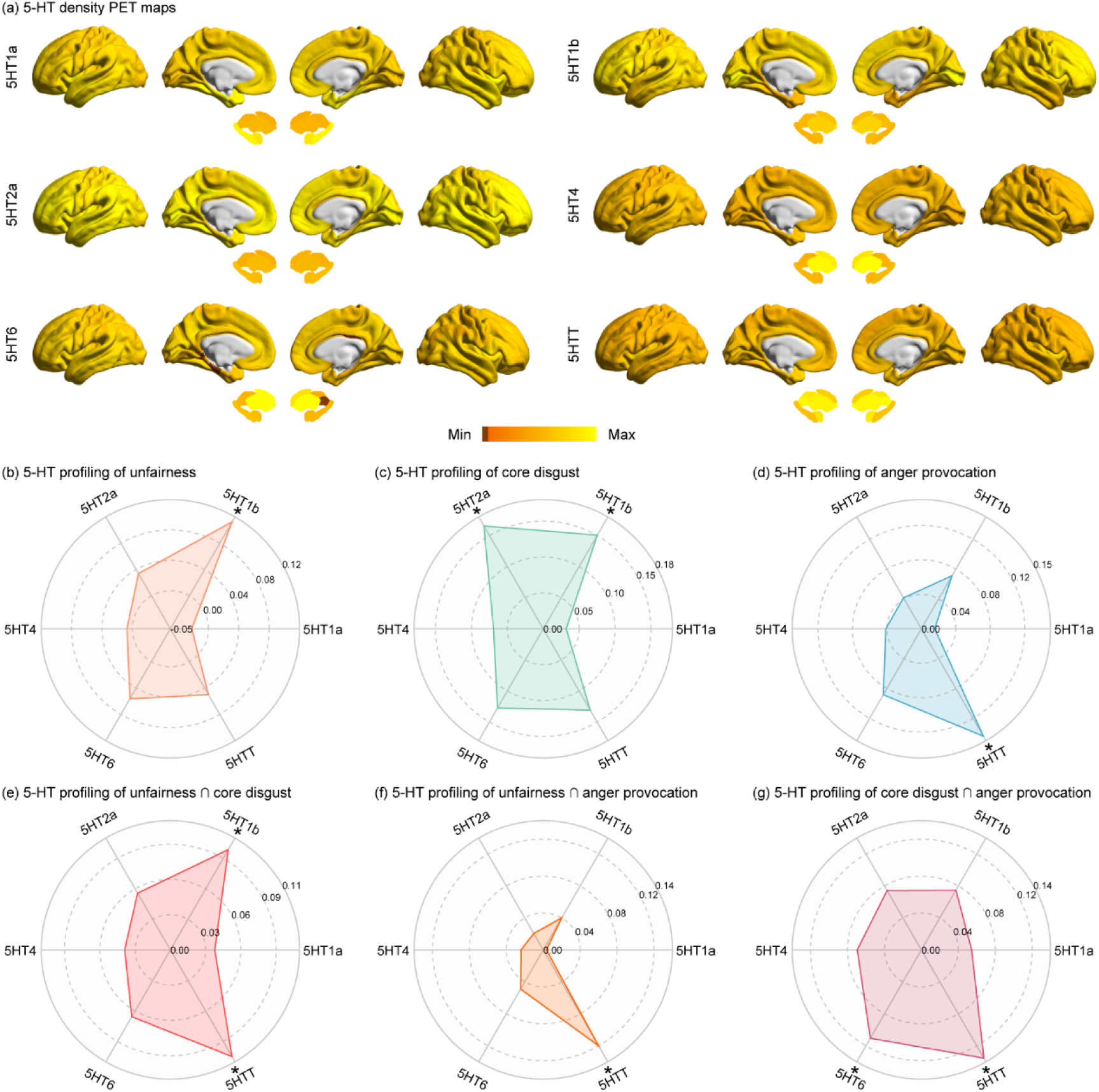
Serotonergic profiling of maps from single dataset analyses and conjunction analyses. (a) The parcellated and z-scored PET maps of serotonin receptors (5HT1a, 5HT1b, 5HT2a, 5HT4, and 5HT6) and transporter (5HTT). (b) Serotonin profiling of the unfairness network from single dataset analysis. (c) Serotonin profiling of the core disgust network from single dataset analysis. (d) Serotonin profiling of the anger provocation network from single dataset analysis. (e) Serotonin profiling of unfairness ∩ core disgust from conjunction analysis. (f) Serotonin profiling of unfairness ∩ anger provocation from conjunction analysis. (g) Serotonin profiling of core disgust ∩ anger provocation from conjunction analysis. * p-spin < 0.05.

### 3.6. The rejection network

The exploratory meta-analysis including 14 experiments on rejection showed increased activation in the left putamen during rejection as compared to acceptance (Supplemental Materials Table S12, Supplemental Materials Figure S9a). A series of conjunction analyses did not yield any overlapping between rejection and core disgust or between rejection and anger provocation. The contrast analyses revealed stronger involvement of the left putamen for rejection compared with core disgust (Supplemental Materials Table S13, Supplemental Materials Figure S9b), while core disgust showed stronger recruitment of the bilateral occipital lobe (e.g., inferior occipital gyrus, lingual gyrus, and middle occipital gyrus) and left parietal lobe (inferior parietal lobule and postcentral gyrus) as compared to rejection (Supplemental Materials Table S13, Supplemental Materials Figure S9b). However, the contrast analyses between rejection and anger provocation did not reveal any results. We further adopted the consensus connectivity approach described above to determine the connectivity profile of the left putamen. The results showed that the consensus connectivity map of the left putamen mainly included the salience network, DMN, frontoparietal, and somatomotor network (Supplemental Materials Table S14, Supplemental Materials Figure S10).

See Supplemental Results, Supplemental Materials Tables S15-S18, and Supplemental Materials Figure S11 for the results of using lenient thresholding to provide some trend of the rejection response.

## 4. Discussion

Fairness and upholding fairness norms constitute a central aspect of human moral behavior (Chapman et al., 2009; Hallsson et al., 2018; Henrich et al., 2004; Huebner et al., 2009; Sheskin, 2017; Tomasello, 2016). When confronted with violations of fairness norms, humans readily punish transgressors at a personal cost (Fehr & Fischbacher, 2003; Fehr & Gächter, 2002; McAuliffe et al., 2017). While this observation challenges both evolutionary and economic theories that portray humans as rational self-interest maximizers (Dawkins, 1989; Henrich et al., 2004; A. Smith, 1776; Stout, 2008), this phenomenon is well-illustrated in experimental paradigms from psychology and behavioral economics such as the UG paradigm, and an abundance of evidence suggests that negative emotions (especially two moral emotions, that is, disgust and anger) elicited by unfairness drive the response to rejections (costly punishment) (Bargh et al., 2012; Chapman & Anderson, 2013; Haidt, 2001; Hastie, 2001; McAuliffe et al., 2017; Oatley & Johnson-Laird, 2014; Rozin et al., 2009; Russell & Giner-Sorolla, 2013; Vavra et al., 2017; Yamagishi et al., 2009). The current coordinate-based meta-analyses capitalized on a large number of studies that examined the underlying brain mechanism to determine the neurobiological basis of the interactions between unfairness, anger, and disgust and to determine whether these mental processes are mediated by common or distinguishable brain systems, respectively. This approach allowed for the first time to test the neural plausibility of appraisal theories of emotion to disentangle the separate contributions of two other-condemning moral emotions (i.e., disgust and anger) in fairness-norm violations (Chapman & Anderson, 2013; Gan et al., 2024; Mullen, 2007; Rozin et al., 2009).

### 4.1. Brain systems mediating unfairness, core disgust, and anger provocation

#### 4.1.1. Unfairness processing

The identified network for unfairness aligned with networks reported in earlier meta-analyses (Feng et al., 2015; Gabay et al., 2014), including bilateral AI, MI, ACC, MCC, dLPFC, vLPFC, dMPFC, OFC, as well as left dorsal striatum and left primary motor cortex. This extensive network may reflect the multifaceted nature of processing unfairness which involves several cognitive and affective processes that simultaneously engage when confronted with unfair offers in the UG. For example, the activation of AI has been shown to reflect negative emotional responses to unfairness (Corradi-Dell’Acqua et al., 2016; Halko, Hlushchuk, Hari, & Schürmann, 2009; Li et al., 2024; McAuliffe et al., 2017; Sanfey et al., 2003), which are thought to play a major role in costly punishment (e.g., rejection) (Feng et al., 2015; Sanfey et al., 2003); although a different interpretation suggests that its activation encodes deviation from norm expectations (Civai, Crescentini, Rustichini, & Rumiati, 2012; Corradi-Dell’Acqua, Civai, Rumiati, & Fink, 2013; Xiang, Lohrenz, & Montague, 2013), which may also drive the emotional response and the decision to reject (Vavra et al., 2017). Despite being less well-studied than AI, MI has been associated with negative subjective emotional experience and may integrate the interoceptive and autonomous signals with the subjective affective experience (Craig, 2009; Ferraro et al., 2022; Mani & Amber, 2015; Mason et al., 2007; Zhou et al., 2020). This region may play a central role in encoding fairness and equality (Lois et al., 2020), such that increased engagement of this region has been associated with inequality-related negative emotional responses (Wright, Symmonds, Fleming, & Dolan, 2011). The ACC has been involved in the detection and resolution of conflict (Chang, Smith, Dufwenberg, & Sanfey, 2011; Fehr & Krajbich, 2014; Feng et al., 2015) and, in the context of punishing fairness norm violations, is considered to reflect the conflict between restoring norms and self-interest (Baumgartner, Knoch, Hotz, Eisenegger, & Fehr, 2011). Apart from the ACC, prefrontal regions including the dLPFC (Baumgartner et al., 2011; Buckholtz & Marois, 2012; Knoch, Pascual-Leone, Meyer, Treyer, & Fehr, 2006; Montague & Lohrenz, 2007; Spitzer, Fischbacher, Herrnberger, Grön, & Fehr, 2007), vLPFC and dMPFC (Feng et al., 2015) as well as OFC (Montague & Lohrenz, 2007; Spitzer et al., 2007), play crucial roles in enforcing norm-related behavior and maintaining fairness. Moreover, the MCC has been implicated in processing social information, especially when predicting and monitoring the outcomes of decisions during social interactions (Apps, Lockwood, & Balsters, 2013). In line with earlier studies (Boccadoro et al., 2021; Feng et al., 2015), we found recruitment of the dorsal striatum in response to unfairness, which, given the role of this region in reward-related anticipation and decision-making initiation (Balleine, Delgado, & Hikosaka, 2007; de Quervain et al., 2004; O’Doherty et al., 2004) as well as in the motivation to punish norm violators (Aoki, Yomogida, & Matsumoto, 2015), might indicate that restoring norm by punishing unfairness might be rewarding enough to override self-interest. In addition, the primary motor cortex has also been demonstrated to be involved in fairness-related decision-making processes (Feng et al., 2019; Gabay et al., 2014).

On the network level, the salience, frontoparietal, DMN, dorsal attention, and limbic networks comprised significant voxels in the unfairness map. Particularly, the first three networks, which together form the ‘triple’ network (V. Menon, 2011), contributed more to the unfairness map than the latter two. The triple network has been considered essential for a variety of cognitive and emotional processes (Hidalgo-Lopez, Engman, Poromaa, Gingnell, & Pletzer, 2023; B. Menon, 2019). Punishing norm-deviant behavior recruited the triple network (Bellucci et al., 2020), which might suggest the salient, motivational, and cognitive appraisal processes underlying the punishment of immoral behavior (Sevinc, Gurvit, & Spreng, 2017). Meta-analytic behavioral decoding based on the Neurosynth database revealed terms primarily associated with norm violations (e.g., error, conflict, stop, inhibition), decision-making (e.g., decision, choice), and salience (e.g., salience, negative feedback, risk taking, losses), which was mirrored by the Brainmap-based decoding with terms mainly related to cognition (e.g., attention, reasoning, social cognition), emotion (e.g., negative, reward, disgust, anger) and action responses (e.g., inhibition, execution, punishment/loss). Finally, echoing previous studies (Crockett, Clark, Lieberman, et al., 2010; Crockett et al., 2008), the involvement of serotonin may point to the affective sources of the ‘irrational’ costly punishment behavior.

Together, these results provide initial support for the notion that emotional responses can play a key role in human decision-making (De Martino, Kumaran, Seymour, & Dolan, 2006) and may moreover reflect that decision-making underlying the punishment choice begins during the confrontation with the unfair offer rather than during the separate actual decision period (i.e., reject the proposer’s offer) (for similar findings see also Boccadoro et al., 2021).

#### 4.1.2. Core disgust processing

In line with one recent meta-analysis on visual core disgust processing (Gan et al., 2022), the multimodal core disgust stimuli yielded robust activation in the bilateral amygdala and adjacent globus pallidus, bilateral AI and MI, a set of prefrontal and orbitofrontal regions (i.e., dMPFC, vMPFC, dLPFC, vLPFC, and OFC) as well as parietal and temporo-occipital regions, which may reflect that direct exposure to potentially poisonous and harmful stimuli engages the evolutionary defensive-avoidance circuits involved in early threat and salience detection and the initiation of behavioral defensive responses. The activation of the AI during core disgust processing has been extensively documented in earlier research (Chapman & Anderson, 2012; Vytal & Hamann, 2010; Wicker et al., 2003), which may reflect coding of salience during emotional processing (Uddin, 2015). In the field of disgust, the MI, while not receiving as much attention as the AI, has also been demonstrated to decode subjective disgust experience induced by core disgust stimuli (Corradi-Dell’Acqua et al., 2016; Gan et al., 2024). Previous studies have extensively reported the activity of the amygdala towards aversive signals (Alexandra Kredlow, Fenster, Laurent, Ressler, & Phelps, 2022; Hagihara et al., 2021; Holley & Fox, 2022; Zald, 2003), such that meta-analytic data showed a higher probability of amygdala activation for stimuli signaling potential threats, including fear and disgust inducing stimuli, as compared to non-threatening stimuli (Costafreda, Brammer, David, & Fu, 2008), indicating a critical role in the detection of salient and in particular threat-related stimuli (Hagihara et al., 2021; Mihov et al., 2013). The amygdala receives dense projections from the visual cortex and via projections with cerebellar regions and the globus pallidus initiates motor and physiological defensive responses towards threatening stimuli (Giovanniello et al., 2020; Lindquist, 2020; Salih et al., 2009; Sambataro et al., 2006). Those responses are strongly modulated by the prefrontal cortex which additionally exhibits dense connections with parietal and temporo-occipital regions (Cechetto & Topolovec, 2002). The identified prefrontal regions play a key role in both, emotional awareness and executive control, including attention and initiation of context-appropriate responses (Alexandra Kredlow et al., 2022; LeDoux, 2014; LeDoux & Pine, 2016; Rozzi & Fogassi, 2017).

On the network level, apart from the frontoparietal network the other 6 large-scale cerebral networks contributed to core disgust processing, which also confirmed the multifaceted evolutionary defensive-avoidance responses induced by core disgust stimuli. For example, the visual cortex has been shown to encode emotional contents (Kragel, Reddan, LaBar, & Wager, 2019), the somatomotor network is responsible for initiating motor behavior (Gordon et al., 2023), and the dorsal attention network encodes and maintains preparatory signals and modulates the activity of the visual and somatomotor regions (Alves, Forkel, Corbetta, & Thiebaut de Schotten, 2022). Additionally, the DMN has been posited to construct disgust experience (Satpute & Lindquist, 2019). The findings not only underscore that subjective emotional experiences are encoded in distributed networks (see also e.g., Gan et al., 2024; Zhou et al., 2021), but also align with the meta-analytic functional characterization in the present study. The Neurosynth decoding revealed terms mainly related to threat-, avoidance-, and salience-related processes (e.g., disgust, suffering, aversive, salient, threatening) and subjective experience (e.g., elicited, inducing, experience, subjective), which was flanked by Brainmap decoding with terms mostly associated with salience (e.g., attention, negative, disgust) and action responses (e.g., inhibition, execution, preparation). Of note, visual-related terms were also evident, which could be driven by the majority of papers included in the core disgust meta-analysis were from the visual modality; however, the behavioral characterization based on both databases also captured terms outside of the visual modality, such as olfaction, gustation, eating and food, corroborating the multimodal nature of the core disgust network. Given the role of serotonin in aversive processing (Crockett, Clark, Apergis-Schoute, Morein-Zamir, & Robbins, 2012; Soubrié, 1986), its engagement in the core disgust network provided complementary evidence to the salient nature of disgust (see also Vicario et al., 2017).

#### 4.1.3. Anger provocation processing

Consistent with the anger-provocation network determined in a prior meta-analysis (Sorella et al., 2021), we found robust activation of the AI, MI, and vLPFC in response to anger-provoking stimuli, which is thought to be related to the subjective experience of anger (Sorella et al., 2021).

The present meta-analysis moreover revealed that the left thalamus, as well as the right MCC, dPMFC, and dorsal striatum were activated by anger provocation. The thalamus has been shown to facilitate salience-related processes (e.g., autonomic arousal, activation of the stress response) upon anger provocation (Alia-Klein et al., 2020; Arshad & Bacha, 2022), the activity of MCC covaried significantly with provocation intensity (Lotze, Veit, Anders, & Birbaumer, 2007; Weidler et al., 2019), and the dMPFC may capture self-referential anger experiences during experimentally elicited angry feelings (Alia-Klein et al., 2020). The activation of the dorsal striatum is also linked to anger experience (Alia-Klein et al., 2020), which could be due to its engagement in the coding of provoking stimuli (Fabiansson, Denson, Moulds, Grisham, & Schira, 2012; Zeki & Romaya, 2008).

On the network level, we observed the involvement of the somatomotor, salience, and frontoparietal networks in response to anger provocation, which dovetailed with a recent meta-analysis on subjective anger experience (Klaus & Schutter, 2021). Besides, the DMN was also implicated in anger provocation, which could be owing to its important role in constructing anger experience (Satpute & Lindquist, 2019). Meta-analytic behavioral decoding from the Neurosynth database showed terms mainly related to negative and salient (e.g., pain, painful, noxious, aversive, unpleasant), subjective experience (e.g., elicit, induced, experiencing, subjective) as well as motor-related (e.g., inhibition) processes, which largely echoed by the Brainmap-based decoding with terms primarily associated with salience (e.g., attention, pain, negative) and action responses (e.g., execution, inhibition, preparation). Among them, pain-related terms are prominent across two databases. Indeed, abundant evidence suggests that the current experience of anger feeling facilitates pain (Adachi, Yamada, Fujino, Enomoto, & Shibata, 2022; Gatchel, Peng, Peters, Fuchs, & Turk, 2007; Gilam et al., 2024; Potegal & Nordman, 2023; Yarns, Cassidy, & Jimenez, 2022), which is particularly evident in situations such as behavioral experiments where the participants tend to suppress the experience or expression of anger (i.e., anger inhibition) (Burns, Quartana, & Bruehl, 2008). Besides, the spatial overlap with the serotonin system provided additional support for the biological plausibility of the anger provocation network observed (Crockett et al., 2012; Passamonti et al., 2012; Soubrié, 1986).

### 4.2. Common and distinct neural correlates between unfairness, core disgust, and anger provocation

#### 4.2.1. Common regions between domains

The bilateral AI and MI were commonly activated for unfairness and core disgust as well as unfairness and anger provocation indicating a key node for the interaction between unfairness and disgust or anger, respectively. The AI and MI have been demonstrated to play a key role in the punishment of norm violations, which may indicate the negative emotional reaction (e.g., disgust and anger in this case) of the punisher to the harm of a norm transgression, thus contributing to their behavioral response (Bellucci et al., 2020; Spitzer et al., 2007; Zinchenko & Arsalidou, 2018). The shared engagement of the AI and MI in two domains was also supported by the evidence showing that the insula represents the shared neural basis for core disgust and moral disgust (e.g., unfairness), which may suggest that the mechanism that initially evolved to process core disgust (e.g., potentially contaminated and infectious stimuli) extends into the sociomoral domain (Vicario et al., 2017). The bilateral vLPFC and OFC were further recruited for both unfairness and core disgust, which may also reflect the negative emotional response induced by aversive stimuli (e.g., unfair offers) (Lois et al., 2020; Moll, de Oliveira-Souza, Bramati, & Grafman, 2002; Rolls, 2023).

On the network level, the salience network and DMN exhibited convergent involvement in unfairness and core disgust as well as in unfairness and anger provocation. Similar results were also reflected by the consensus connectivity profiles which revealed the bilateral AI from the conjunction of unfairness and core disgust as well as from the conjunction of unfairness and anger provocation mainly engaged the salience network and DMN. The salience network plays a central role in the detection of behaviourally relevant or salient stimuli (V. Menon & Uddin, 2010; Uddin, 2015). For example, the salience network was recruited when detecting information related to morality (Sevinc et al., 2017), and its dysfunction could contribute to a broad range of cognitive and affective deficits (Schimmelpfennig, Topczewski, Zajkowski, & Jankowiak-Siuda, 2023). A previous meta-analysis also observed common activation of the DMN for both moral cognition and moral emotions (Sevinc & Spreng, 2014), and its engagement in moral tasks might reflect a role of emotion in moral judgment (Greene, Nystrom, Engell, Darley, & Cohen, 2004; Greene, Sommerville, Nystrom, Darley, & Cohen, 2001; Harrison et al., 2008).

Consistent with the notion that serotonergic mechanisms play a role in both core and moral disgust (Vicario et al., 2017), the hypothesis-driven exploratory neurotransmitter analysis confirmed a shared serotonergic basis for both unfairness and core disgust as well as unfairness and anger, suggesting a vital role of serotonin in mediating affect in moral judgment and in turn driving the punishment behavior (Crockett, Clark, Lieberman, et al., 2010; Hallsson et al., 2018; Vicario et al., 2017). Evidence from the meta-analytic decoding perspective further substantiates the above findings. Interestingly, the results of meta-analytic behavioral decoding for the conjunction of unfairness and disgust highly resembled those for the conjunction of unfairness and anger across both the Brainmap and Neurosynth databases, which may indicate that disgust and anger share a common sociomoral core. Specifically, both conjunction maps captured salience (e.g., attention, negative) and action responses (e.g., inhibition, execution) based on the Brainmap database, as well as more nuanced and granular terms revealed by the Neurosynth database such as the need for engagement (e.g., stop, error, stop signal, signaling), salience (e.g., salience, risk, risk taking, losses), action (e.g., inhibition, conflict) and decision-making (e.g., decision, choice), indicating the key role of both moral emotions in the identification of norm-violating behavior and initiating protective responses.

Core disgust and anger provocation showed convergent activation in the left AI and MI, which on the network level pointed to the salience network, reflecting domain-general salience processing in response to negative emotional stimuli (Corradi-Dell’Acqua et al., 2016) despite both emotions being different in many ways (e.g., physiological responses, action tendencies and facial expressions) (Chapman & Anderson, 2013; Russell & Giner-Sorolla, 2013). Such observations were mirrored by a shared serotonin mechanism for core disgust and anger provocation, indicating the aversive nature of both emotions (Dayan & Huys, 2008). Parallel to the consensus connectivity profiles for the AI from the conjunction of unfairness and core disgust as well as unfairness and anger provocation, we identified a consensus connectivity network for the left MI from the conjunction of core disgust and anger provocation encompassing the salience network and DMN. The DMN has been assumed to shape current affective experiences (e.g., disgust, anger) by utilizing autobiographical experiences and knowledge (Satpute & Lindquist, 2019). However, the consensus connectivity map of the left MI also coupled with regions associated with motor planning (e.g., postcentral gyrus, inferior parietal lobule, supramarginal gyrus) (Andersen, 2011; Buxbaum & Randerath, 2018; Holmes et al., 2024), suggesting that motor-related processes are vital for both subjective emotional experiences (Braine & Georges, 2023). The above findings fit well with the meta-analytic behavioral decoding results of the conjunction of core disgust and anger provocation, which revealed terms overwhelmingly dominated by negative affect and salience (e.g., suffering, aversive, affective, unpleasant, threat, salience, arousal), subjective experience (e.g., experienced, elicited, inducing, subjective).

#### 4.2.2. Domain-specific regions

With respect to differences between domains, regions that showed stronger activity for unfairness than core disgust included the bilateral ACC, MCC, dMPFC, dLPFC, OFC, AI, and MI, as well as right vLPFC and left dorsal striatum, which was remarkably similar to the results of the higher activity during unfairness compared to anger provocation. These findings may suggest a conflict resolution model that requires a higher burden of social cognitive processing in the context of norm violations, relative to core disgust or anger provocation stimuli (i.e., domain-specific unfairness processing). When exposed to unfair treatment, the activation of each of these brain regions subserves specific functions. The AI and MI may reflect negative emotional reactions elicited by norm violations, representing a driving force to punish the transgressors (Feng et al., 2015; Lois et al., 2020; McAuliffe et al., 2017; Zinchenko & Arsalidou, 2018). However, the punishment motivation in enforcing fairness norms entails a cost, which in turn conflicts with economic self-interest. Resolving such conflict requires behavioral control. The ACC is associated with the detection of conflict (Greene et al., 2004; McAuliffe et al., 2017), and then it activates higher cognitive control circuits to resolve the conflict, including the dMPFC that is related to the assessment of the norm violator’s intentions (Krueger & Hoffman, 2016), the vLPFC that down- regulates activity in regions (i.e., insula) suppressing negative affect to enable the rational decision-making (Tabibnia, Satpute, & Lieberman, 2008), the dLPFC and OFC that comply with social norms by suppressing self-interest (Boccadoro et al., 2021; Huebner et al., 2009; Spitzer et al., 2007), and the MCC that regulates cognitive control to guide the punishment decisions (Boccadoro et al., 2021). Finally, the activation in the dorsal striatum may reflect the anticipated reward experience from punishing norm defectors, which is more rewarding than maximizing self-interest (Boccadoro et al., 2021; Crockett, Clark, Lieberman, et al., 2010; de Quervain et al., 2004; Hallsson et al., 2018). Network-level evidence further corroborates the above processes, with the frontoparietal network and DMN preferentially engaging in the evaluation of fairness norm violations and the salience network also showing a high degree of involvement. The three networks have been well-documented in social punishment models (Bellucci et al., 2020; Krueger & Hoffman, 2016). This explanation is also supported by the consensus connectivity profiles of the bilateral AI from the contrast of unfairness > core disgust as well as unfairness > anger provocation, which converged at the salience network, DMN, and frontoparietal network. Collectively, being exposed to unfair treatment triggers multiple affective and cognitive processes that interact with one another to enforce fairness norms through costly punishment.

Hyperactivation for core disgust compared with unfairness encompassed bilateral amygdala and adjacent globus pallidus, parahippocampal gyrus, visual occipital (e.g., fusiform gyrus, inferior and middle occipital gyrus), and motor-related (postcentral gyrus, inferior parietal lobule) regions, which resemble the network that showed a stronger engagement during core disgust as compared to anger provocation. These findings may suggest a neurobiological model for domain-specific core disgust processing. The engagement of visual regions represents evolutionarily hard-wired mechanisms for the automatic detection of threatening stimuli (Bradley et al., 2003; Pessoa & Adolphs, 2010). The amygdala serves as the hub for evaluating the biological significance and guiding survival-relevant responses (Holley & Fox, 2022; Pessoa & Adolphs, 2010), while the parahippocampal gyrus may further qualify the contextual relevance of the stimuli (Tao et al., 2021). Moreover, the survival-relevant reaction requires fast withdrawal behavior (Rolls, 1999, 2000), which is potentially mediated by inferior parietal and pallidal regions. The inferior parietal lobule is a key location for motor planning (Buxbaum & Randerath, 2018), receiving inputs from both visual and somatosensory areas, and is therefore an optimal locus for integrating these modalities (Andersen, 2011). Finally, the globus pallidus is associated with implementing rapid motor responses to emotional stimuli (Sambataro et al., 2006) and via connections with the amygdala regulates reactions to aversive stimuli (Giovanniello et al., 2020). On the network level, the visual, dorsal attention, and limbic networks exhibited the largest contributions to domain-specific core disgust processing, and the somatomotor network made a modest contribution as well, further confirming the proposed neurobiological model. Combined, these regions may constitute the core defensive-avoidance circuits underlying domain-specific core disgust processing.

Regions exhibited stronger activation for anger provocation as compared to unfairness converged in the left hemisphere (including the thalamus, primary somatosensory cortex, AI, and MI). The consensus connectivity profile of the left AI provides further important insights into the mechanisms underlying this left-lateralized hyperactivation during anger provocation, such that the left AI coupled mainly with the salience network; besides, regions associated with motor processes such as the primary motor cortex and inferior parietal lobule were prominently connected. Moreover, the left AI also coupled with the thalamus. These results mutually confirmed each other and may indicate that motor-related areas and the thalamus represent distinctive signatures of discriminating anger provocation from unfairness. The hyperactivation of anger provocation in comparison to core disgust comprised the bilateral AI and MI, left thalamus, right dorsal striatum, right MCC, and dMPFC, which mainly mirrored the results of the meta-analytic anger provocation network and might imply the contributions of these regions in constructing subjective anger experience despite both emotions being high arousal in nature. On the consensus connectivity level, the bilateral AI coupled with regions mainly from the salience network, with smaller contributions from the frontoparietal network and DMN, which largely echoed the network-level results. More interestingly, the thalamus was also robustly involved in the consensus connectivity profile, resonating with the consensus connectivity profile of the left AI of anger provocation larger than unfairness. Overall, the thalamus represents a unique representation of domain-specific anger provocation processing.

### 4.3. Methodological considerations

Previous studies have repetitively reported the multifaceted behavioral functions of the AI, such as salience (Uddin, 2015), emotional awareness (X. Gu, Hof, Friston, & Fan, 2013), interoception (Craig, 2009), and social emotions (Lamm & Singer, 2010). In this study, the peak of each insula region from the conjunction and contrast analyses mostly centered on the anterior part. To avoid rather ‘coarse’ inferences of the potential mechanisms underlying each of the AI based mainly on previous studies, we adopted a new method by combining task-based connectivity with task-free connectivity approaches and further derived the consensus connectivity profile from the two approaches (e.g., Bellucci et al., 2020; Bore et al., 2024). As detailed in the previous section, the consensus connectivity profiles empowered us to dissect differential contributions of the specific insula regions on the network and behavioral level. On the one hand, all of the consensus connectivity maps across different conditions included the salience network and DMN, which might reflect the domain-general role of the two networks in coupling with the AI; on the other hand, there were exceptions such that the two networks were more distributed in conjunction conditions in comparison to contrast conditions, while the AI required the robust engagement of motor regions for anger provocation relative to unfairness but not vice versa and further coupled with thalamus for domain-specific anger provocation processing.

### 4.4. Biologically-informed model

Appraisal models of emotion have long posited that reactions towards unfairness are largely shaped by strongly negative emotional responses (especially two emotions that involve the condemning of others, that is, anger and disgust) (Chapman & Anderson, 2013; Gan et al., 2024; Mullen, 2007; Rozin et al., 2009). Previous studies confirmed shared neural representations for unfairness and core disgust (Corradi-Dell’Acqua et al., 2016; Gan et al., 2024), however, the extent to which the appraisals of unfairness are distinct from those of core disgust has not been investigated. Moreover, few studies have systematically examined common and separable appraisals of unfairness and anger provocation at the neural level. In this study, we employed neuroimaging-based meta-analysis aiming at providing neurobiological evidence to answer these questions.

We found that a moral event such as an unfair monetary offer triggers brain systems involved in cognitive appraisals that partly overlap with those triggered by core disgust as well as anger provocation. As shown in Figure 8, the AI and MI represent the shared neurofunctional bases of unfairness and core disgust appraisals. We further demonstrate that both regions also represent the shared neural substrates of unfairness and anger provocation appraisals, while vLPFC and OFC moreover serve as common neural bases of unfairness and core disgust appraisals. From an evolutionary viewpoint, these results may indicate that some of the appraisal systems mediating the strong emotional responses towards threatening or provoking (core disgust or anger provocation) stimuli were further preadapted into the sociomoral domain and may serve to maintain fairness norms in social groups. These results were further confirmed by a shared serotonergic basis for unfairness and core disgust (also compatible with Vicario et al., 2017) as well as anger provocation. The above findings are consistent with the view that emotion plays a key role in moral cognition (Bargh et al., 2012; Chapman & Anderson, 2013; Greene & Haidt, 2002; Haidt, 2001; Prinz, 2006; Russell & Giner-Sorolla, 2013; Sharvit, Lin, Vuilleumier, & Corradi-Dell’Acqua, 2020; Tybur et al., 2013). The engagement of disgust and anger in the evaluation of moral transgressions may indicate the two emotions share a common sociomoral core such that both of them function to guard our social body against immoral behaviors and motivate the enforcement of moral norms via punishment and condemnation etc (Andersson et al., 2024; Molho et al., 2017; Molho et al., 2020; Skitka et al., 2008), while the co-occurrence of anger and disgust in response to moral violations may lead to harsher punitive actions (Olatunji & Puncochar, 2014) despite the personal cost, deviating from the rational ‘Homo economicus’ model of behavior. Apart from shared neural representations, we also observed other brain regions involved in domain-specific processing, which may point to separable appraisals subserving specific facets of respective domains.

**Figure 8.**
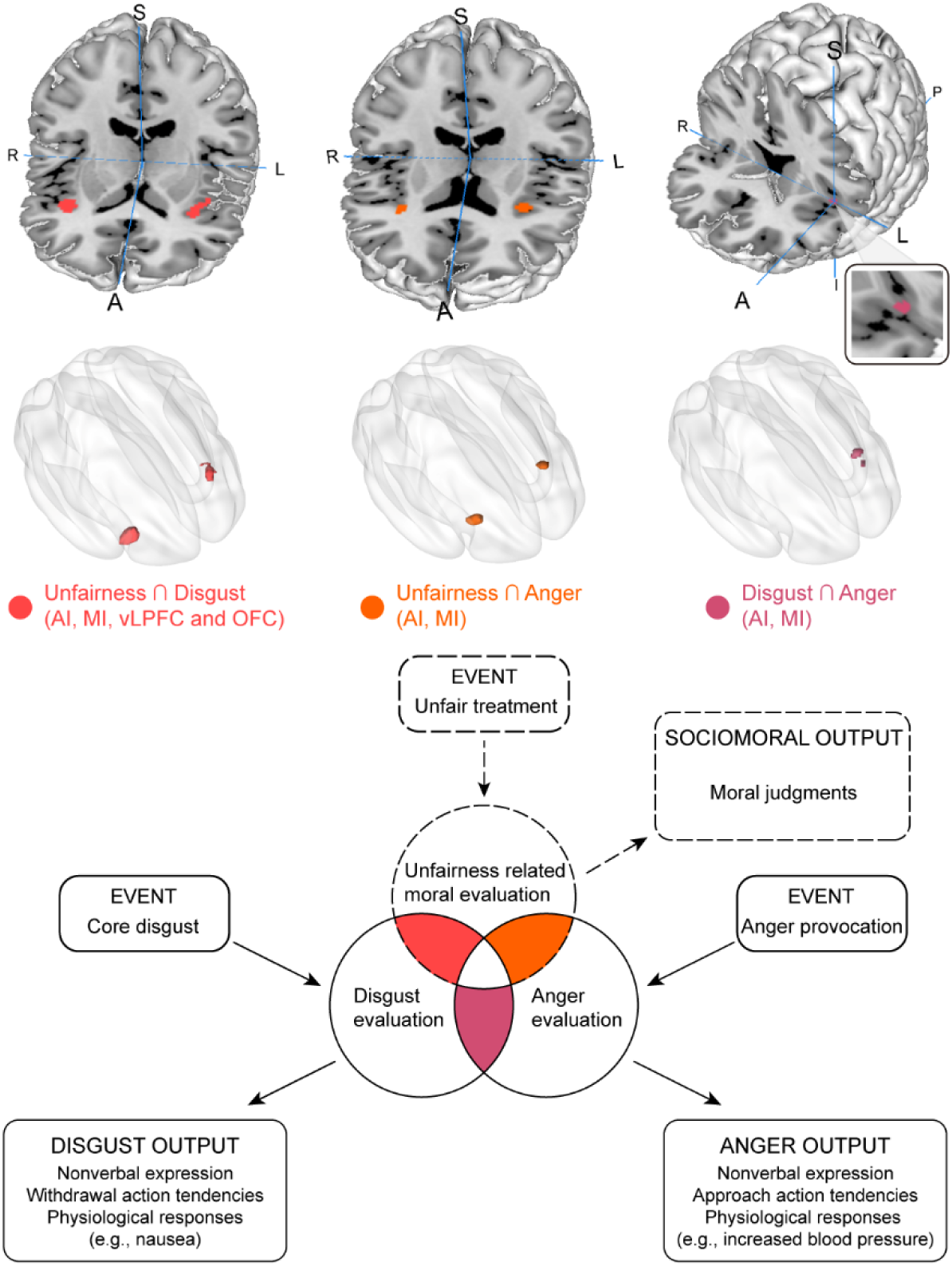
Neural systems underlying the shared appraisals between unfairness and disgust (AI, MI, OFC, and vLPFC), unfairness and anger (AI and MI), as well as between disgust and anger (AI and MI).

### 4.5. The rejection network

The left dorsal striatum robustly and specifically activated during rejecting as compared to accepting unfair offers, which replicated previous meta-analyses on the same topic (Boccadoro et al., 2021; Gabay et al., 2014). The dorsal striatum has been implicated in the punishment of an unfair proposer (Aoki et al., 2015; Crockett et al., 2013; Crockett, Clark, Lieberman, et al., 2010; de Quervain et al., 2004), suggesting that sanctioning norm transgressors (i.e., rejecting an unfair offer) is inherently rewarding. On the consensus connectivity level, the left striatum coupled with regions from the salience network, DMN, and frontoparietal network as well as the somatomotor network. The first three networks have been demonstrated to play a vital role in social punishment models (Bellucci et al., 2020; Krueger & Hoffman, 2016) and the somatomotor network may additionally be involved in the implementation of punishment behaviors. Unexpectedly, we did not obtain any overlap between rejection and core disgust as well as anger provocation.

After relaxing the threshold, we observed more distributed regions involved in the rejection response, including the left dorsal striatum, ACC, MCC, and amygdala as well as bilateral AI, MI, and parietal motor regions, which may suggest a model of motivational conflict during the rejection decision-making process, with possible influence from the reward and affective systems.

The following conjunction analyses revealed that the amygdala was commonly engaged during rejection and core disgust, which may suggest a shared neuroaffective basis. However, these findings were exploratory due to the limited number of experiments (i.e., 14) included in the rejection domain as well as the lenient threshold, thus warranting much more consideration in future research.

### 4.6. Limitations

The present study has the following limitations. First, in this study, we focused on violating fairness norms that are deemed to be central to the moral domain (Chapman et al., 2009; Henrich et al., 2004; Huebner et al., 2009; Sheskin, 2017; Tomasello, 2016), nevertheless, the evidence provided by the current study may not generalize to all form of moral behavior (Graham et al., 2013; Haidt, 2007; Haidt & Graham, 2007; Hopp et al., 2023). Second, we specifically concentrated on second-party punishment of unfairness as a way to reduce heterogeneities and biases and we did not further account for other forms of fairness such as third-party punishment of norm violations (McAuliffe et al., 2017), procedural justice (Lind & Tyler, 1988), and retributive justice (Carlsmith & Darley, 2008), future studies may test whether disgust or anger plays a role in the decision-making about these forms of fairness, as Mullen highlighted affect as a relevant component of any theory of justice (Mullen, 2007). Thirdly, given that two other-condemning moral emotions (i.e., disgust and anger) have been extensively implicated in previous empirical studies and have been hypothesized to be particularly relevant to the perception of unfairness (Mullen, 2007; Olatunji & Puncochar, 2014; Rozin et al., 2009), this study sought to provide neurobiological evidence of the underlying mechanisms of the two emotions in fairness norm decision-making. Nonetheless, there are other kinds of moral emotions (e.g., contempt, guilt, shame, and compassion) (Rozin et al., 1999; Urbanska, McKeown, & Taylor, 2019; Yu et al., 2020; X. Zheng et al., 2024), investigating the role of these emotions in enforcing fairness norms is an exciting avenue for future work.

### 4.7. Conclusions

The present study employed coordinate-based neuroimaging meta-analyses to determine common and distinct brain systems engaged in unfairness and disgust as well as in unfairness and anger and further combined multimodal analytic approaches to provide a more comprehensive dissection of the role of the two other-condemning moral emotions when witnessing fairness norm violations. The confrontation with unfair offers evoked robust activation in regions involved in multiple cognitive and affective processes that interact to enforce fairness norms via costly punishment, exposure to disgust-inducing stimuli engaged the evolutionary core defensive-avoidance circuits involved in early salience and threat detection and the initiation of defensive motor responses, while anger provoking stimuli elicited robust activity in a network including regions primarily engaged in aversive and salience processes. The AI and MI were commonly activated for unfairness and core disgust as well as for unfairness and anger provocation, the vLPFC and OFC were further commonly engaged during unfairness and core disgust, suggesting an involvement of these regions across unfairness and disgust as well as unfairness and anger. Together, this study for the first time validated the neural plausibility of utilizing the appraisal theories of emotion to explain the role of emotions in fairness norm violations and provided the neurobiological evidence for the shared and separable appraisals between unfairness and two other-condemning moral emotions (i.e., disgust and anger).

## Declaration of Competing Interest

We declare no conflict of interest.

## Acknowledgements

The work was partly supported by the China MOST2030 Brain Project (grant no. 2022ZD0208500), the National Natural Science Foundation of China (grants no. 32250610208 and 82271583), and a start-up funding from The University of Hong Kong.

## Supplemental Materials

Supplemental data associated with this article can be found in the online version at

## Notes

### Competing Interest Statement

The authors have declared no competing interest.

