## Supplemental Materials for "Does Unfairness Evoke Anger or Disgust? A Quantitative Neurofunctional Dissection Based on 25 Years of Neuroimaging"

### Supplemental Results

#### Behavioral characterization results based on the BrainMap database

We adopted another data-driven approach (i.e., behavioral analysis) using the BrainMap database to complement the Neurosynth functional characterization. The whole-brain patterns of unfairness from single dataset analysis were characterized by a pool of different subdomains covering all five domains, such as cognition (e.g., attention, memory, reasoning, language, and social cognition), emotion (e.g., negative, reward, disgust, intensity, and anger), action (e.g., execution, inhibition, and preparation), perception (e.g., somesthesis, pain, and vision), and interoception (e.g., sexuality) (Figure S8a). Analogously, the whole-brain map of core disgust from single dataset analysis revealed subdomains belonging to all five domains, including cognition (e.g., attention, memory, reasoning, and language), emotion (e.g., negative, disgust, and reward), action (e.g., inhibition, execution, and preparation), perception (e.g., vision and pain), interoception (e.g., sexuality) (Figure S8b). Moreover, the whole-brain map of anger provocation from single dataset analysis also showed subdomains across all five domains: cognition (e.g., attention, memory, reasoning, and language), emotion (e.g., negative and reward), action (e.g., execution, inhibition, and preparation), perception (e.g., somesthesis, pain, and vision), and interoception (e.g., sexuality) (Figure S8c).

The whole-brain map of unfairness ∩ core disgust from conjunction analysis was mainly associated with subdomains of four domains: cognition (e.g., attention, memory, reasoning, and language), emotion (e.g., negative and reward), action (e.g., inhibition and execution), and perception (e.g., somesthesis, pain, and vision) (Figure S8d). Likewise, the whole-brain map of unfairness ∩ anger provocation from conjunction analysis was primarily characterized by subdomains covering four domains: cognition (e.g., attention, memory, reasoning, and language), emotion (e.g., negative and reward), action (e.g., execution and inhibition), and perception (e.g., pain and vision) (Figure S8e). Note that no term obtained z-score larger than 3 for the whole-brain map of core disgust ∩ anger provocation from conjunction analysis.

#### Lenient thresholding for the single dataset analysis of rejection and exploratory conjunction and contrast analyses

We used lenient thresholding (voxel-level uncorrected $p < 0.001$) for the single dataset analysis aiming to provide some trend of the rejection response. The results showed the activation of the left putamen, MCC, ACC, amygdala, parahippocampal gyrus, right cuneus, cerebellar regions, and a bilateral network including the MI, AI, claustrum, postcentral gyrus, and inferior parietal gyrus in the rejection of unfair offers (Table S15, Figure S11a). Next, a series of conjunction and contrast analyses were conducted to find whether there were common and separable neural substrates between this ‘lenient’ rejection network and core disgust or anger provocation. The conjunction analysis for rejection and core disgust revealed that the left amygdala extending into the adjacent parahippocampal gyrus was activated during both processes (Table S16, Figure S11b), and the contrast analyses between the two domains largely resembled the previous one (i.e., Figure S9b) with the exception that the bilateral AI, MI, and claustrum were further implicated in the contrast of rejection > core disgust (Table S17, Figure S11b). Moreover, the conjunction analysis did not receive any overlap between rejection and anger provocation. The contrast analyses revealed stronger recruitment of the left putamen for rejection in comparison to anger provocation, whereas anger provocation showed a stronger involvement of the right AI, MI, and claustrum compared with rejection (Table S18, Figure S11b). However, all these results are suggestive and preliminary because of the low number of experiments included in the rejection domain, more research is needed to ensure robust inferences.

Table S1

Characteristics of the included studies with reference.

| Authors | Gender Ratio (f/m) | Age | Contrasts | Coordinates | Analyze software |
|---|---|---|---|---|---|
| *Unfairness studies* | | | | | |
| (Cortes et al., 2018) | 8/19 | M=40.5; SD=7.92 | Unfair>Fair | TAL | AFNI |
| (Y. Zheng et al., 2017) | 18mix | M=about 22.8; SD=about 1.4 | Unfair>Fair | MNI | SPM8 |
| (Wei et al., 2018) | 25mix | M=about 22.46; SD=about 2.62 | Unfair>Fair | MNI | SPM8 |
| (Zheng et al., 2015) | 18/7 | M=21.44; SD=3.38 | Unequal>Equal | MNI | SPM5 |
| (Guo et al., 2014) | 13/5 | M=21.06; SD=2.10 | Unfair>Fair | MNI | SPM5 |
| (Guo et al., 2013) | 21mix | M=about 22.44; SD=about 3.49 | Unfair>Fair | MNI | SPM5 |
| (Hu et al., 2016) | 13/10 | M=21.22; SD=1.73 | Unfair>Fair | MNI | SPM8 |
| (Lois, Schneider, Kaurin, & Wessa, 2020) | 21/20 | M=45.3; SD=13.8 (20–66) | Very unfair>Fair offers | MNI | SPM12 |
| (Fatfouta, Meshi, Merkl, & Heekeren, 2018) | 8/15 | M=24.35; SD=3.80 | unfair unknown>fair unknown | MNI | FSL |
| (Harlé & Sanfey, 2012) | 23/15 | M=22.4 (young group); M=64.1 (older group) | Unfair>Fair | TAL | Brain Voyager v1.10 |
| (Ouyang et al., 2020) | 15/6 | M=22.62; SD=3.61; (19-34) | Unfair>Fair | MNI | SPM8 |
| (Servaas et al., 2015) | 114mix | M=about 20.8; SD=about 2.0 | Unfair>Fair | MNI | SPM8 |

| | | | | | |
|---|---|---|---|---|---|
| | | (18-25) | | | |
| (Verdejo-Garcia, Verdejo-Román, Albein-Urios, Martínez-González, & Soriano-Mas, 2017) | 1/18 | M=30.84; SD=4.13 | Unfair>Fair | MNI | SPM8 |
| (Kirk, Downar, & Montague, 2011) | 21/19 | M=36.8; SD=10.1 | Controls unfair>Controls fair | MNI | SPM2 |
| (Sanfey, Rilling, Aronson, Nystrom, & Cohen, 2003) | 11/8 | M=21.8; SD=7.8 | Unfair>Fair | TAL | Brain Voyager |
| (Civai, Crescentini, Rustichini, & Rumiati, 2012) | 12/7 | Not report | Unequal>Equal | MNI | SPM8 |
| (Güroğlu, van den Bos, van Dijk, Rombouts, & Crone, 2011) | 32/36 | 10- to 20-year-olds | Unfair>Fair | MNI | SPM5 |
| (Halko, Hlushchuk, Hari, & Schürmann, 2009) | 8/15 | M=29; (22-46) | Unfair>Fair (No-competition) | MNI | SPM2 |
| (Zhou, Wang, Rao, Yang, & Li, 2014) | 15/13 | M=25.07; SD=3.35 | Unfair>Fair | MNI | SPM5 |
| (Yoder & Decety, 2020) | 27mix | M=26.7; (18–54) | Self Unfair>Self Fair | MNI | SPM12 |
| (L. Zheng et al., 2017) | 19/18 | M=about 22.79; SD=about 2.81 | Unfair>Fair | MNI | SPM8 |
| (Farmer, Apps, & Tsakiris, 2016) | 14/4 | M=21.1; SD=2.4 | Unfair>Fair | MNI | SPM8 |
| (Verdejo-García et al., 2015) | 25/19 (Normal weight); 24/12 (Excess weight) | M=15.32; SD=1.69 (Normal weight); M=15.06; SD=1.88 (Excess weight) | Unfair>Fair | MNI | SPM8 |
| (Pan et al., 2022) | 22/7 | M=about 20.3; SD=about 2.1 | Unfair>Fair | MNI | SPM12 |
| (Wang et al., 2019) | 193mix | M=about 19.32; | Unfair>Fair | MNI | SPM12 |

| | | | | | |
|---|---|---|---|---|---|
| | | SD=about 1.38 | (data applied from the corresponding author) | | |
| (Haruno, Kimura, & Frith, 2014) | 33/26 | about 20-year-old | parametric analysis: positive correlation with inequity | MNI | SPM8 |
| (Gradin et al., 2015) | 23mix | M=25.44; SD=5.02 | parametric analysis: positive correlation with inequity | MNI | SPM8 |
| (Kirk et al., 2016) | 50mix | about 30-year-old | parametric analysis: positive correlation with unfair offer sizes (Pre-training) | MNI | SPM8 |
| *Rejection studies* | | | | | |
| (Y. Zheng et al., 2017) | 18mix | M=about 22.8; SD=about 1.4 | Reject unfair>Accept unfair (data applied from the corresponding author) | MNI | SPM8 |
| (Wei et al., 2018) | 25mix | M=about 22.46; SD=about 2.62 | Reject unfair>Accept unfair | MNI | SPM8 |
| (Guo et al., 2013) | 21mix | M=about 22.44; SD=about 3.49 | Reject unfair>Accept unfair | MNI | SPM5 |
| (Ouyang et al., 2020) | 15/6 | M=22.62; SD=3.61; (19-34) | Reject unfair>Accept unfair | MNI | SPM8 |
| (Servaas et al., 2015) | 114mix | M=about 20.8; SD=about 2.0 (18-25) | Reject unfair>Accept unfair | MNI | SPM8 |
| (Verdejo-Garcia et al., 2017) | 1/18 | M=30.84; SD=4.13 | Reject unfair>Accept unfair | MNI | SPM8 |
| (Civai et al., 2012) | 10mix | Not report | Reject>Accept | MNI | SPM8 |
| (L. Zheng et al., 2017) | 19/18 | M=about 22.79; SD=about 2.81 | Reject unfair>Accept unfair | MNI | SPM8 |
| (Verdejo-García et al., 2015) | 25/19 (Normal weight); | M=15.32; SD=1.69 (Normal | Reject unfair>Accept unfair | MNI | SPM8 |

| | | | | | |
|---|---|---|---|---|---|
| | 24/12 (Excess weight) | weight); M=15.06; SD=1.88 (Excess weight) | | | |
| (Corradi-Dell'Acqua, Civai, Rumiati, & Fink, 2013) | 9/14 | M=23.5 (18–35) | Reject>Accept | MNI | SPM8 |
| (Lois et al., 2020) | 21/20 | M=45.3; SD=13.8 (20–66) | Reject>Accept | MNI | SPM12 |
| (Güroğlu, van den Bos, Rombouts, & Crone, 2010) | 13/10 | M=20.4; SD=1.7 | Reject unfair>Accept unfair | MNI | SPM5 |
| (Cheng et al., 2017) | 18mix | M=about 23.8; SD=about 1.9 | Reject>Accept | MNI | SPM8 |
| *Core disgust studies* | | | | | |
| (Schienle et al., 2002) | 12/0 | M=26.3; (21–41) | Disgust>Neutral | MNI | SPM99 |
| (Schienle, Übel, Schöngaßner, Ille, & Scharmüller, 2014) | 34/0 | M=23.9; SD=4 | Disgust>Neutral | MNI | SPM8 |
| (Calder et al., 2007) | 7/5 | M=22; SD=2.4 | Disgusting>Non-foods; Disgusting>Bland; Disgusting>Appetizing | MNI | SPM99 |
| (Pujol et al., 2018) | 15/15 | M=27.9; SD=7.8 (19-45) | Disgusting>Appetizing | MNI | SPM8 |
| (Tettamanti et al., 2012) | 19/0 | M=24.1; SD=5.2 (19–39) | Disgust>Neutral | MNI | SPM5 |
| (Stark et al., 2007) | 32/34 | M=24.7; SD=5.2 (19-44) | Disgust>Neutral; Significant activations for the explorative analysis of the parametric modulator DISGUST | MNI | SPM2 |
| (Jabbi, Bastiaansen, & Keysers, 2008) | 6/6 | Not report | Disgust>Neutral (Taste; Imagination) | MNI | SPM2 |
| (Wicker et al., 2003) | 0/14 | (20-27) | Disgusting odorant>Rest | MNI | SPM99 |

| | | | | | |
|---|---|---|---|---|---|
| (Shapira et al., 2003) | 5/3 | M=38 (34-44) | Disgust>Neutral | TAL | MEDx |
| (Schienle & Scharmüller, 2013) | 34/0 | M=23; SD=3.4 | Disgust>Neutral | MNI | SPM8 |
| (Schäfer, Schienle, & Vaitl, 2005) | 10/10 | M=23.93 (19–32) | Disgust>Neutral (block design; and event-related design) | MNI | SPM2 |
| (Wabnegger, Übel, Suchar, & Schienle, 2018) | 16/0 | M=31.13; SD=12.18 | Disgust>Neutral | MNI | SPM12 |
| (Baumann & Mattingley, 2012) | 15/15 | M=22.2; SD=2.9 (18-30) | Disgust>Neutral | MNI | SPM5 |
| (Schienle, Höfler, Keck, & Wabnegger, 2020) | 5/21 | M=32.65; SD=13.47 | Disgust>Neutral | MNI | SPM12 |
| (Phillips et al., 1998) | 0/6 | M=37 (25-43) | Disgust>Neutral | TAL | ANMR |
| (Phillips et al., 2000) | 7/7 | M=31 (20–48) | Disgust>Neutral | TAL | Other |
| (Schienle, Schäfer, Hermann, & Vaitl, 2009) | 19/0 | M=22.3; SD=2.6 | Disgust>Neutral healthy control-normal weight | MNI | SPM2 |
| | 17/0 | M=25.0; SD=4.7 | Disgust>Neutral healthy control-outweight | | |
| (Wright, He, Shapira, Goodman, & Liu, 2004) | 4/4 | (20–26) | Mutilation>Neutral | TAL | Brain Voyager v4.9.6 |
| | | | Contamination>Neutral | | |
| (Schienle et al., 2006) | 12/0 | (19-41) | Contamination>Neutral | MNI | SPM2 |
| | | | Mutilation>Neutral | | |
| (Benuzzi, Lui, Duzzi, Nichelli, & Porro, 2008) | 15/0 | M=23.5; (19–31) | Disgust>Neutral | MNI | SPM2 |
| (Stark, Schienle, Girod, et al., 2005) | 6/6 | M about 28.2; (20–40) | Disgust>Neutral (nonSM) | MNI | SPM99 |
| | 6/6 | M about 28.2; (20–40) | Disgust>Neutral (SM) | | |
| (Viol et al., 2019) | 17mix | M age about 43.5 | Disgust>Neutral | MNI | SPM12 |
| (Schienle et al., 2004) | 12/0 | M=26.3; SD=6.4 | Disgust>Neutral | MNI | SPM99 |

| | | | | | |
|---|---|---|---|---|---|
| (Schienle, Schäfer, Stark, Walter, & Vaitl, 2005) | 63/0 | M=27.3; SD=8.4 | Disgust>Neutral | MNI | SPM99 |
| (Schäfer, Leutgeb, Reishofer, Ebner, & Schienle, 2009) | 18/0 | M=24.8; SD=2.4 | Disgust>Neutral | MNI | SPM2 |
| (Hermann et al., 2007) | 10/0 | M=27.6; SD=10.7 | Disgust>Neutral | MNI | SPM2 |
| (Schienle, Schäfer, Walter, Stark, & Vaitl, 2005) | 13/0 | M=23.9; SD=6.8 | Disgust>Neutral | MNI | SPM99 |
| (Schienle, Ille, & Wabnegger, 2015) | 11/11 | M=51.8; SD=9.8 | Disgust>Neutral | MNI | SPM12 |
| (Radua et al., 2014) | 28/12 | M=38; SD=11 (19–59) | Disgust>Neutral | MNI | FEAT (part of FSL) |
| (Stark et al., 2004) | 0/24 | M=25.5; SD=2.65 (20–31) | Disgust>Neutral | MNI | SPM99 |
| (Lassalle et al., 2019) | 3/17 | M=24.15; SD=7.57 | Disgust>Neutral | MNI | FEAT (part of FSL) v6 |
| (Karama, Armony, & Beauregard, 2011) | 0/18 | M=about 25.5; SD=about 3.4 (about 21-30) | Disgust>Neutral | MNI | SPM5 |
| (Stark, Schienle, Sarlo, et al., 2005) | 11/4 | M=29.1 (20–41) | Disgust>Neutral | MNI | SPM2 |
| (Phillips et al., 2001) | 2/4 | M=33.8 (24-48) | Aversive>Neutral | TAL | Other |
| (Wittfoth, Pfeiffer, Bohne, Lanfermann, & Wittfoth, 2020) | 8/9 | M=23.47; SD=2.45 | Disgust>Neutral | MNI | SPM12 |
| (Harris & Fiske, 2006) | 10mix | Not report | Disgust>Fixation | TAL | Brain Voyager |
| (Ahn et al., 2014) | 42/41 | M=29.0; SD=11.3 (18–62) | Disgust>Neutral | MNI | SPM8 |
| (Harrison, Gray, Gianaros, & Critchley, 2010) | 7/5 | M=25.9; SD=5.7 | Disgust>Control | MNI | SPM5 |
| (S. Chen et al., 2021) | 15/16 | M=21.34 | Watching-disgust>Watching-neutral | MNI | SPM8 |

| | | | | | |
|---|---|---|---|---|---|
| (Shimamura, Marian, & Haskins, 2013) | 13/7 | M=21.1 (18–33) | Disgust express>Neutral express | MNI | SPM2 |
| (Pitskel, Bolling, Kaiser, Crowley, & Pelphrey, 2011) | 6/9 | M=13.03; SD=2.20 (7–17) | Look-disgust>Look-neutral | TAL | Brain Voyager |
| (Goldin, McRae, Ramel, & Gross, 2008) | 17/0 | M=22.7; SD=3.5 | Watch-disgust>Watch-neutral | TAL | AFNI |
| (Meier et al., 2015) | 0/20 | M=25.3; SD=3.0 (20-31) | Disgusting odor>Pleasant odor | MNI | SPM8 |
| *Anger provocation studies* | | | | | |
| (Repple et al., 2018) | 20/22 | M=24.77; SD=2.8 (Men); M=27.45; SD=9.3 (Women) | High provocation>Low provocation; Aggression after high provocation>Aggression after low provocation (Taylor Aggression Paradigm) | MNI | SPM8 |
| (Chester & DeWall, 2019) | 44/17 | M=18.61; SD=0.84 (18–22) | High provocation>Low provocation (Taylor Aggression Paradigm) | MNI | FSL v 5.0 |
| (Chester et al., 2019) | 37/24 | M=18.98; SD=1.07 (18–22) | Retaliatory>Non-Retaliatory Aggression (High provocation>Low provocation) (Adapted Taylor Aggression Paradigm) | MNI | FSL v 5.0 |
| (Repple et al., 2017) | 0/29 | M=about 23.6; SD=about 3.2 | Aggression after high provocation>Aggression after low provocation; High provocation>Low provocation (Modified Taylor Aggression Paradigm) | MNI | SPM8 |
| (Dambacher et al., 2015) | 0/15 | M=22.33; SD=2.35 | Provocation>No provocation (Taylor Aggression Paradigm) | TAL | Brain Voyager QX |
| (Krämer, Jansma, Tempelmann, & Münte, 2007) | 15mix | M=about 22.9; SD=about 2.2 | High provocation>Low provocation (Taylor Aggression Paradigm) | TAL | Brain Voyager QX |

| | | | | | |
|---|---|---|---|---|---|
| (Chester, Lynam, Milich, & DeWall, 2018) | 38/22 | M=20.28; SD=2.77 (18-30) | Retaliatory aggression>Non-retaliatory aggression (High provocation>Low provocation) (Taylor Aggression Paradigm) | MNI | FSL v 5.0 |
| (Beyer, Münte, Erdmann, & Krämer, 2014) | 30/0 | M=about 23.2; SD=about 2.7 | High provocation>Low provocation (Taylor Aggression Paradigm) | MNI | SPM8 |
| (Buades-Rotger, Beyer, & Krämer, 2017) | 36/0 | M=22; SD=4 | High>Low selection phase (High provocation>Low provocation) (Variant of the Taylor Aggression Paradigm) | MNI | SPM12 |
| (Chester & DeWall, 2016) | 47/22 | M=18.70; SD=0.93 | Retaliatory>Non-retaliatory aggression (High provocation>Low provocation) (Taylor Aggression Paradigm) | MNI | FSL v 5.0 |
| (Emmerling et al., 2016) | 0/15 | M=22.33; SD=2.35 | Retaliation to provoking opponent>Non-retaliation to non-provoking opponent (High provocation>Low provocation) (Taylor Aggression Paradigm) | TAL | Brain Voyager QX |
| (Chester & DeWall, 2018) | 13/11 | M=23.04; SD=2.46 (21–30) | Retaliatory Aggression>Non-Retaliatory Aggression (High provocation>Low provocation) (Taylor Aggression Paradigm) | MNI | FSL v 5.0 |
| (Zheltyakova et al., 2022) | 26/13 | M=24.5; SD=3.6 | High provocation>Low provocation (Taylor aggression paradigm) | MNI | SPM12 |
| (B. Chen et al., 2021) | 16/18 | About 20-year-old | Provocation>Baseline; | MNI | SPM12 |

| | | | | | |
|---|---|---|---|---|---|
| | | | Aggressive responses>Baseline (Point subtraction aggression paradigm) | | |
| (Skibsted et al., 2017) | 11/8 | M=24.6; SD=2.9 (20–31) | Provocation event>Monetary response (High provocation>Low provocation); Aggressive response>Monetary response (High provocation>Low provocation) (Point subtraction aggression paradigm) | MNI | SPM8 |
| (Denson, Pedersen, Ronquillo, & Nandy, 2009) | 12/8 | M=18.68; SD=0.75 | Interpersonal Provocation>Baseline (Interpersonal Provocation procedure) | TAL | Brain Voyager QX |
| (Denson, Ronay, von Hippel, & Schira, 2013) | 0/19 | M=22.58; SD=4.05 | Interpersonal Provocation>Baseline (Interpersonal Provocation procedure) | TAL | Brain Voyager QX |
| (Seok & Cheong, 2019) | 0/13 | M=24.2; SD=2.26 (21–27) | Anger-provoking>Neutral conditions (Anger-provoking films) | TAL | SPM12 |
| (Park, Lee, & Sohn, 2016) | 0/16 | M=50.06; SD=6.10 (31–61) | Anger-provoking>Neutral conditions (Anger-provoking films) | TAL | SPM8 |
| (White, Brislin, Meffert, Sinclair, & Blair, 2013) | 7/13 | M=14.15; SD=2.29 | Positive modulation by punishment level; parametric analysis (Adaptation of Taylor Aggression Paradigm) | MNI | AFNI |
| (White, Brislin, Sinclair, & Blair, 2014) | 9/12 | M=28.1; SD=8.1 (21–49) | Positive modulation by punishment level; parametric analysis (Adaptation of Taylor Aggression Paradigm) | MNI | AFNI |

| | | | | | |
|---|---|---|---|---|---|
| (Weidler et al., 2019) | 0/52 | About 25-year-old | Provocation phase; parametric modulation "provocation intensity" (Taylor Aggression Paradigm) | MNI | SPM12 |
| (Konzok et al., 2022) | 61mix | M=about 23.62; SD=about 3.81 | Positive modulation by selected aggression levels; parametric analysis (Taylor Aggression Paradigm) | MNI | SPM12 |
| (Lotze, Veit, Anders, & Birbaumer, 2007) | 0/14 | M=about 28.6; SD=about 6.5 | Receiving aversive stimuli; parametric analysis (Taylor Aggression Paradigm) | MNI | SPM2 |

Table S2

Brain regions activated by conjunction analysis between anger provocation and core disgust in healthy subjects

| Volume ($mm^3$) | Side | Brain region | BA | Peak voxel location (x, y, z) | | | ALE value ($10^{-2}$) |
|---|---|---|---|---|---|---|---|
| 56 | L | MI/AI | 13 | -40 | 10 | 0 | 1.56 |

Abbreviations: AI = anterior insula, ALE = Activation Likelihood Estimation, BA = Brodmann Area, L = left hemisphere, MI = middle insula, R = right hemisphere.

Voxel coordinates are in the MNI space.

Table S3

Brain regions activated by contrast analysis between anger provocation and core disgust in healthy subjects

| Volume ($mm^3$) | Side | Brain region | BA | Peak voxel location (x, y, z) | | | Z |
|---|---|---|---|---|---|---|---|
| *Anger provocation > Core disgust* | | | | | | | |
| 1048 | L | AI/MI | 13 | -34 | 12 | 10 | 3.04 |
| | L | AI/Claustrum | | -28 | 16 | 8 | 2.85 |
| 832 | R | AI/MI/Claustrum/Putamen | | 26 | 21 | 6 | 2.99 |
| 728 | L | Thalamus | | -17 | -19 | 5 | 3.35 |
| | L | Thalamus | | -14 | -20 | 12 | 3.16 |
| 712 | R | MCC | 24 | 6 | 6 | 44 | 3.54 |
| | R | MCC | 24 | 6.6 | 10.3 | 45.4 | 3.43 |
| | R | MCC | 24 | 9.3 | 10.7 | 36.7 | 3.29 |
| | R | MCC/dMPFC | 24/32 | 12 | 6 | 38 | 3.24 |

| | | | | | | | |
|---|---|---|---|---|---|---|---|
| *Core disgust > Anger provocation* | | | | | | | |
| 1424 | L | Amygdala/Parahippocampal Gyrus | | -22 | -2 | -24 | 2.82 |
| | L | Uncus | 34 | -16 | 0 | -26 | 2.70 |
| 1392 | L | Inferior Parietal Lobule/ Postcentral Gyrus | 40/2 | -56 | -22 | 34 | 2.79 |
| | L | Inferior Parietal Lobule/ Postcentral Gyrus | 40/2 | -56 | -26 | 34 | 2.78 |
| | L | Inferior Parietal Lobule/ Postcentral Gyrus | 40/2 | -58 | -26 | 28 | 2.44 |
| 968 | R | Amygdala/Parahippocampal Gyrus/Uncus | | 30 | 2 | -22 | 2.78 |
| | R | Amygdala/Parahippocampal Gyrus/Uncus/Globus Pallidus | | 22 | -4 | -26 | 2.37 |
| 664 | R | Inferior Occipital Gyrus | 17 | 12 | -96 | 0 | 2.47 |
| | R | Inferior Occipital Gyrus | 17 | 14 | -96 | -6 | 2.43 |
| | R | Inferior Occipital Gyrus | 17 | 16 | -96 | 0 | 2.37 |
| 384 | L | Fusiform Gyrus | 37 | -45 | -56 | -6 | 2.15 |
| | L | Fusiform Gyrus | 37 | -42 | -52 | -12 | 2.12 |
| 208 | R | Postcentral Gyrus/Inferior Parietal Lobule | 2/40 | 60 | -22 | 34 | 1.94 |
| | R | Postcentral Gyrus/Inferior Parietal Lobule | 2/40 | 64 | -18 | 34 | 1.73 |

Abbreviations: dMPFC = dorsomedial prefrontal cortex, MCC = middle cingulate cortex.
Voxel coordinates are in the MNI space.

Table S4
Conjunction analysis of MACM and RSFC results for bilateral anterior insula from Unfairness ∩ Core disgust

| Volume ($mm^3$) | Side | Brain region | BA | Peak voxel location (x, y, z) | | |
|---|---|---|---|---|---|---|
| 12552 | L | Insula/Inferior Frontal Gyrus/IFG Pars Orbitalis/Superior Temporal Gyrus/ Precentral Gyrus/Claustrum/Rolandic Operculum/Putamen/Middle Frontal Gyrus | 13/47/45/ 44/22/38 | -32 | 22 | 0 |
| 11664 | R | Insula/Inferior Frontal Gyrus/IFG Pars Orbitalis/Putamen/Precentral Gyrus/ Superior Temporal Gyrus/Claustrum/ Middle Frontal Gyrus | 13/47/45/ 44/22/38 | 36 | 24 | -2 |
| 8216 | R | Middle Cingulate/Anterior Cingulate/ Superior Frontal Gyrus/Medial Frontal Gyrus/Supplementary Motor Area | 32/24/9/6 | 6 | 20 | 40 |
| | L | Middle Cingulate/Anterior Cingulate/ Superior Frontal Gyrus/Medial Frontal | 32/24/9/6 | | | |

| | | | | | | |
|---|---|---|---|---|---|---|
| | | Gyrus/Supplementary Motor Area | | | | |
| 232 | R | Middle Frontal Gyrus | | 40 | 44 | 18 |

Voxel coordinates are in the MNI space.

Table S5

Conjunction analysis of MACM and RSFC results for bilateral anterior insula from Unfairness ∩ Anger provocation

| Volume (mm³) | Side | Brain region | BA | Peak voxel location (x, y, z) | | |
|---|---|---|---|---|---|---|
| 14520 | R | Insula/Inferior Frontal Gyrus/Putamen/ Precentral Gyrus/IFG Pars Orbitalis/ Rolandic Operculum/Claustrum/Superior Temporal Gyrus/Caudate/Middle Frontal Gyrus | 13/47/44/ 45/22/9 | 34 | 22 | 2 |
| 13232 | L | Insula/Inferior Frontal Gyrus/Precentral Gyrus/Superior Temporal Gyrus/Putamen/ IFG Pars Orbitalis/Rolandic Operculum/ Claustrum/Putamen | 13/47/22/ 44/45/9 | -32 | 20 | 4 |
| 9504 | R | Middle Cingulate/Anterior Cingulate/ Superior Frontal Gyrus/Medial Frontal Gyrus/Supplementary Motor Area | 32/24/9/6 /8 | 6 | 18 | 40 |
| | L | Middle Cingulate/Superior Frontal Gyrus/ Anterior Cingulate/Medial Frontal Gyrus/ Supplementary Motor Area | 32/24/9/6 /8 | | | |
| 888 | R | Middle Frontal Gyrus/Superior Frontal Gyrus | 10 | 40 | 42 | 18 |
| 568 | L | Middle Frontal Gyrus/Superior Frontal Gyrus | 10 | -34 | 44 | 24 |
| 560 | R | Superior Temporal Gyrus/Middle Temporal Gyrus | 22 | 58 | -26 | 6 |
| 328 | L | Insula/Superior Temporal Gyrus | | -52 | -40 | 20 |
| 160 | L | Putamen/Caudate | | -16 | 6 | 8 |

Voxel coordinates are in the MNI space.

Table S6

Conjunction analysis of MACM and RSFC results for left middle insula from Core disgust ∩ Anger provocation

| Volume (mm³) | Side | Brain region | BA | Peak voxel location (x, y, z) | | |
|---|---|---|---|---|---|---|
| 21568 | R | Insula/Inferior Frontal Gyrus/Superior Temporal Gyrus/Rolandic Operculum/ Putamen/Postcentral Gyrus/Precentral Gyrus/IFG Pars Orbitalis/Supramarginal Gyrus/Claustrum/Inferior Parietal Lobule/ | 13/47/22/ 44/41/45/ 40/43/38/ 42 | 42 | 10 | 2 |

| | | | | | | |
|---|---|---|---|---|---|---|
| | | Middle Temporal Gyrus/Pallidum | | | | |
| 18608 | L | Insula/Inferior Frontal Gyrus/Superior Temporal Gyrus/Precentral Gyrus/ Rolandic Operculum/Putamen/IFG Pars Orbitalis/Claustrum/Postcentral Gyrus/ Pallidum/Middle Temporal Gyrus | 13/47/22/ 44/38/45/ 6/43/42 | -40 | 10 | 0 |
| 2424 | L | Superior Temporal Gyrus/Postcentral Gyrus/Supramarginal Gyrus/Inferior Parietal Lobule/Insula/Middle Temporal Gyrus | 40/42/13 | -54 | -34 | 18 |

Voxel coordinates are in the MNI space.

Table S7

Conjunction analysis of MACM and RSFC results for bilateral anterior insula from Unfairness > Core disgust

| Volume ($mm^3$) | Side | Brain region | BA | Peak voxel location (x, y, z) | | |
|---|---|---|---|---|---|---|
| 7376 | L | Insula/Inferior Frontal Gyrus/IFG Pars Orbitalis/Claustrum/Superior Temporal Gyrus/Putamen | 13/47/45/ 22/38 | -34 | 18 | -8 |
| 6584 | R | Insula/Inferior Frontal Gyrus/IFG Pars Orbitalis/Putamen/Claustrum/Superior Temporal Gyrus | 47/13/45 | 36 | 20 | -6 |
| 336 | R | Inferior Frontal Gyrus | 44 | 54 | 12 | 6 |

Voxel coordinates are in the MNI space.

Table S8

Conjunction analysis of MACM and RSFC results for bilateral anterior insula from Unfairness > Anger provocation

| Volume ($mm^3$) | Side | Brain region | BA | Peak voxel location (x, y, z) | | |
|---|---|---|---|---|---|---|
| 8360 | L | Insula/Inferior Frontal Gyrus/IFG Pars Orbitalis/Superior Temporal Gyrus/ Claustrum/Putamen | 47/13/38/ 45/22 | -36 | 18 | -8 |
| 7400 | R | Insula/Inferior Frontal Gyrus/IFG Pars Orbitalis/Superior Temporal Gyrus/ Putamen | 47/13/38/ 45/22 | 38 | 20 | -8 |

Voxel coordinates are in the MNI space.

Table S9

Conjunction analysis of MACM and RSFC results for bilateral anterior insula from Anger provocation > Core disgust

| Volume ($mm^3$) | Side | Brain region | BA | Peak voxel location (x, y, z) |
|---|---|---|---|---|

| | | | | | | |
|---|---|---|---|---|---|---|
| 6040 | L | Middle Cingulate/Supplementary Motor Area/Medial Frontal Gyrus/Superior Frontal Gyrus/Anterior Cingulate | 32/24/6/8 | 8 | 16 | 40 |
| | R | Middle Cingulate/Supplementary Motor Area/Medial Frontal Gyrus/Superior Frontal Gyrus | 32/24/6/8 | | | |
| 4664 | L | Insula/Putamen/Inferior Frontal Gyrus/ Claustrum | 13/47/45 | -32 | 16 | 8 |
| 3608 | R | Insula/Inferior Frontal Gyrus/Putamen/ Claustrum | 13/47/45 | 32 | 20 | 6 |
| 1384 | L | Thalamus | | -14 | -16 | 8 |
| 1120 | R | Thalamus | | 12 | -12 | 6 |
| 464 | R | Inferior Frontal Gyrus | | 48 | 10 | 24 |
| 304 | L | Inferior Frontal Gyrus | | -46 | 6 | 28 |
| 120 | R | Putamen | | 24 | 8 | 6 |

Voxel coordinates are in the MNI space.

Table S10

Conjunction analysis of MACM and RSFC results for left anterior insula from Anger provocation > Unfairness

| Volume ($mm^3$) | Side | Brain region | BA | Peak voxel location (x, y, z) | | |
|---|---|---|---|---|---|---|
| 24928 | L | Insula/Inferior Frontal Gyrus/Precentral Gyrus/Middle Frontal Gyrus/Superior Temporal Gyrus/Rolandic Operculum/ Claustrum/IFG Pars Orbitalis/Putamen | 13/44/9/ 45/46/47/ 10/22/6 | -38 | 18 | 8 |
| 11320 | R | Insula/Inferior Frontal Gyrus/Precentral Gyrus/Putamen/Rolandic Operculum/ Claustrum/Superior Temporal Gyrus/ Middle Frontal Gyrus | 13/47/44/ 45/9/22 | 36 | 20 | 6 |
| 1284 | L | Inferior Parietal Lobule | | -34 | -48 | 44 |
| 952 | L | Thalamus | | -12 | -16 | 6 |
| 672 | R | Thalamus | | 12 | -16 | 6 |

Voxel coordinates are in the MNI space.

Table S11

Neurotransmitter receptors and transporters included in analyses

| Receptor/ transporter | Neurotransmitter | Tracer | Measure | N | Age | Primary reference |
|---|---|---|---|---|---|---|
| 5HT1a | Serotonin | [11C]WAY-100635 | BPnd | 35 | 26.3±5.2 | (Savli et al., 2012) |
| 5HT1b | Serotonin | [11C]P943 | BPnd | 23 | 28.7±7 | (Savli et al., 2012) |

| | | | | | | |
|---|---|---|---|---|---|---|
| 5HT2a | Serotonin | [18F]ALTANSERIN | BPnd | 19 | 28.2±5.7 | (Savli et al., 2012) |
| 5HT4 | Serotonin | [11C]SB207145 | Bmax | 59 | 25.9±5.3 | (Beliveau et al., 2017) |
| 5HT6 | Serotonin | [11C]GSK215083 | BPnd | 30 | 36.6±9.0 | (Radhakrishnan et al., 2018) |
| 5HTT | Serotonin | [11C]DASB | BPnd | 18 | 30.5±9.5 | (Savli et al., 2012) |

Abbreviations: BPnd, non-displaceable binding potential; Bmax, density (pmol $ml^{-1}$) converted from binding potential (5-HT) using autoradiography-derived densities.

Table S12

Brain regions activated by rejection in healthy subjects (cluster-FWE p<0.05, voxel-level p<0.001 (uncorrected), 5,000 permutations)

| Volume ($mm^3$) | Side | Brain region | BA | Peak voxel location (x, y, z) | | | ALE value ($10^{-2}$) | Z |
|---|---|---|---|---|---|---|---|---|
| 936 | L | Putamen | | -22 | 10 | 0 | 2.50 | 5.67 |

Voxel coordinates are in the MNI space.

Table S13

Brain regions activated by contrast analysis between rejection and core disgust in healthy subjects

| Volume ($mm^3$) | Side | Brain region | BA | Peak voxel location (x, y, z) | | | Z |
|---|---|---|---|---|---|---|---|
| *Rejection > Core disgust* | | | | | | | |
| 936 | L | Putamen | | -26 | 14 | 0 | 3.43 |
| | | | | -24 | 7 | 3 | 3.09 |
| | | | | -22 | 12 | 0 | 3.04 |
| *Core disgust > Rejection* | | | | | | | |
| 960 | R | Lingual gyrus | 18 | 12.7 | -86.7 | -0.3 | 3.72 |
| | R | Inferior Occipital Gyrus | 17 | 14 | -94 | -6 | 3.29 |
| | R | Lingual Gyrus | 17 | 18 | -93 | 0 | 3.19 |
| | R | Lingual Gyrus | 18 | 18 | -94 | -8 | 3.06 |
| | R | Inferior Occipital Gyrus | 17 | 20 | -94 | -4 | 2.99 |
| 680 | L | Inferior Occipital Gyrus | 19 | -42 | -80 | 4 | 1.85 |
| | L | Inferior Temporal Gyrus | 19 | -46 | -81.3 | 5.3 | 1.85 |
| | L | Middle Occipital Gyrus | 37 | -44 | -74 | 4 | 1.76 |
| 288 | L | Inferior Parietal Lobule/ Postcentral Gyrus | 40/2 | -57.3 | -21.3 | 36 | 1.98 |

Voxel coordinates are in the MNI space. Note that both ALE maps used for contrast analysis were based on the following standard threshold: cluster-FWE p<0.05, voxel-level p<0.001 (uncorrected), 5,000 permutations.

Table S14

Conjunction analysis of MACM and RSFC results for left putamen from the rejection network

| Volume ($mm^3$) | Side | Brain region | BA | Peak voxel location (x, y, z) | | |
|---|---|---|---|---|---|---|
| 15984 | L | Putamen/Thalamus/Insula/Caudate/ Pallidum/Globus Pallidus/Claustrum/ Inferior Frontal Gyrus | | -20 | 8 | 0 |
| 12336 | R | Putamen/Thalamus/Caudate/Insula/ Pallidum/Inferior Frontal Gyrus/Globus Pallidus/Claustrum | | 24 | 8 | 0 |
| 11160 | L | Supplementary Motor Area/Middle Cingulate/Medial Frontal Gyrus/Superior Frontal Gyrus/Anterior Cingulate | 32/6/24/8 /9 | -2 | 10 | 48 |
| | R | Middle Cingulate/Medial Frontal Gyrus/ Supplementary Motor Area/Superior Frontal Gyrus/Anterior Cingulate | | | | |
| 912 | L | Inferior Frontal Gyrus/ Precentral Gyrus | 44 | -52 | 10 | 16 |

Voxel coordinates are in the MNI space.

Table S15

Brain regions activated by rejection in healthy subjects (voxel-level p<0.001, uncorrected)

| Volume ($mm^3$) | Side | Brain region | BA | Peak voxel location (x, y, z) | | | ALE value ($10^{-2}$) | Z |
|---|---|---|---|---|---|---|---|---|
| 936 | L | Putamen | | -22 | 10 | 0 | 2.50 | 5.67 |
| 344 | R | MI/AI/Claustrum | 13 | 38 | 0 | 10 | 1.66 | 4.38 |
| 312 | L | Postcentral Gyrus/Inferior Parietal Lobule/Superior Temporal Gyrus/MI | 40 | -56 | -26 | 20 | 1.47 | 4.01 |
| 264 | L | MCC | 24 | -8 | 6 | 42 | 1.07 | 3.33 |
| 256 | L | MI/AI/Claustrum | 13 | -32 | 4 | 12 | 1.57 | 4.21 |
| 256 | R | Cuneus | 18 | 20 | -78 | 30 | 1.42 | 3.93 |
| 208 | R | Postcentral Gyrus/Inferior Parietal Lobule | 40 | 58 | -22 | 22 | 1.22 | 3.59 |
| 200 | L | Parahippocampal Gyrus/Amygdala | 34 | -22 | 0 | -20 | 1.28 | 3.69 |
| 168 | L | ACC | 32 | -4 | 42 | 14 | 1.26 | 3.66 |
| 160 | L | Culmen | | -22 | -54 | -26 | 1.15 | 3.48 |

Abbreviation: ACC = anterior cingulate cortex. Voxel coordinates are in the MNI space.

Table S16

Brain regions activated by conjunction analysis between rejection and core disgust in healthy subjects

| Volume (mm$^3$) | Side | Brain region | BA | Peak voxel location (x, y, z) | | | ALE value ($10^{-2}$) |
|---|---|---|---|---|---|---|---|
| 200 | L | Amygdala/Parahippocampal Gyrus | 34 | -22 | 0 | -20 | 1.28 |

Voxel coordinates are in the MNI space. Note that the ALE map of rejection used for conjunction analysis was based on the following threshold: voxel-level p<0.001, uncorrected; while the ALE map of core disgust used for conjunction analysis was based on the following threshold: cluster-FWE p<0.05, voxel-level p<0.001 (uncorrected), 5,000 permutations.

Table S17

Brain regions activated by contrast analysis between rejection and core disgust in healthy subjects

| Volume (mm$^3$) | Side | Brain region | BA | Peak voxel location (x, y, z) | | | Z |
|---|---|---|---|---|---|---|---|
| *Rejection* > *C*ore disgust | | | | | | | |
| 936 | L | Putamen | | -26 | 13 | -1 | 3.72 |
| | L | Putamen | | -24 | 7 | 3 | 3.19 |
| 328 | R | MI/AI | 13 | 42 | 2 | 10 | 2.10 |
| | R | Claustrum | | 36 | 2 | 10 | 2.10 |
| 256 | L | MI/AI/Claustrum | 13 | -34 | 8 | 12 | 2.42 |
| *C*ore disgust > *Rejection* | | | | | | | |
| 960 | R | Lingual Gyrus | 18 | 13.2 | -87.2 | -1.7 | 3.89 |
| | R | Inferior Occipital Gyrus | 17 | 12 | -94 | -2 | 3.72 |
| | R | Lingual Gyrus | 17 | 18 | -93 | 0 | 3.43 |
| | R | Inferior Occipital Gyrus | 17 | 14 | -94 | -6 | 3.35 |
| | R | Lingual Gyrus | 18 | 18 | -94 | -8 | 3.09 |
| | R | Inferior Occipital Gyrus | 17 | 20 | -94 | -4 | 2.97 |
| 672 | L | Inferior Occipital Gyrus | 19 | -42 | -80 | 4 | 1.87 |
| | L | Inferior Temporal Gyrus | 19 | -46 | -81.3 | 5.3 | 1.86 |
| | L | Inferior Temporal Gyrus | | -48 | -74 | 6 | 1.85 |
| | L | Inferior Temporal Gyrus | | -50 | -74 | 2 | 1.85 |
| | L | Middle Occipital Gyrus | 37 | -44 | -74 | 4 | 1.74 |
| 280 | L | Inferior Parietal Lobule/ Postcentral Gyrus | 40/2 | -57.3 | -21.3 | 36 | 1.98 |

Voxel coordinates are in the MNI space. Note that the ALE map of rejection used for contrast analysis was based on the following threshold: voxel-level p<0.001, uncorrected; while the ALE map of core disgust used for contrast analysis was based on the following threshold: cluster-FWE p<0.05, voxel-level p<0.001 (uncorrected), 5,000 permutations.

Table S18

Brain regions activated by contrast analysis between rejection and anger provocation in healthy subjects

| Volume ($mm^3$) | Side | Brain region | BA | Peak voxel location (x, y, z) | | | Z |
|---|---|---|---|---|---|---|---|
| *Rejection* > Anger provocation | | | | | | | |
| 856 | L | Putamen | | -20 | 14 | -2 | 2.54 |
| | L | Putamen | | -24 | 8 | 4 | 2.37 |
| Anger provocation > *Rejection* | | | | | | | |
| 208 | R | AI/MI | 13 | 34 | 24 | 2 | 1.91 |
| | R | Claustrum | | 28 | 26 | 2 | 1.87 |

Voxel coordinates are in the MNI space. Note that the ALE map of rejection used for contrast analysis was based on the following threshold: voxel-level p<0.001, uncorrected; while the ALE map of core disgust used for contrast analysis was based on the following threshold: cluster-FWE p<0.05, voxel-level p<0.001 (uncorrected), 5,000 permutations.

------------------------ Insert Figure S1 about here ----------------------------

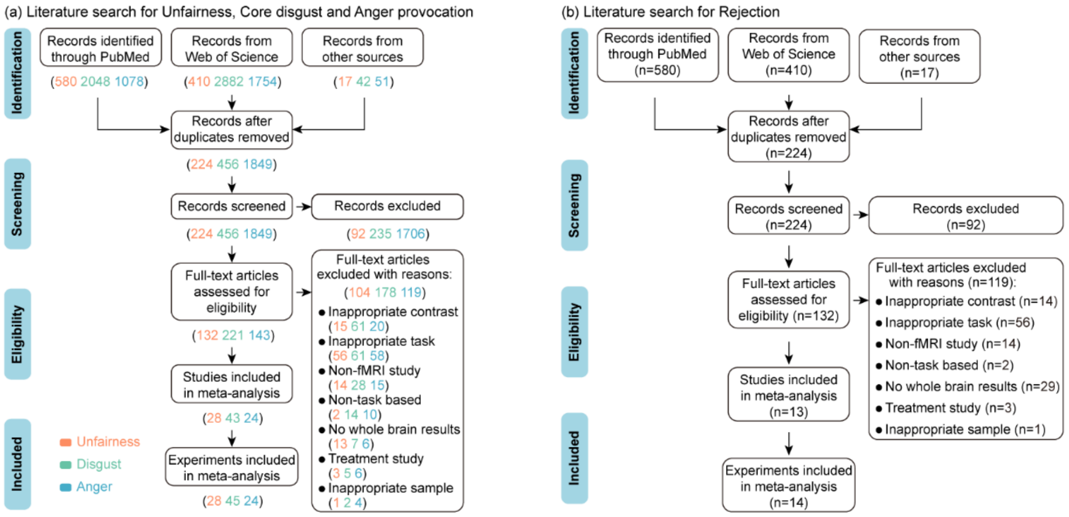

Figure S1. Flowchart illustrating the article screening, exclusion, and inclusion process according to the PRISMA procedure. (a) for Unfairness, Core disgust, and Anger provocation. (b) for Rejection.

------------------------ Insert Figure S2 about here ----------------------------

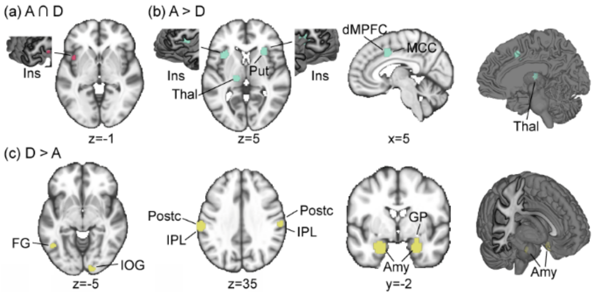

Figure S2. Overview of significant brain regions from the conjunction and contrast analyses between anger provocation and core disgust. (a) Common activations for anger provocation and core disgust in healthy subjects. (b) Greater activations for anger provocation than for core disgust (anger provocation > core disgust) in healthy subjects. (c) Greater activations for core disgust than for anger provocation (core disgust > anger provocation) in healthy subjects. Coordinates are in the MNI space. Abbreviations: A = anger provocation, Amy = Amygdala, D = core disgust, dMPFC = dorsomedial prefrontal cortex, FG = Fusiform Gyrus, GP = Globus Pallidus, Ins = Insula, IOG = Inferior Occipital Gyrus, IPL = Inferior Parietal Lobule, MCC = middle cingulate cortex, Postc = Postcentral Gyrus, Put = Putamen, Thal = Thalamus.

------------------------ Insert Figure S3 about here ----------------------------

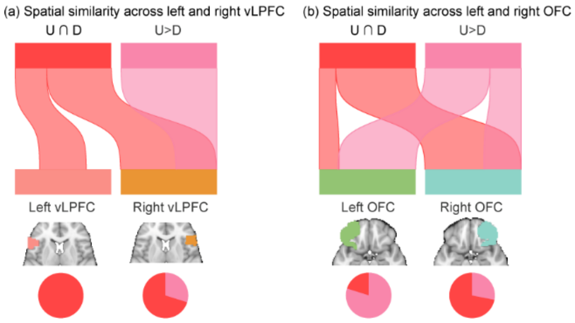

Figure S3. (a) River plots showing spatial similarity (cosine similarity) between bilateral vLPFC and maps of unfairness ∩ core disgust as well as unfairness > core disgust. (b) River plots showing spatial similarity (cosine similarity) between bilateral OFC and maps of unfairness ∩ core disgust as well as unfairness > core disgust. Ribbons are normalized by the max cosine similarity across left and right vLPFC (a) or OFC (b). Ribbon locations in relation to the boxes are arbitrary. Pie charts show relative contributions of each map to each ROI (i.e., percentage of voxels with the highest cosine similarity for each map). Abbreviations: OFC = orbital frontal cortex, U = unfairness, vLPFC = ventrolateral prefrontal cortex.

------------------------ Insert Figure S4 about here ----------------------------

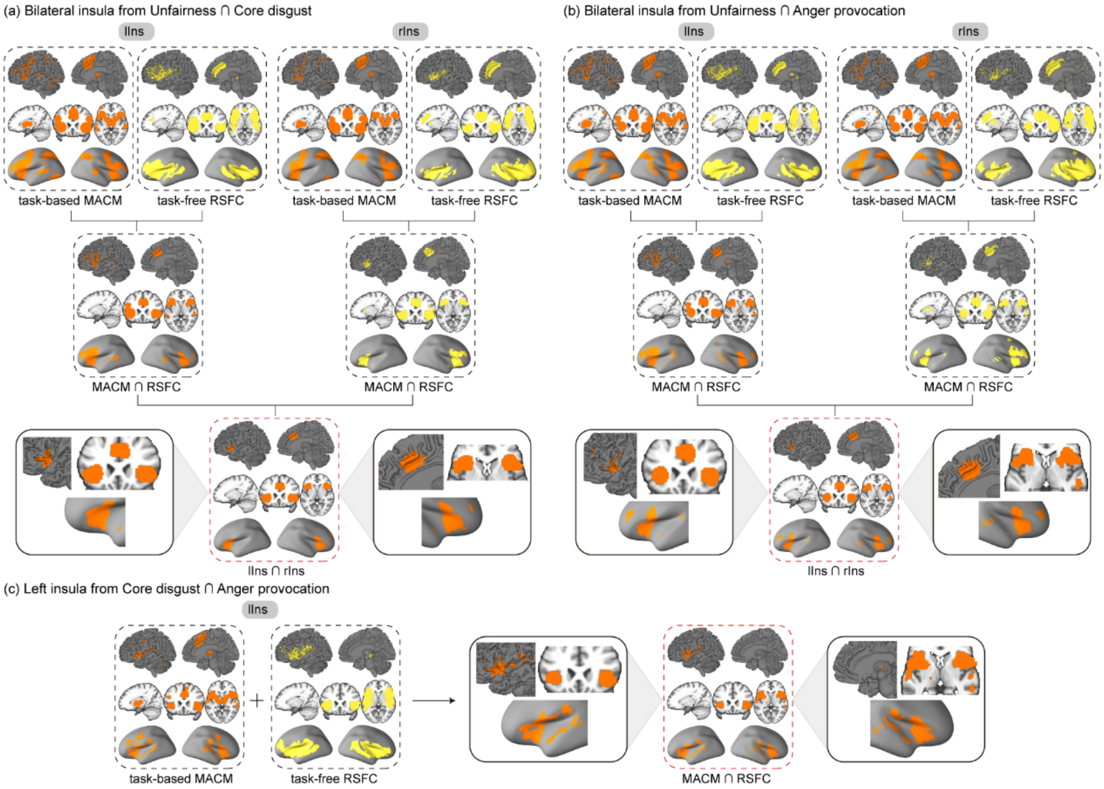

Figure S4. The detailed methodological workflows of consensus connectivity networks for the insula from conjunction analyses. (a) Results of consensus connectivity map for bilateral AI from the conjunction of unfairness and core disgust. (b) Results of consensus connectivity map for bilateral AI from the conjunction of unfairness and anger provocation. (c) Results of consensus connectivity map for left MI from the conjunction of core disgust and anger provocation. Abbreviations: lIns = left insula, MACM = meta-analytic connectivity modeling, rIns = right insula, RSFC = resting-state functional connectivity.

------------------------ Insert Figure S5 about here ----------------------------

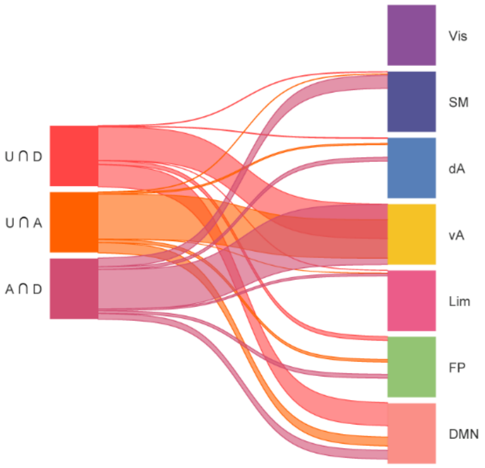

Figure S5. Network assignment of the consensus connectivity maps for the insula from conjunction analyses using river plots. Abbreviations: dA = dorsal attention network, DMN = default mode network, FP = frontoparietal network, Limb = limbic network, SM = somatomotor network, vA = ventral attention network, Vis = visual network.

------------------------ Insert Figure S6 about here ----------------------------

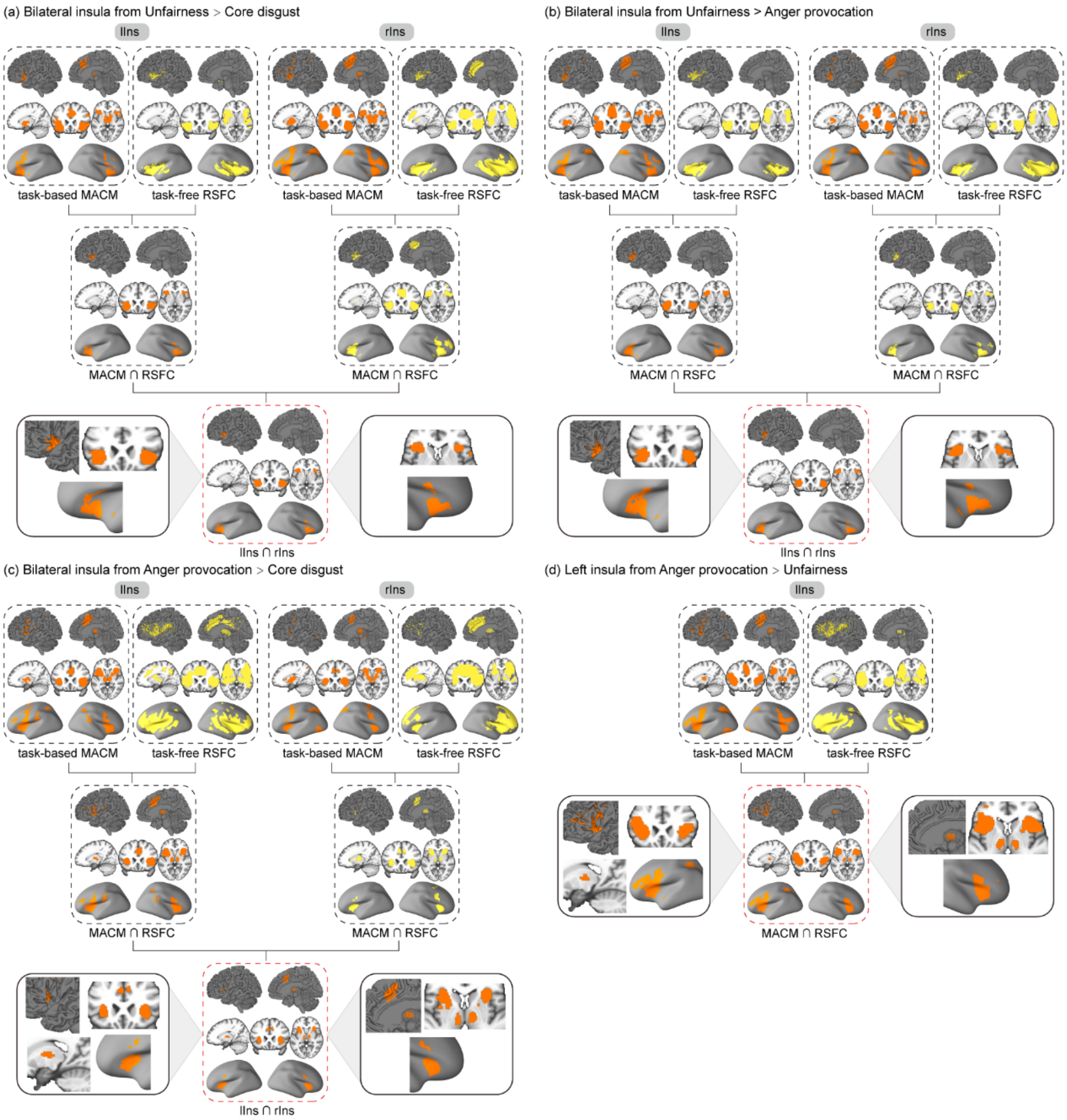

Figure S6. The detailed methodological workflows of consensus connectivity networks for the insula from contrast analyses. (a) Results of consensus connectivity map for bilateral AI from the contrast of unfairness > core disgust. (b) Results of consensus connectivity map for bilateral AI from the contrast of unfairness > anger provocation. (c) Results of consensus connectivity map for bilateral AI from the contrast of anger provocation > core disgust. (d) Results of consensus connectivity map for left AI from the contrast of anger provocation > unfairness.

------------------------ Insert Figure S7 about here ----------------------------

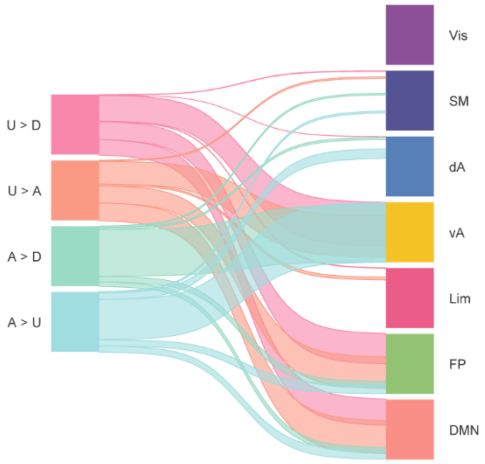

Figure S7. Network assignment of the consensus connectivity maps for the insula from contrast analyses using river plots.

------------------------ Insert Figure S8 about here ----------------------------

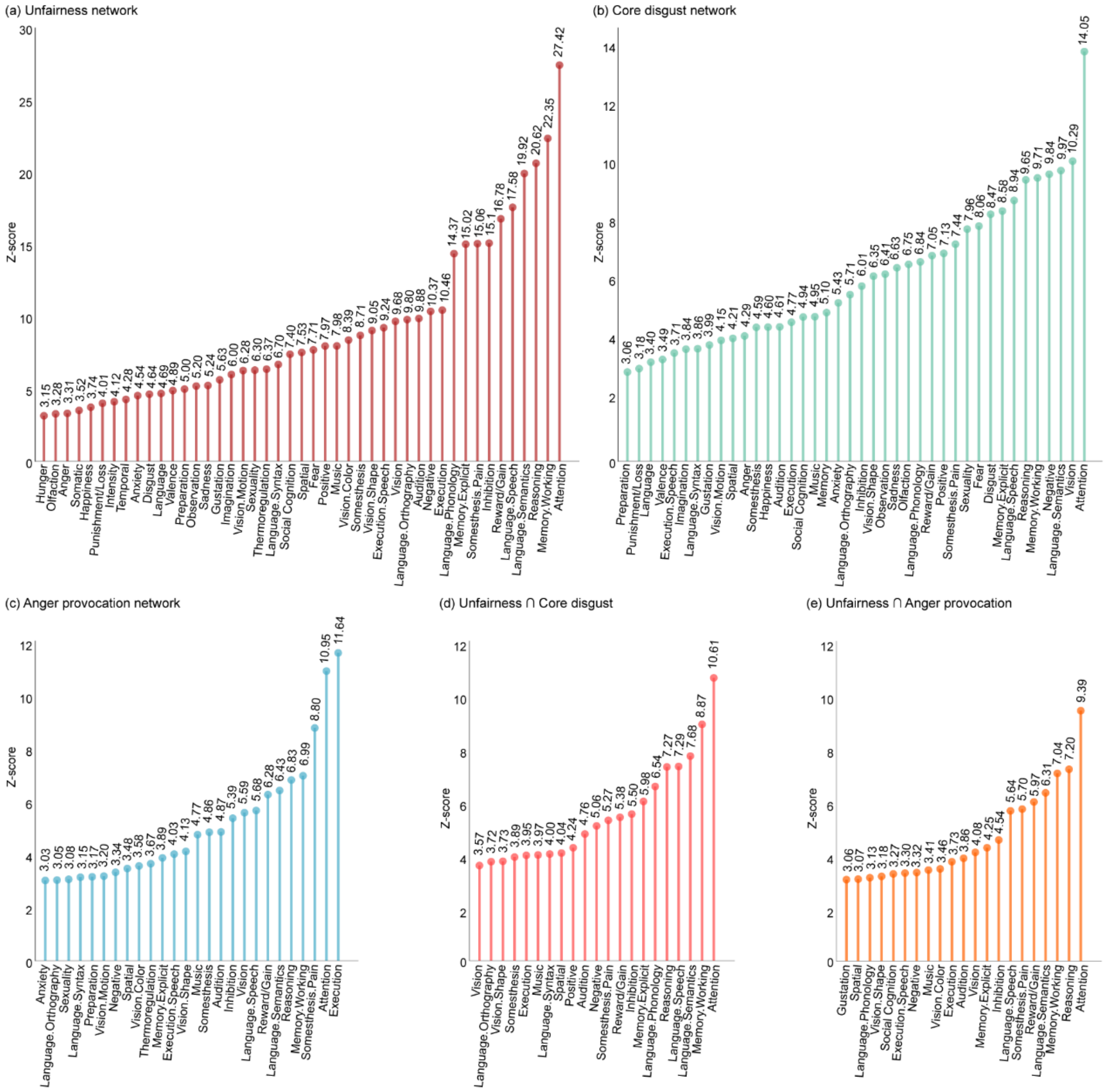

Figure S8. BrainMap-based behavioral characterization results of whole brain ALE maps from single dataset analyses and conjunction analyses. (a) Behavioral characterization of the unfairness network from single dataset analysis. (b) Behavioral characterization of the core disgust network from single dataset analysis. (c) Behavioral characterization of the anger provocation network from single dataset analysis. (d) Behavioral characterization of unfairness ∩ core disgust from conjunction analysis. (e) Behavioral characterization of unfairness ∩ anger provocation from conjunction analysis.

------------------------ Insert Figure S9 about here ----------------------------

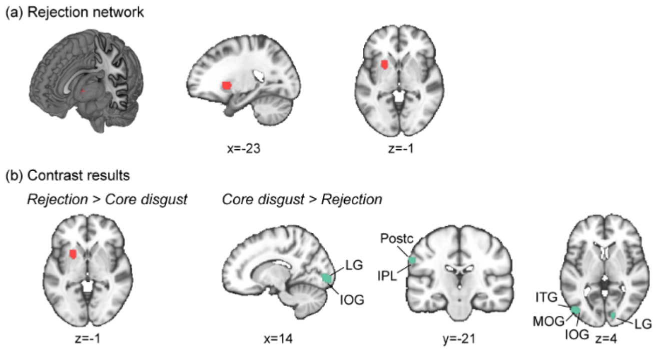

Figure S9. Overview of significant brain regions for the rejection domain. (a) Rejection network from single dataset analysis in healthy subjects (standard thresholding: cluster-FWE $p<0.05$, voxel-level $p<0.001$ (uncorrected), 5,000 permutations). (b) Contrast analyses between rejection and core disgust in healthy subjects. Note that both the ALE maps of rejection and core disgust used for contrast analyses were based on the following standard threshold: cluster-FWE $p<0.05$, voxel-level $p<0.001$ (uncorrected), 5,000 permutations. Abbreviations: ITG = Inferior Temporal Gyrus, LG = Lingual Gyrus, MOG = Middle Occipital Gyrus.

------------------------ Insert Figure S10 about here ----------------------------

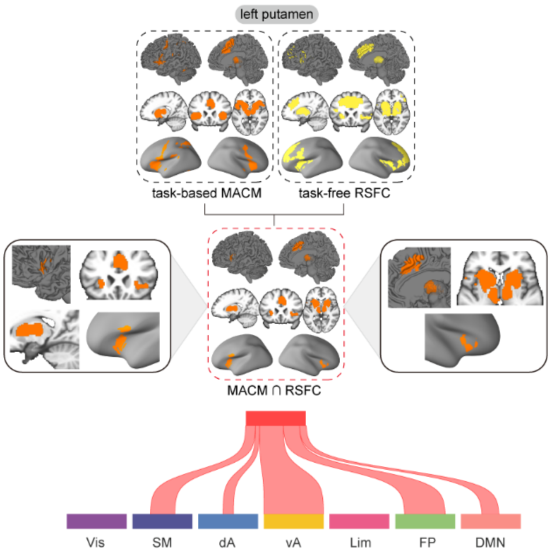

Figure S10. The detailed methodological workflow of consensus connectivity network for the left putamen (upper panel) and network assignment of the consensus connectivity map for the left putamen using river plots (lower panel).

------------------------ Insert Figure S11 about here ----------------------------

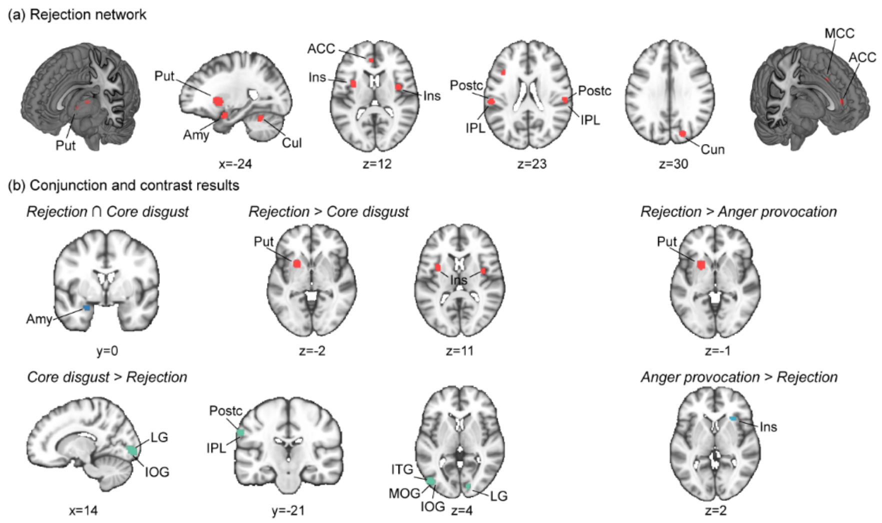

Figure S11. Overview of significant brain regions for the rejection domain. (a) Rejection network from single dataset analysis in healthy subjects (lenient thresholding: voxel-level p<0.001, uncorrected). (b) Conjunction and contrast analyses between rejection and core disgust in healthy subjects (left panel), contrast analyses between rejection and anger provocation (right panel). Note that the ALE map of rejection used for contrast and conjunction analyses was based on the following lenient threshold: voxel-level p<0.001, uncorrected; while the ALE map of core disgust (or anger provocation) used for contrast and conjunction analyses was based on the following standard threshold: cluster-FWE p<0.05, voxel-level p<0.001 (uncorrected), 5,000 permutations. Abbreviations: Cul = Culmen, Cun = Cuneus.
